# Non-modular Fatty Acid Synthases Yield Unique Acylation in Ribosomal Peptides

**DOI:** 10.1101/2023.10.25.564083

**Authors:** Hengqian Ren, Chunshuai Huang, Yuwei Pan, Haiyang Cui, Shravan R. Dommaraju, Douglas A. Mitchell, Huimin Zhao

## Abstract

Recent efforts in genome mining of ribosomally synthesized and post-translationally modified peptides (RiPPs) have expanded the diversity of post-translational modification chemistries^1, 2^. However, RiPPs are rarely reported as hybrid molecules incorporating biosynthetic machineries from other natural product families^3–8^. Here, we report lipoavitides, a class of RiPP/fatty acid hybrid lipopeptides that display a unique, membrane-targeting 4-hydroxy-2,4-dimethylpentanoyl (HMP)-modified *N*-terminus. The HMP is formed via condensation of isobutyryl-CoA and methylmalonyl-CoA catalyzed by a 3-ketoacyl-ACP synthase III enzyme, followed by successive tailoring reactions in the fatty acid biosynthetic pathway. The HMP and RiPP substructures are then connected by an acyltransferase exhibiting promiscuous activity towards the fatty acyl and RiPP substrates. Overall, the discovery of lipoavitides contributes a prototype of RiPP/fatty acid hybrids and provides possible enzymatic tools for lipopeptide bioengineering.

---

Recent advances in genome sequencing, bioinformatics, synthetic biology, and analytical chemistry have enabled the genome mining of ribosomally synthesized and post-translationally modified peptides (RiPPs)^1, 2^. RiPP biosynthesis starts with the expression of genetically encoded precursor peptides, followed by an organized post-translational modification (PTM) process. While only starting from 20 proteinogenic amino acids, RiPPs reveal vast structural diversity and hence a broad range of biological functions endowed by the disparate PTMs^9, 10^. To date, 49 class-defining PTMs have been identified^2^, most of which are derived from the precursor peptide backbone and side chains. For example, the oxazol(in)e and thiazol(in)e heterocycles, which are found in the linear azol(in)e-containing peptide (LAP)^11^, bottromycin^12^, thiopeptide^13^, and cyanobactin^14^ RiPP classes, are formed through multistep reactions between the hydroxyl or thiol group of serine, threonine, or cysteine and the preceding backbone carbonyl^15^. RiPPs can also be modified with structures that are not produced by their biosynthetic gene clusters (BGCs), as exemplified by glycocins, which are synthesized using host endogenous NDP-monosaccharides as glycosyl donors^16^. In some cases, modifying enzymes from different RiPP classes are also found in a single BGC to produce RiPPs with combined class-defining PTMs^17^. However, RiPP BGCs comprising non-RiPP biosynthetic genes/pathways for synthesizing unconventional PTMs are rarely reported.

Hybrid BGCs, encoding biosynthetic machineries from multiple natural product families, are widely distributed in nature, often resulting in products with merged structural features^18–20^. Notable examples include polyketide synthase–nonribosomal peptide synthetase hybrid BGCs, in which modules of type I polyketide synthases and nonribosomal peptide synthetases work together to produce compounds sharing structural features from both families^21^. Only a few RiPPs, including the *e*-series thiopeptides^3^, pheganomycins^4^, cacaoidin^5^, lipolanthines^6, 7^, and goadvionins^8^, are known to be synthesized by hybrid machinery (Scheme 1). To synthesize the non-RiPP-derived components of these molecules, the corresponding BGCs encode enzymes of other natural product families, such as the polyketide synthases in the goadvionin BGC. Investigation into these hybrid BGCs has unveiled new biosynthetic modes for the synthesis and fusion of RiPP and non-RiPP parts, providing potential routes for peptide natural product bioengineering.

**Scheme 1.**
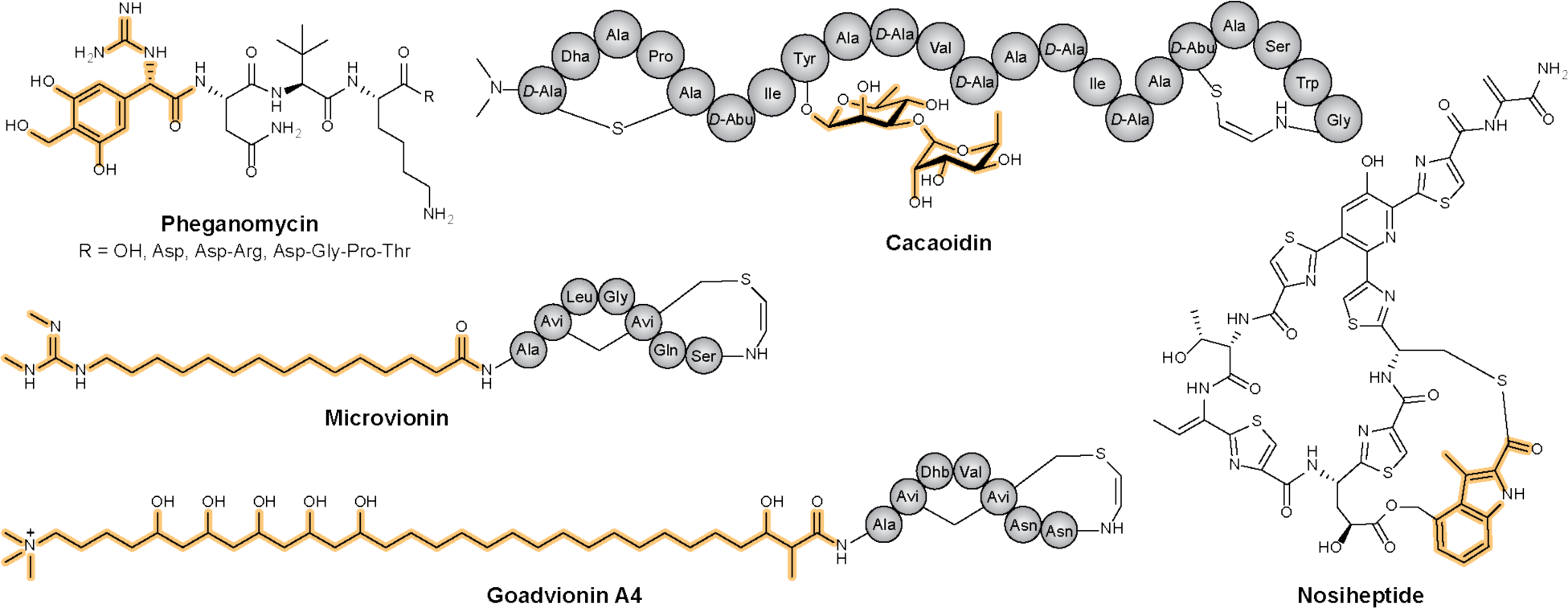
Representative RiPPs produced by RiPP/non-RiPP hybrid BGCs. Moieties synthesized by the non-RiPP pathways are highlighted in yellow. Microvionin, goadvionin A4, and nosiheptide are listed as class representatives for lipolanthins, goadvionins, and *e*-series thiopeptides, respectively.

In this work, we uncovered a class of hybrid BGCs that produce ribosomally derived lipopeptides. In addition to RiPP biosynthetic enzymes, these BGCs ubiquitously encode a 3-ketoacyl-ACP synthase III (FabH) and other enzymes resembling type II fatty acid biosynthesis. Direct cloning and heterologous expression of a representative BGC from *Streptomyces* sp. NRRL S-1521 yielded a 2-aminovinyl-(3-methyl)-cysteine (AviMeCys)-containing peptide with a 4-hydroxy-2,4-dimethylpentanoyl (HMP) modified *N*-terminus. This natural product thus represents the founding member of a new RiPP class termed the lipoavitides (AviMeCys-containing lipopeptides). The isolated lipoavitide exhibited hemolytic activity, and the subsequent structure-activity relationship study suggested an essential role of the *N*-terminal fatty acyl moiety. The HMP moiety is synthesized through condensation of isobutyryl-CoA and methylmalonyl-CoA by the FabH, followed by successive oxidation and reduction reactions, yielding HMP-CoA. An acyltransferase then utilizes HMP-CoA as the acyl donor and appends HMP to the RiPP portion of lipoavitide. Characterization of the acyltransferase showed a broad substrate tolerance which implies future potential in lipopeptide engineering. Overall, the discovery of lipoavitide serves as a prototype for RiPP/fatty acid hybrids.

## Discovery of RiPP BGCs encoding enzymes for fatty acid biosynthesis

During the genome mining of *Streptomyces* sp. NRRL S-1521 by AntiSMASH^22^, we identified a putative BGC (*lpv*) consisting of genes for synthesizing multiple families of natural products (Fig. 1, Supplementary Table 1, Supplementary Table 2). Encoded by *lpvBCD* are a HopA1-phosphotransferase pair and a flavin-dependent decarboxylase, a set of enzymes commonly known for synthesizing aminovinyl(methyl)-cysteine [Avi(Me)Cys] moieties^23^ in RiPPs, exemplified by the thioamitides^24^ and class V lanthipeptides^5, 25, 26^. A plethora of other known RiPP biosynthetic enzymes, such as luciferase-like monooxygenase (LpvJ), methyltransferase (LpvM1), and metalloproteases (LpvP1 and LpvP2), are also encoded in *lpv* (Supplementary Fig. 1, Supplementary Fig. 2). The *lpv* pathway further encodes a putative NDP-monosaccharide pathway (LpvLNOQM2) and a glycosyltransferase LpvI, suggesting the BGC produces a glycosylated RiPP(s). Of note, several biosynthetic enzymes (LpvFGHVS) in *lpv* are predicted to be involved in fatty acid/polyketide cyclization, elongation, keto/enoyl reduction, and oxidation^27^ (Supplementary Table 3, Supplementary Fig. 3, Supplementary Fig. 4). These enzymes are uncommon in RiPP BGCs and might produce an acyl group for usage by the acyltransferase LpvE.

**Fig. 1.**
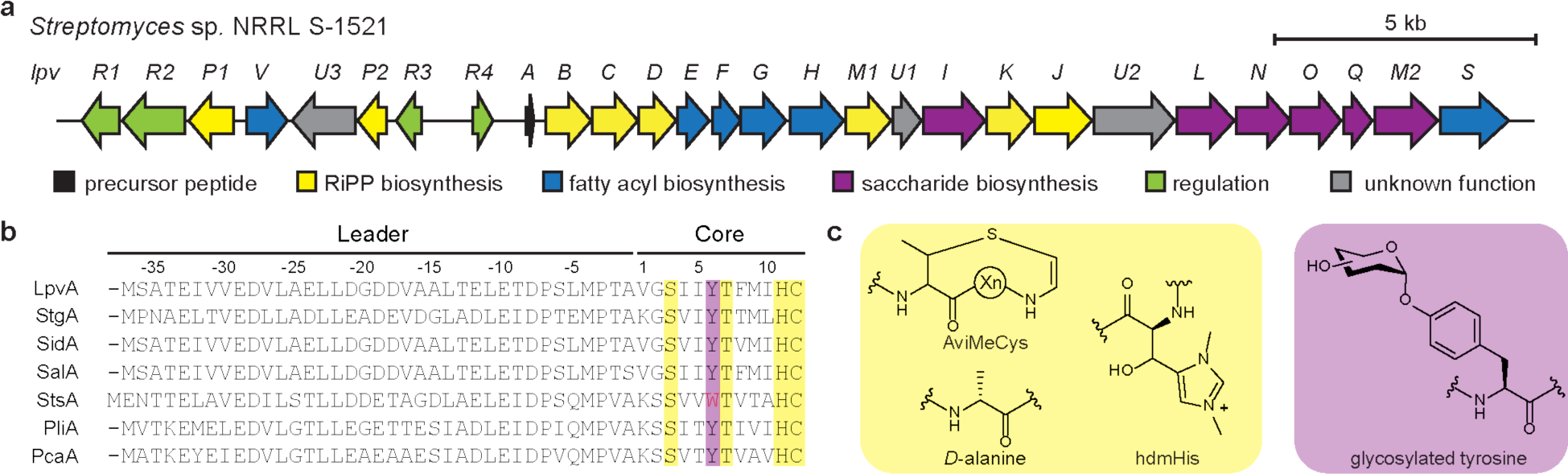
The lpv BGC identified in Streptomyces sp. NRRL S-1521. **a,** Gene organization of the *lpv* BGC. **b**, Alignment of LpvA with other putative precursor peptides from *lpv* homologous BGCs. Residues predicted to be modified by RiPP and saccharide biosynthetic enzymes are highlighted in yellow and purple, respectively. The tryptophan residue of StsA at the putative site of glycosylation is highlighted in red. **c**, Proposed PTMs in the final product.

To explore the distribution of *lpv* BGCs, we performed PSI-BLAST searches of the NCBI non-redundant database using LpvB (phosphotransferase), LpvC (HopA1), LpvE (acyltransferase), and LpvG (FabH) as queries, yielding 15 BGCs that have local homologs of all four enzymes and encompass seven unique precursor peptide sequences (Fig. 1, Supplementary Fig. 5). Alignment of these precursor peptides manifests a conserved TxxxHC motif, commonly found in thioviridamide-like thioamitides (Supplementary Table 4). We therefore hypothesized that the threonine and cysteine would form the AviMeCys moiety, and the histidine would be transformed into β-hydroxy-*N,N*-dimethyl-*L*-histidine (hdmHis) through methylation and hydroxylation by LpvM1 and LpvK, respectively^28, 29^ (Supplementary Fig. 1). Sequence alignment also identified a conserved serine residue, which is predicted to be converted into dehydroalanine by LpvB and LpvC, and subsequently reduced to *D*-alanine by the luciferase-like monooxygenase LpvJ^25^. Despite being absent in one of the seven precursor peptides, the tyrosine next to the TxxxHC motif is a likely glycosylation site, as observed in cacaoidin^5^ (Scheme 1). The replacement of tyrosine by tryptophan in the precursor peptide StsA concurred with the absence of the NDP-monosaccharide pathway and glycosyltransferase in the corresponding BGC, which supports our hypothesis (Fig. 1, Supplementary Fig. 5). Through bioinformatics analysis, we tentatively assigned functions to many of the encoded genes; however, no indications were found for residue(s) that may undergo acylation. To investigate the function of the acyltransferase and enzymes for fatty acid/polyketide synthesis, we next experimentally characterized the *lpv* BGC.

## Discovery of lipoavitide with an acylated *N*-terminus

To being our functional investigation, we used the CAPTURE (Cas12a-assisted precise targeted cloning using in vivo Cre-*lox* recombination) method to directly clone *lpv* for heterologous expression^30^. The cloned *lpv* and the pBE45 empty vector were individually conjugated into *Streptomyces lividans* TK24 and *Streptomyces albus* J1074. After 5 d of cultivation, colonies were picked, extracted by methanol, and analyzed by matrix-assisted laser desorption/ionization-time-of-flight mass spectrometry (MALDI-TOF-MS). A prominent ion at *m/z* 1636 was only observed from the *lpv*-expressing *S. albus* J1074 (Fig. 2, Supplementary Fig. 6). A 10 L production culture was then prepared to afford ∼10 mg of material for structural characterization (Supplementary Fig. 7).

**Fig. 2.**
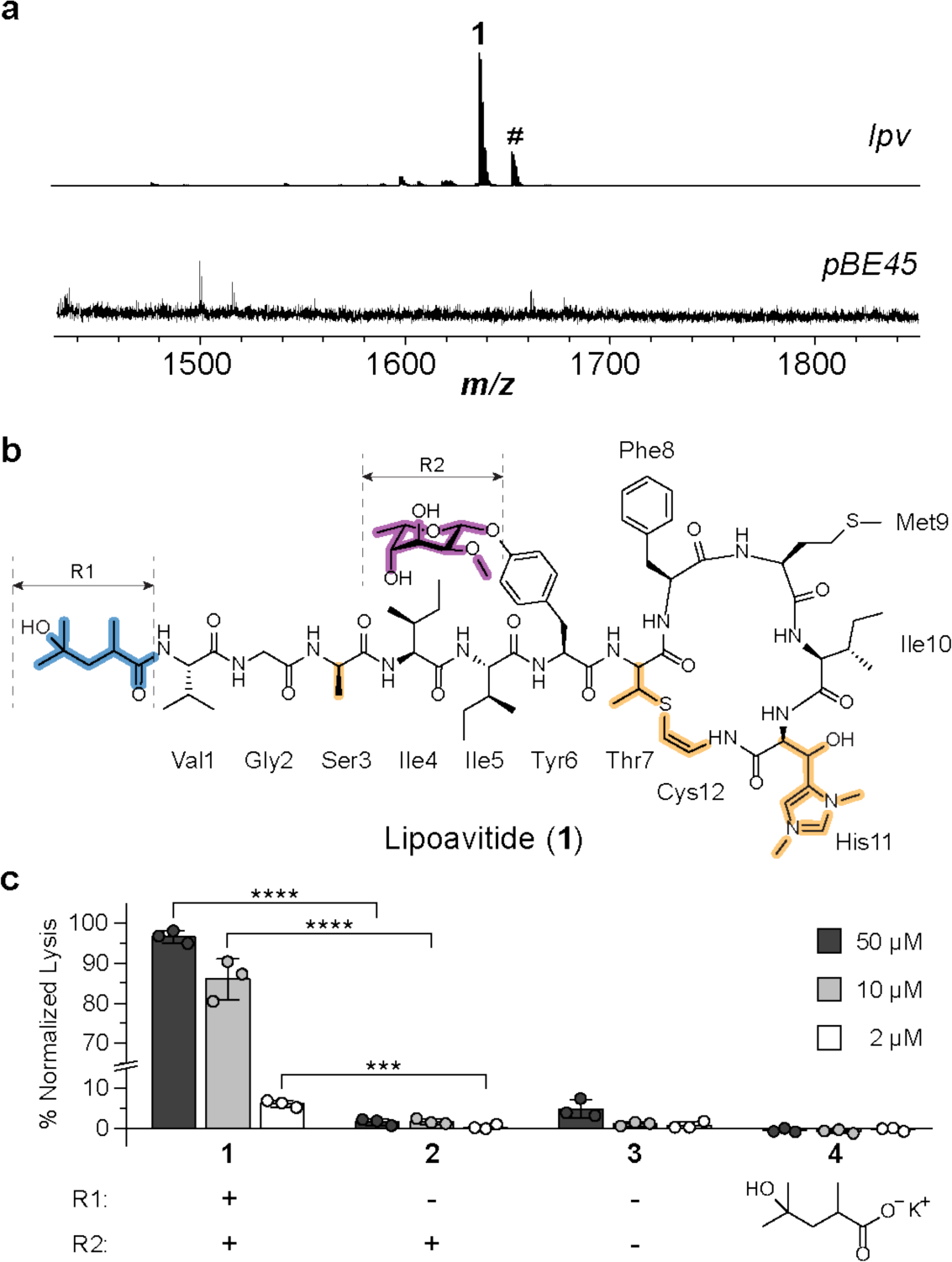
Heterologous expression and product characterization of lpv. **a**, MALDI-TOF mass spectra of *S. albus* J1074 containing *lpv* (**1**, *m/z* 1636) and the pBE45 empty vector. The signal denoted by ‘#’ (*m/z* 1652) suggests partial oxidation of the methionine sulfur in **1**, leading to a gain of 16 Da. **b**, Proposed structure of lipoavitide **1**. NMR spectra of **1** and assigned signals are shown in Supplementary Fig. 9-19 and Supplementary Table 5, respectively. **c**, Hemolytic activity of **1-4**. The presence or absence of R1 and R2 in **1**-**3** are indicated by‘’+’’ and‘’-‘’. All data represent the mean of n=3 biologically independent samples and error bars shows standard deviation. Statistical significance of the difference between results of **1** and **2** are shown (two-tailed t-test: \*\*\*\**P* < 0.0001; \*\*\**P* < 0.001). Individual *P* values are shown in the Source data.

The HPLC-purified product was first analyzed by high-resolution mass spectrometry and tandem mass spectrometry (HR-MS/MS) (Supplementary Fig. 8). The hdmHis was supported by the formation of 4-formyl-1,3-dimethyl-1*H*-imidazolium fragment through a retro aldol reaction^24, 28, 31^. We also assigned a series of y-ions to the precursor peptide, with molecular weight differences implicating serine conversion into alanine and tyrosine glycosylation. NMR analysis was then performed to fully elucidate the structure of lipoavitide **1** which consists of most of the proposed PTMs (Fig. 2, Supplementary Table 5, Supplementary Fig. 9-19). The AviMeCys moiety was formed between the Cys and Thr at the *C*-terminus as expected. The tyrosine modification was uncovered as an *O*-glycosylation by 2-*O*-methyl-*β*-6-deoxygulose. The serine was converted into *D*-alanine, as assessed by Marfey’s assay^32^ (Supplementary Fig. 20). Notably, we also found a 4-hydroxy-2,4-dimethylpentanoyl (HMP) moiety attached to the *N*-terminus, corroborating our initial hypothesis that *lpv* produces an acylated peptide. To explore the uniqueness of HMP and 2-*O*-methyl-*β*-6-deoxygulose, we performed substructures search and found no reported natural products consisting of these units. As the product can be structurally classified as a lipopeptide, we therefore termed this RiPP class lipoavitides (AviMeCys-containing lipopeptides).

## Isolated lipoavitide is a hemolysin

We first tested lipoavitide **1** for antimicrobial activity using a standard agar disk diffusion assay. No growth inhibition was observed for the tested Bacillota, Actinomycetota, Pseudomonadota, and Ascomycota strains (Supplementary Table 6). Owing to their amphiphilicity, many lipopeptides insert into the lipid bilayer and disrupt the integrity of cellular membrane^33^. To evaluate hemolytic activity, lipoavitide **1** was incubated with bovine erythrocytes. We observed cell lysis at low micromolar concentrations (Fig. 2). To investigate if the HMP-modified *N*-terminus contributed to the hemolytic activity, **1** was hydrolyzed under mildly acidic conditions^34^, yielding a peptide with a free *N*-terminus (**2**) and the aglycone of **2** (**3**) (Supplementary Fig. 21-24). The racemic 4-hydroxy-2,4-dimethylpentanoate (**4**) was obtained commercially. Products **2**-**4** were then tested for hemolytic activity with none causing significant cell lysis, suggesting the essentiality of HMP to the hemolytic activity of **1**.

## Production of HMP-CoA via a fatty acid biosynthetic pathway

Bioinformatic analysis suggested that the HMP moiety is produced by the fatty acid/polyketide-like biosynthetic enzymes in *lpv*. Considering the predicted functional annotations of enzymes in *lpv* and the structure of **1**, we hypothesized that HMP biosynthesis starts with condensation of isobutyryl-CoA **5** and methylmalonyl-CoA **6** catalyzed by LpvG, resulting in 3-oxo-2,4-dimethylpentanoyl-CoA **7**. Subsequently, **7** would be successively converted into possible intermediates such as **8** to yield HMP-CoA **9**. LpvE then utilizes **9** as the acyl donor to generate the HMP-modified *N*-terminus (Fig. 3). To confirm or refute this hypothesis, we first disrupted *lpvE* and *lpvG*, respectively, using the RecET recombination system^35^. The resulting BGCs were expressed in *S. albus* J1074, and both produced intermediate **2** with an unmodified *N*-terminus (Fig. 3). We next performed isotopic-labeling experiments to gain insight into HMP biosynthesis. *S. albus* J1074 containing the *lpv*-locus was cultured in ISP4 solid medium supplemented with either [^13^C_5_] *L*-valine or [2-^13^C] propionate to obtain ^13^C-labeled **5** and **6**, respectively^36, 37^ (Supplementary Fig. 25). Isotopically labeled **1** was produced from both substrates as confirmed by MALDI-TOF-MS (Supplementary Fig. 26). Analysis by ^13^C-NMR confirmed isotopic incorporation at C-3, C-4, C-5 and C-7 in HMP when supplementing [^13^C_5_] *L*-valine, and at C-2 in HMP when supplementing [2-^13^C] propionate (Fig. 3, Supplementary Fig. 27).

**Fig. 3.**
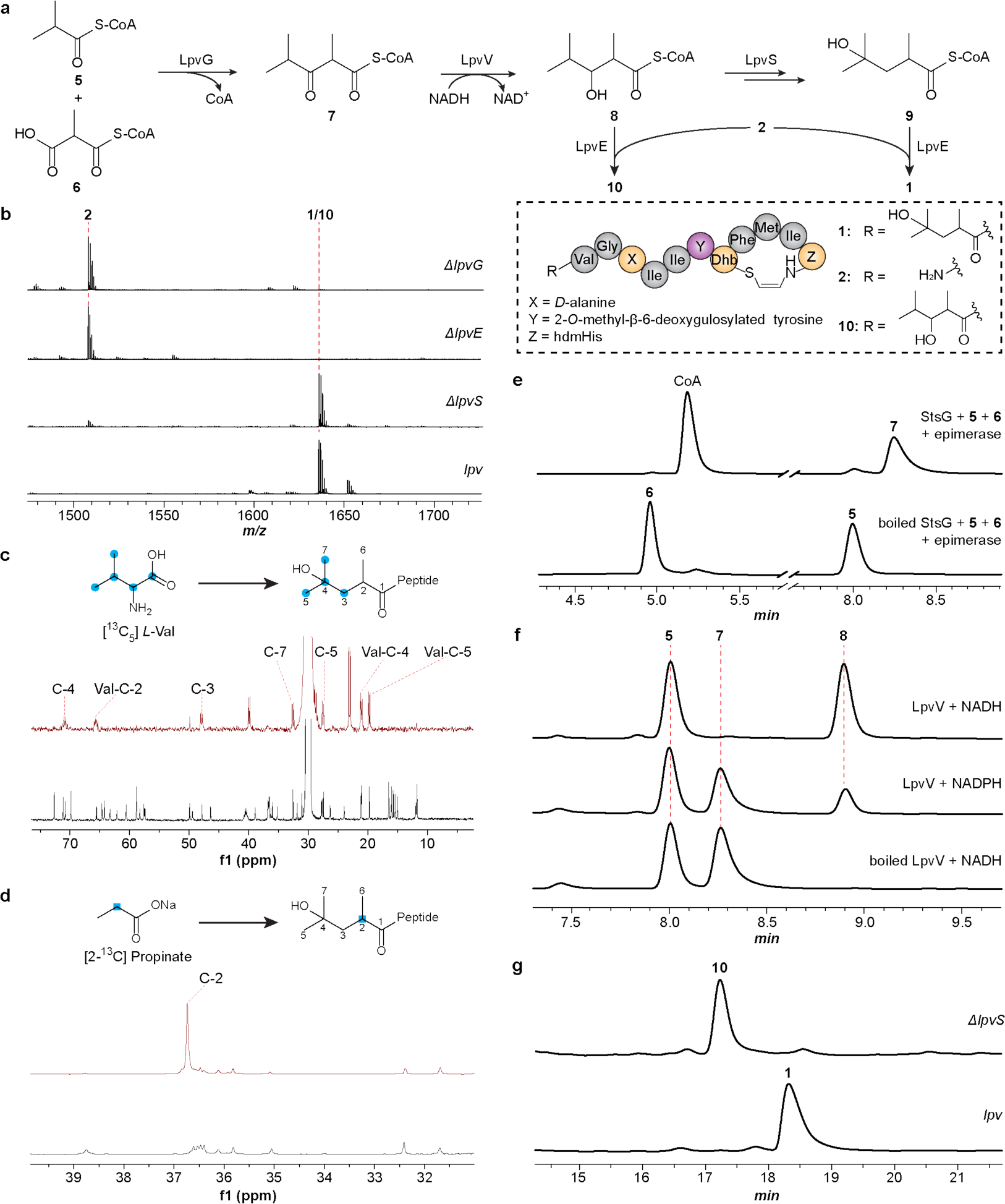
Biosynthesis of the HMP moiety. **a**, Proposed HMP biosynthesis. **b**, MALDI-TOF mass spectra of the methanol extracts of *S. albus* J1074 containing *lpv* and disruption mutants thereof. **c**, Isotopic-labeling of **1** by [^13^C_5_] *L*-valine. Top, ^13^C NMR spectrum of isotopically labeled **1**. Bottom, ^13^C NMR spectrum of unlabeled **1**. Enriched ^13^C signals are indicated in the spectra and assigned to the positions in HMP. Of note, Val1 of **1** is also labeled by [^13^C_5_] *L*-valine and signals are assigned in the spectra accordingly. Full spectra are shown in Supplementary Fig. 27. **d**, Isotopic-labeling of **1** by [2-^13^C] propionate. Top, ^13^C NMR spectrum of isotopically labeled **1**. Bottom, ^13^C NMR spectrum of unlabeled **1**. Enriched ^13^C signals are indicated in the spectrum and assigned to the positions in HMP. Full spectra are shown in Supplementary Fig. 27. **e**, HPLC traces (monitored at 260 nm) of the StsG reaction using **5** and **6** as substrates. **f**, HPLC traces (260 nm) of the StsG/LpvV two-enzyme reaction. **g**, HPLC traces (220 nm) of fractionated methanol extracts of *S. albus* J1074 containing *lpv* and *lpvS*-disrupted constructs.

Next, we characterized LpvG *in vitro*. Unlike FabH from canonical fatty acid synthesis, which requires an ACP-bound extender unit, we predicted LpvG would directly utilize acyl-CoA because *lpv* lacks ACP and FabD homologs (Supplementary Fig. 3). Such ACP-independent activities have been reported in the FabH family^38, 39^. Unfortunately, *E. coli* expression failed to produce soluble LpvG, and thus we examined other FabH-like enzymes from putative orthologous *lpv* BGCs. StsG from *Streptomyces* sp. NRRL S-920 is highly sequence similar to LpvG (70% identity over 94% coverage) and was expressible and purifiable from *E. coli* in a soluble form (Supplementary Fig. 28). Using isobutyryl-CoA **5** and racemic methylmalonyl-CoA **6** as substrates, we reconstituted the activity of StsG *in vitro*, leading to the formation of 3-oxo-2,4-dimethylpentanoyl-CoA **7** as confirmed by HPLC and HR-MS (Supplementary Fig. 29, Supplementary Fig. 30). We observed that even when extended time or higher enzyme concentrations were used, only half of the substrates were consumed. Therefore, we suspected that StsG is selective towards only one diastereomer of **6**. We then conducted the reaction at varying substrate ratios, and the ratio of product **7** to unreacted **6** remained 1:1, even when **5** was supplied in excess. However, when a methylmalonyl-CoA epimerase^40^ was added into the reaction that interconverts the diastereomers of **6**, both **5** and **6** were consumed to completeness as anticipated (Fig. 3, Supplementary Fig. 31).

LpvV, a predicted short-chain dehydrogenase/reductase, was next characterized (Supplementary Table 1, Supplementary Table 2). Disruption of *lpvV* did not alter the production of **1**, suggesting compensation by endogenous *S. albus* enzymes (Supplementary Fig. 32). LpvV was expressed and purified from *E. coli* for *in vitro* characterization (Supplementary Fig. 28). When LpvV was supplied to the StsG reaction with NADH or NADPH, the 3-oxo-2,4-dimethylpentanoyl-CoA **7** was converted to 3-hydroxy-2,4-dimethylpentanoyl (3-HMP)-CoA **8** as monitored by HPLC and HR-MS (Fig. 3, Supplementary Fig. 33). LpvV prefers NADH, as demonstrated by the incomplete conversion of **7** even when NADPH was supplied at a concentration ten times higher than used for NADH. Conversion of 3-HMP-CoA **8** to HMP-CoA **9** putatively occurs through dehydration, reduction, and hydroxylation catalyzed by LpvF, LpvH, and LpvS, respectively (Supplementary Fig. 34). No intermediates were detected from the *lpvF*- or *lpvH*-disrupted BGCs. However, the *lpvS*-disrupted BGC produced **10**, which is isomeric with **1** (Fig. 3, Supplementary Fig. 35). We then isolated the product and determined its structure using NMR, which shares all structural features of **1** but contains a 3-HMP-modified *N*-terminus (Supplementary Table 7, Supplementary Fig. 36, Supplementary Fig. 37). Therefore, we predict that LpvS is critical for the conversion of 3-HMP to HMP, but further experiments are needed to rigorously confirm the biosynthetic route to HMP.

## Acylation of the *N*-terminus by LpvE with substrate promiscuity

From gene disruption studies, we propose LpvE forms **1** via condensation of **2** and **9**. To further characterize its acyltransferase activity *in vitro*, LpvE was expressed in *E. coli* and purified (Supplementary Fig. 28). Although substrate **2** could be hydrolytically prepared (Supplementary Fig. 21), obtaining HMP-CoA **9** was considerably more challenging. However, isolation of **10** suggested that LpvE can also use 3-HMP-CoA **8** as a substrate, allowing us to examine the activity of LpvE, since **8** can be generated by the FabH enzyme StsG and the short-chain dehydrogenase/reductase LpvV. Therefore, we supplied LpvE and **2** to the StsG/LpvV reaction, which was expected to yield **10**. MS analysis of the reaction products showed a prominent peak at *m/z* 1636, consistent with formation of **10** (Fig. 4). The identity of *m/z* 1636 was confirmed through HPLC co-elution of the reaction mixture and a standard of **10** produced from the *lpvS*-disrupted BGC (Fig. 4, Supplementary Fig. 38). Omission of StsG or LpvV from the reaction abolished production of **10**. However, mass spectral analysis revealed the addition of isobutyryl or 3-oxo-2,4-dimethylpentanoyl moieties to **2**, yielding **11** and **12**, implying that LpvE tolerates isobutyryl-CoA **5** and 3-oxo-2,4-dimethylpentanoyl-CoA **7** as acyl donors (Supplementary Fig. 39). The inability of LpvE to use methylmalonyl-CoA **6** also suggested that LpvG-catalyzed condensation occurs prior to acylation. To further evaluate LpvE, 2,4-dimethylpentanoyl-CoA was prepared using a carbonyldiimidazole (CDI)-activation strategy^41^ (Supplementary Fig. 40). The resulting acyl donor successfully reacted with **2** requiring only LpvE as the catalyst, providing additional evidence that LpvE is an acyltransferase (Supplementary Fig. 41).

**Fig. 4.**
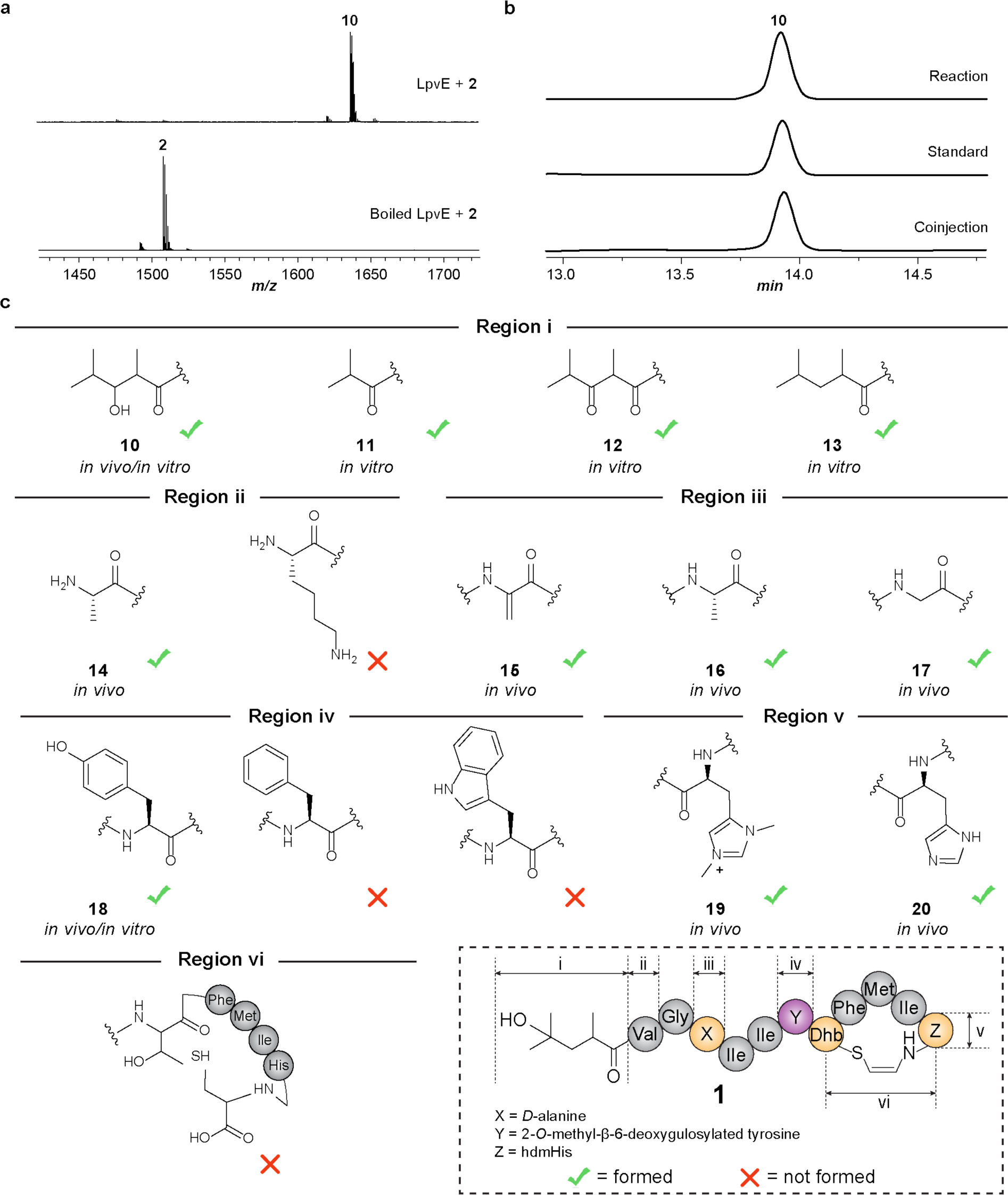
Characterization of the acyltransferase LpvE. **a**, MALDI-TOF mass spectra of the StsG/LpvV/LpvE reaction. **b**, HPLC analysis of the StsG/LpvV/LpvE three-enzyme reaction. Shown are chromatograms (280 nm) for the reaction product (top), standard isolated from *S. albus* J1074 containing the *lpvS*-deleted *lpv* (middle), and a 1:1 co-injection of these two samples (bottom). **c**, Summary of structural variants of **1** generated by *in vivo* and *in vitro* approaches. The structure of **1** is shown for reference.

Since LpvE tolerated various acyl donors, we next investigated tolerance towards various peptide substrates. We first asked if modified residues in the linear region of **1** would affect acylation. Val1 of the core peptide was replaced with Ala using RecET-assisted recombination, yielding **14**, which is 28 Da less than **1**, suggesting that the product harbored an HMP-modified *N*-terminus (Fig. 4, Supplementary Fig. 42). However, when Val1 was substituted with Lys, no expected product was detected by MS, possibly due to interference in acylation or leader peptide removal. Acylated products were also observed upon disruption of LpvJ (luciferase-like monooxygenase) or substitution of the core residue Ser3, suggesting acylation was independent of *D*-alanine formation (Supplementary Fig. 43). Disrupting LpvI (glycosyltransferase) led to the formation of acylated products lacking glycosylated tyrosine. However, substitution of this residue with either phenylalanine or tryptophan abolished product synthesis (Supplementary Fig. 44). Additionally, the *in vitro* acylation of **3** was significantly slower than for **2**, indicating glycosylation may occur before acylation (Supplementary Fig. 45).

We next investigated the LpvE tolerance to the PTMs in the macrocyclic region. The hdmHis moiety exerts little impact on acylation, as acylated products consisting of unmodified and dimethylated histidine were observed upon disruption of LpvM1 and LpvK, respectively^28, 29^ (Supplementary Fig. 46). Nevertheless, *in vitro* acylation of linear peptide VG^D^AIIYTFMIHC was unsuccessful, which revealed that the AviMeCys macrocyclization was required for LpvE-catalyzed acylation (Supplementary Fig. 47).

To shed more light on the promiscuous activity of LpvE, we predicted the protein structure by AlphaFold2^42^ and performed molecular docking studies with AutoDock^43^ (Supplementary Fig. 48). The *N*-terminus of **2** and the HMP moiety of **9** were directed into the putative active site for condensation. Most of the other residues in **2** are located outside the substrate-binding pocket of LpvE, possibly explaining the tolerance to structural variations at these sites. In contrast, an analog of **2** without the AviMeCys was randomly docked with LpvE, indicating the structural rigidity afforded by the macrocycle in **2** might be essential. Phylogenetic analysis revealed that lipoavitide acyltransferases form a distinct clade within the GCN5-related *N*-acetyltransferase (GNAT) superfamily (Supplementary Fig. 49, Supplementary Table 8). To further investigate the substrate tolerance of these enzymes, we successfully reconstituted the activity of StsE *in vitro* using both **2** and **3** as substrates, even though 8 of the 12 residues in the StsA core peptide are different from that in LpvA (Fig. 4, Supplementary Fig. 28, Supplementary Fig. 50). We anticipate that lipoavitide acyltransferases will use a broad range of substrates, making them ideal candidate biocatalysts for lipopeptide synthesis.

## Discussion

Advances in DNA sequencing and bioinformatics have enabled the detection of hybrid BGCs with increasing frequency. These pathways often produce natural products consisting of structural characteristics that span multiple families, of which the genome mining can significantly expand natural product structural diversity and biosynthetic routes^44–46^. However, only a few RiPP/non-RiPP hybrid BGCs have been discovered. Here, we leveraged the CAPTURE method to enable rapid characterization of high complexity BGCs. In doing so, we uncovered a class of RiPP/fatty acid hybrid BGCs that produce lipoavitides characterized by a unique fatty acyl moiety (HMP), an AviMeCys macrocycle, and other modified amino acids. Further experiments have confirmed the role of hybrid biosynthesis in forming the fatty acyl and peptide moieties.

Lipoavitides are structurally categorized as lipopeptides, a class of natural products known for their amphiphilic properties^47^. Lipopeptides typically contain a hydrophobic moiety composed of fatty acyl units varying in length and functionalities, connected to a hydrophilic peptide that often contains macrocycles and non-proteinogenic amino acids. These molecules are characterized by a myriad of biological functions and have shown tremendous potential in medical applications, as shown by the clinical usage of lipopeptide antibiotics including daptomycin^48^, polymyxin^49^, and echinocandin^50^. Although nature has developed multiple pathways to synthesize peptidic natural products, known lipopeptides are predominantly synthesized by NRPSs, which function as megaenzymatic assembly lines capable of incorporating a wide range of amino acid building blocks. To date, ribosomal synthesis has only been found to produce a small subset of lipopeptides, such as lipolanthins, goadvionins, and selidamides. In this study, we characterized the lipoavitides as ribosomally synthesized lipopeptides. Structure-activity relationship studies demonstrated the essential role of the fatty acyl moiety in hemolytic activity, suggesting it aids interaction with cellular membranes. Due to the versatility of lipoavitide biosynthesis, characterization of lipoavitide BGCs may also offer valuable tools in the engineering of lipopeptides for drug development purposes.

Among ribosomal peptides characterized with Avi(Me)Cys moieties, thioviridamide-like thioamitides are most structurally related to lipoavitides. In addition to the presence of hdmHis, thioamitides also resemble lipoavitides with an acylated *N*-terminus (Supplementary Table 4). However, the acyl moieties of thioamitides, specifically pyruvyl and lactyl, are formed via dehydration and tautomerization of *N*-terminal serine residues^31^, whereas lipoavitides employ fatty acid biosynthetic enzymes to produce fatty acyl-CoAs, which can be transferred to the *N*-terminus. In addition, lipoavitides contain modified residues including *D*-alanine and glycosylated tyrosine, while thioamitides contain thioamides in the *N*-terminal region. These structural differences of lipoavitides and thioamitides may confer disparate biological functions which deserve further investigation.

LpvG was characterized as a FabH-like enzyme, which initiates the synthesis of HMP via condensation of isobutyryl-CoA **5** and methylmalonyl-CoA **6**. FabH enzymes possessing similar activities were also reported to produce other fatty acylated natural products. One such example is trehangelin, an oligosaccharide natural product harboring a unique fatty acyl group named angelyl^39^. ThgI, the FabH enzyme encoded in the trehangelin BGC, condenses acetyl-CoA and methylmalonyl-CoA, yielding 2-methyl-acetoacetyl-CoA that is further converted to angelyl-CoA by the ketoreductase ThgK and enoyl-CoA dehydratase ThgH successively. Similar biosynthetic enzymes were also identified in a type II polyketide BGC, producing the tetracycline SF2575 that also comprise an angelyl moiety^51^. With the continued efforts toward mining genomes, we envision that FabH enzymes will be increasingly implicated in the biosynthesis of natural products with unique fatty acyl moieties.

Acyltransferases of the GNAT family are widely distributed in RiPP BGCs. While previously known for *N*-acetylation of LAPs^52^, lasso peptides^53^, and graspetides^54^, the recent discoveries of lipolanthins^6, 7^, goadvionins^8^, and selidamides^55^, have shown their ability to incorporate relatively long chain fatty acyl units into RiPPs. GdvG, the acyltransferase involved in goadvionin biosynthesis, can append a fatty acyl group of 32 carbons to an avionin-containing peptide. However, GdvG only accepts ACP-bound fatty acyl units, which somewhat restricts its application as a biocatalyst. In this study, the lipoavitide acyltransferase LpvE accepts fatty acyl-CoA substrates varying in length and functionalities. In addition, LpvE and its homologous enzyme StsE revealed high degree of substrate tolerance. Such promiscuity makes these enzymes promising biocatalysts, offering valuable opportunities for the preparation of lipopeptide libraries and engineering of new-to-nature lipopeptides.

## Acknowledgments

We wish to thank Justine Arrington from the Roy J. Carver Biotechnology Center for HR-MS/MS assistance. This work was supported in part by grants from the National Institutes of Health (AI144967).

## Author contributions

HR and CH contributed equally to this work. HR performed bioinformatics analysis, direct cloning, heterologous expression, HR-MS/MS analysis, bioactivity assays, mutational analysis, and enzymatic reactions. CH performed compound purifications, isotopic feeding study, and intermediate characterization. CH also designed, performed, and analyzed NMR experiments. YP assisted in compound purifications and genetic manipulation. HC performed AlphaFold2 and AutoDock analysis. DAM contributed to the bioinformatics analysis of *lpv* genes. SRD performed stereochemical group determinations with oversight from DAM. HR and CH wrote the manuscript with editorial oversight from HZ, along with input from all other authors. HZ conceived of and supervised the overall project.

## Competing interests

The authors declare no competing interests.

## Additional information

**Supplementary information** is available for this paper at https:

## Data and materials availability

We declare that all data supporting the findings of this study are presented in the main text and supplementary information. NCBI (https://www.ncbi.nlm.nih.gov/), PDB (https://www.rcsb.org/), and Uniprot (https://www.uniprot.org/) accessions are referenced in the supplementary information, and these accessions are publicly accessible on the respective NCBI, RCSB PDB, and Uniprot websites. Data is available from the corresponding authors upon request. Source data are provided with this paper.

## Notes

### Competing Interest Statement

The authors have declared no competing interest.

## References and Notes

1. Arnison, P.G. et al. Ribosomally synthesized and post-translationally modified peptide natural products: Overview and recommendations for a universal nomenclature. Nat Prod Rep 30, 108–160 (2013).

2. Montalban-Lopez, M. et al. New developments in RiPP discovery, enzymology and engineering. Nat Prod Rep 38, 130–239 (2021).

3. Just-Baringo, X., Albericio, F. & Alvarez, M. Thiopeptide antibiotics: Retrospective and recent advances. Mar Drugs 12, 317–351 (2014).

4. Noike, M. et al. A peptide ligase and the ribosome cooperate to synthesize the peptide pheganomycin. Nat Chem Biol 11, 71–76 (2015).

5. Ortiz-Lopez, F.J. et al. Cacaoidin, first member of the new lanthidin ripp family. Angew Chem Int Ed 59, 12654–12658 (2020).

6. Wiebach, V. et al. The anti-staphylococcal lipolanthines are ribosomally synthesized lipopeptides. Nat Chem Biol 14, 652–654 (2018).

7. Wiebach, V. et al. An amphipathic alpha-helix guides maturation of the ribosomally-synthesized lipolanthines. Angew Chem Int Ed 59, 16777–16785 (2020).

8. Kozakai, R. et al. Acyltransferase that catalyses the condensation of polyketide and peptide moieties of goadvionin hybrid lipopeptides. Nat Chem 12, 869–877 (2020).

9. Cao, L., Do, T. & Link, A.J. Mechanisms of action of ribosomally synthesized and posttranslationally modified peptides (RiPPs). J Ind Microbiol Biotechnol 48, kuab005 (2021).

10. Ongpipattanakul, C. et al. Mechanism of action of ribosomally synthesized and post-translationally modified peptides. Chem Rev 122, 14722–14814 (2022).

11. Melby, J.O., Nard, N.J. & Mitchell, D.A. Thiazole/oxazole-modified microcins: Complex natural products from ribosomal templates. Curr Opin Chem Biol 15, 369–378 (2011).

12. Franz, L., Kazmaier, U., Truman, A.W. & Koehnke, J. Bottromycins-biosynthesis, synthesis and activity. Nat Prod Rep 38, 1659–1683 (2021).

13. Vinogradov, A.A. & Suga, H. Introduction to thiopeptides: Biological activity, biosynthesis, and strategies for functional reprogramming. Cell Chem Biol 27, 1032–1051 (2020).

14. McIntosh, J.A., Donia, M.S. & Schmidt, E.W. Insights into heterocyclization from two highly similar enzymes. J Am Chem Soc 132, 4089–4091 (2010).

15. Burkhart, B.J., Schwalen, C.J., Mann, G., Naismith, J.H. & Mitchell, D.A. YcaO-dependent posttranslational amide activation: Biosynthesis, structure, and function. Chem Rev 117, 5389–5456 (2017).

16. Norris, G.E. & Patchett, M.L. The glycocins: In a class of their own. Curr Opin Struc Biol 40, 112–119 (2016).

17. Saad, H. et al. Nocathioamides, uncovered by a tunable metabologenomic approach, define a novel class of chimeric lanthipeptides. Angew Chem Int Ed 60, 16472–16479 (2021).

18. Medema, M.H., Cimermancic, P., Sali, A., Takano, E. & Fischbach, M.A. A systematic computational analysis of biosynthetic gene cluster evolution: Lessons for engineering biosynthesis. Plos Comput Biol 10, e1004016 (2014).

19. Robey, M.T., Caesar, L.K., Drott, M.T., Keller, N.P. & Kelleher, N.L. An interpreted atlas of biosynthetic gene clusters from 1,000 fungal genomes. Proc Natl Acad Sci USA 118, e2020230118 (2021).

20. Gavriilidou, A. et al. Compendium of specialized metabolite biosynthetic diversity encoded in bacterial genomes. Nat Microbiol 7, 726–735 (2022).

21. Walsh, C.T., Brien, R.V.O. & Khosla, C. Nonproteinogenic amino acid building blocks for nonribosomal peptide and hybrid polyketide scaffolds. Angew Chem Int Ed 52, 7098–7124 (2013).

22. Blin, K. et al. AntiSMASH 6.0: Improving cluster detection and comparison capabilities. Nucleic Acids Res 49, W29–W35 (2021).

23. Grant-Mackie, E.S., Williams, E.T., Harris, P.W.R. & Brimble, M.A. Aminovinyl cysteine containing peptides: A unique motif that imparts key biological activity. Jacs Au 1, 1527–1540 (2021).

24. Eyles, T.H., Vior, N.M., Lacret, R. & Truman, A.W. Understanding thioamitide biosynthesis using pathway engineering and untargeted metabolomics. Chem Sci 12, 7138–7150 (2021).

25. Xu, M. et al. Functional genome mining reveals a class V lanthipeptide containing a d-amino acid introduced by an F420H2-dependent reductase. Angew Chem Int Ed 59, 18029–18035 (2020).

26. Kloosterman, A.M. et al. Expansion of RiPP biosynthetic space through integration of pan-genomics and machine learning uncovers a novel class of lantibiotics. Plos Biol 18, e3001026 (2020).

27. Schujman, G.E. & de Mendoza, D. Regulation of type II fatty acid synthase in Gram-positive bacteria. Curr Opin Microbiol 11, 148–152 (2008).

28. Hu, L. et al. Characterization of histidine functionalization and its timing in the biosynthesis of ribosomally synthesized and posttranslationally modified thioamitides. J Am Chem Soc 144, 4431–4438 (2022).

29. Sikandar, A., Lopatniuk, M., Luzhetskyy, A., Muller, R. & Koehnke, J. Total in vitro biosynthesis of the thioamitide thioholgamide andinvestigation of the pathway. J Am Chem Soc 144, 5136–5144 (2022).

30. Enghiad, B. et al. Cas12a-assisted precise targeted cloning using in vivo Cre-lox recombination. Nat Commun 12, 1171 (2021).

31. Frattaruolo, L., Lacret, R., Cappello, A.R. & Truman, A.W. A genomics-based approach identifies a thioviridamide-like compound with selective anticancer activity. ACS Chem Biol 12, 2815–2822 (2017).

32. Bhushan, R. & Bruckner, H. Use of Marfey’s reagent and analogs for chiral amino acid analysis: Assessment and applications to natural products and biological systems. J Chromatogr B 879, 3148–3161 (2011).

33. Bender, C.L., Alarcon-Chaidez, F. & Gross, D.C. Pseudomonas syringae phytotoxins: Mode of action, regulation, and biosynthesis by peptide and polyketide synthetases. Microbiol Mol Biol R 63, 266–292 (1999).

34. Nilsson, J. et al. Enrichment of glycopeptides for glycan structure and attachment site identification. Nat Methods 6, 809–811 (2009).

35. Fu, J. et al. Full-length RecE enhances linear-linear homologous recombination and facilitates direct cloning for bioprospecting. Nat Biotechnol 30, 440–446 (2012).

36. Tang, L., Zhang, Y.X. & Hutchinson, C.R. Amino acid catabolism and antibiotic synthesis: Valine is a source of precursors for macrolide biosynthesis in Streptomyces ambofaciens and Streptomyces fradiae. J Bacteriol 176, 6107–6119 (1994).

37. Dayem, L.C. et al. Metabolic engineering of a methylmalonyl-CoA mutase-epimerase pathway for complex polyketide biosynthesis in Escherichia coli. Biochemistry 41, 5193–5201 (2002).

38. Okamura, E., Tomita, T., Sawa, R., Nishiyama, M. & Kuzuyama, T. Unprecedented acetoacetyl-coenzyme A synthesizing enzyme of the thiolase superfamily involved in the mevalonate pathway. Proc Natl Acad Sci USA 107, 11265–11270 (2010).

39. Inahashi, Y. et al. Biosynthesis of trehangelin in polymorphospora rubra k07-0510: Identification of metabolic pathway to angelyl-CoA. Chembiochem 17, 1442–1447 (2016).

40. Bobik, T.A. & Rasche, M.E. Identification of the human methylmalonyl-CoA racemase gene based on the analysis of prokaryotic gene arrangements - implications for decoding the human genome. J Biol Chem 276, 37194–37198 (2001).

41. Peter, D.M., Vogeli, B., Cortina, N.S. & Erb, T.J. A chemo-enzymatic road map to the synthesis of CoA esters. Molecules 21, 517 (2016).

42. Jumper, J. et al. Highly accurate protein structure prediction with AlphaFold. Nature 596, 583–589 (2021).

43. Morris, G.M., Goodsell, D.S., Huey, R. & Olson, A.J. Distributed automated docking of flexible ligands to proteins: Parallel applications of AutoDock 2.4. J Comput Aided Mol Des 10, 293–304 (1996).

44. Park, H.B., Perez, C.E., Barber, K.W., Rinehart, J. & Crawford, J.M. Genome mining unearths a hybrid nonribosomal peptide synthetase-like-pteridine synthase biosynthetic gene cluster. Elife 6, e25229 (2017).

45. Schor, R., Schotte, C., Wibberg, D., Kalinowski, J. & Cox, R.J. Three previously unrecognised classes of biosynthetic enzymes revealed during the production of xenovulene A. Nat Commun 9, 1963 (2018).

46. Yee, D.A. et al. Genome mining of alkaloidal terpenoids from a hybrid terpene and nonribosomal peptide biosynthetic pathway. J Am Chem Soc 142, 710–714 (2020).

47. Gotze, S. & Stallforth, P. Structure elucidation of bacterial nonribosomal lipopeptides. Org Biomol Chem 18, 1710–1727 (2020).

48. Robbel, L. & Marahiel, M.A. Daptomycin, a bacterial lipopeptide synthesized by a nonribosomal machinery. J Biol Chem 285, 27501–27508 (2010).

49. Poirel, L., Jayol, A. & Nordmann, P. Polymyxins: Antibacterial activity, susceptibility testing, and resistance mechanisms encoded by plasmids or chromosomes. Clin Microbiol Rev 30, 557–596 (2017).

50. Huttel, W. Echinocandins: Structural diversity, biosynthesis, and development of antimycotics. Appl Microbiol Biotechnol 105, 55–66 (2021).

51. Pickens, L.B. et al. Biochemical analysis of the biosynthetic pathway of an anticancer tetracycline SF2575. J Am Chem Soc 131, 17677–17689 (2009).

52. Ozaki, T. et al. Dissection of goadsporin biosynthesis by in vitro reconstitution leading to designer analogues expressed in vivo. Nat Commun 8, 14207 (2017).

53. Zong, C., Cheung-Lee, W.L., Elashal, H.E., Raj, M. & Link, A.J. Albusnodin: An acetylated lasso peptide from Streptomyces albus. Chem Commun (Camb) 54, 1339–1342 (2018).

54. Ziemert, N., Ishida, K., Liaimer, A., Hertweck, C. & Dittmann, E. Ribosomal synthesis of tricyclic depsipeptides in bloom-forming cyanobacteria. Angew Chem Int Ed 47, 7756–7759 (2008).

55. Hubrich, F. et al. Ribosomally derived lipopeptides containing distinct fatty acyl moieties. Proc Natl Acad Sci USA 119, e2113120119 (2022).

