## Supplementary Materials for "Non-modular Fatty Acid Synthases Yield Unique Acylation in Ribosomal Peptides"

### Materials and Methods

**Bacterial strains and reagents.** *E. coli* strain NEB10β and BL21(DE3) were purchased from New England Biolabs (Ipswich, MA) and used for cloning and protein expression experiments, respectively. *E. coli* strain WM6026, a gift from William Metcalf (University of Illinois at Urbana-Champaign), was used as the donor strain for conjugation to *Streptomyces* heterologous hosts[1]. *Streptomyces albus* J1074 and *Streptomyces lividans* TK24 were used for heterologous expression of *lpv*. *Streptomyces* sp. NRRL S-1521, *Streptomyces* sp. NRRL S-920, *S. albus* J1074, and *S. lividans* TK24 were obtained from the Agricultural Research Service Culture Collection (Peoria, IL). Chemicals were purchased from ThermoFisher Scientific (Waltham, MA). No further purification was performed for any purchased chemicals before use. Restriction enzymes, NEBuffers, T4 DNA polymerase, *E. coli* DNA ligase, dNTPs, $NAD^+$, NEBuilder HiFi DNA assembly mastermix, and Q5 DNA polymerase were purchased from New England Biolabs (Ipswich, MA). All DNA oligonucleotides were ordered from Integrated DNA Technologies (Coralville, IA).

**Generation of acyltransferase phylogenetic tree.** The acyltransferases for lipoavitide, lipolanthin, and goadvionin biosynthesis, along with other characterized members of the GNAT families, were aligned by Mega 11 using the ClustalW method[2]. A maximum-likelihood phylogenetic tree was then generated based on this alignment with the Jones-Taylor-Thornton (JTT) model using 1000 bootstrap replications. The tree was visualized using the Interactive Tree of Life (iTOL) website (https://itol.embl.de/).

**Direct cloning of the *lpv* BGC from *Streptomyces* sp. NRRL S-1521.** Direct cloning of *lpv* was achieved by the CAPTURE method[3]. Briefly, the *Streptomyces* sp. NRRL S-1521 was recovered on ISP2 agar medium (malt extract 10 g/L, yeast extract 4 g/L, glucose 4 g/L, agar 20g/L, pH 7.2–7.4) at 37 °C until colony appears (about 3 days). A single colony was inoculated into 5 ml ISP2 liquid medium and grown at 37 °C 250 rpm until saturation (3 d). An aliquot of 1 mL cell culture was then transferred into 50 mL fresh ISP2 liquid medium and cultivated for 18-20 h. Cells were then harvested by centrifugation at 3000 × g for 15 min. All the rest of the direct cloning experiments, including the genomic DNA isolation, *in vitro* gRNA transcription, DNA receivers preparation, genomic DNA digestion and Cre-lox *in vivo* recombination, were performed following the CAPTURE method as described elsewhere[3]. Sequences of the DNA templates for the *in vitro* gRNA transcription were shown in **Supplementary Table 7** and synthesized by Integrated DNA Technologies (Corallville, IA). The cloned BGC was then transformed into WM6026 (supplemented with 2,6-diaminopimelic acid at the final concentration 40 µg/mL for growth) and conjugated into *S. lividans* TK24 and *S. albus* J1074 for expression by the protocol described elsewhere[4].

**Product analysis from colony extracts.** The exconjugants of *S. lividans* TK24 and *S. albus* J1074 containing *lpv* or the empty pBE45 vector were selected and grew on fresh MS (mannitol 20 g/L, soybean flour 20 g/L, agar 20 g/L) and ISP4 (soluble starch 10 g/L, $K_2HPO_4$ 1 g/L, $MgSO_4 \cdot 7H_2O$ 1 g/L, NaCl 1 g/L, $(NH_4)_2SO_4$ 2 g/L, $CaCO_3$ 2 g/L, $FeSO_4$ 1 mg/L, $MnCl_2$ 1 mg/L, $ZnSO_4$ 1 mg/L, agar 20 g/L) agar medium supplied with apramycin at a final concentration of 50 µg/mL and incubated under 30 °C for 5 d. A pinhead-sized portion of cell mass was picked from the plate, placed in 20 µL methanol and incubated at room temperature for 1 h. An aliquot of 1 µL methanol extract was then mixed with an equal volume of 15 mg/mL 70% aq. acetonitrile solution of α-cyano-4-hydroxycinnamic acid (CHCA) with 0.1% trifluoroacetic acid (TFA) (*v/v*) on a ground steel MALDI target, and the droplet was dried under ambient conditions. Samples were analyzed using a Bruker UltrafleXtreme MALDI-TOF MS for reflector positive mode.

**Heterologous expression and product isolation.** Freshly obtained exconjugants of *S. albus* J1074 containing wild type or mutated *lpv* were individually grew on MS plates with apramycin and incubated under 30 °C for 5 d. Colonies were verified for producing the lipoavitide by MALDI-TOF MS by the method described above. The colonies were then scratched individually from the plate by sterile cotton swabs, spread on fresh ISP4 plates, and allowed to grow for another 5 d. The ISP4 plates were then extracted by methanol under 4 °C overnight. Methanol extracts were then harvested by filtration, mixed with an equal volume of water and loaded onto an Agilent Bond Elut C18 Solid Phase Extraction (SPE) column (bed mass, 10 g; volume, 60 mL; particle size 120 µm), which was pre-equilibrated by 50 mL 5% B (solvent A

= 0.1% TFA in water; solvent B = 0.1% TFA in acetonitrile). The compounds were then eluted using a step gradient with increasing percentage of solvent B in 150 mL volumes: 5%, 20%, 40%, 60%, 80% and 100% B. The eluted compounds were monitored by MALDI-TOF mass spectrometry. Fractions containing the compounds (60% B) were lyophilized to dryness and powders were redissolved into methanol. Semi-preparative HPLC purification was performed using a 1290 Infinity II Preparative LC System equipped with a Phenomenex Luna C5 column (5 µm, 100 Å, 250 × 10 mm) equilibrated in 5% B. Compounds were eluted by an increase to 100% B over 20 min with a flow rate of 3 mL/min. Collected fractions were analyzed by MALDI-TOF mass spectrometry, lyophilized to dryness, and stored at -80 °C until further use. Regarding to the feeding study, fermentation and product purification of **1** followed the same methods except that [$^{13}$C] L-valine and [$^{13}$C] propionate were supplemented into the ISP4 solid medium to a final concentration of 2 mg/mL.

**Preparation of 2 and 3.** Hydrolysis of **1** was performed by dissolving the product in 100 mM formic acid to a final concentration of 1 mg/mL, which was then incubated in Eppendorf ThermoMixer C with a heated lid for 20 h at 80 °C. Afterwards, the solution was purified by semi-preparative HPLC using a 1290 Infinity II Preparative LC System equipped with a Phenomenex Luna C5 column (5 µm, 100 Å, 250 × 10 mm) equilibrated in 30% B. Compounds were eluted by an increase to 80% B over 10 min with a flow rate of 3 mL/min. Under these conditions, **2** and **3** were eluted at 7.6 and 7.8 min, respectively. Collected fractions were analyzed by MALDI-TOF mass spectrometry, lyophilized to dryness, and stored at -80 °C until further use.

**High-resolution mass spectrometry.** Lyophilized, HPLC-purified **1** was resuspended in methanol. Samples were directly infused onto a Thermo Scientific Q Exactive HF-X hybrid quadrupole-Orbitrap ESI-MS using an Advion TriVersa Nanomate 100. Calibration was performed using Pierce LTQ Velos ESI Positive Ion Calibration Solution (ThermoFisher). The MS was operated with the following parameters: 100,000 resolution, 2 *m/z* isolation width (MS/MS), 0.4 activation q value (MS/MS), and 30 ms activation time (MS/MS). Fragmentation was performed using collision-induced dissociation (CID) at normalized collision energy. Each peptide was subjected to 30 normalized collision energy (MS/MS). Data analysis was conducted using the Qualbrowser application of Xcalibur software (version 4.1.31.9, ThermoFisher Scientific).

**NMR spectroscopy.** The sample was prepared by dissolving ~5 mg of the HPLC-purified product in 500 µL of acetone-$d_6$ (99.9 atom % D, Sigma-Aldrich). NMR spectra were recorded on an Agilent VNMRS 750 MHz narrow bore magnet spectrometer equipped with a 5 mm triple resonance ($^1$H-$^{13}$C-$^{15}$N) triaxial gradient probe and pulse-shaping capabilities. Samples were held at 25 °C during acquisition. Standard Varian pulse sequences were used for the following experiments: $^1$H, $^{13}$C, $^1$H-$^1$H COSY, $^1$H-$^1$H TCOSY, $^1$H-$^{13}$C HSQC, and $^1$H-$^{13}$C HMBC. Spectra were recorded with VNMRJ 3.2A software. All NMR data were processed using MestReNova 11.0.3. Chemical shifts were referenced internally to the solvent peak ($\delta_H$ = 2.05, $\delta_c$ = 29.8).

**Antimicrobial activity assay.** Lipoavitide **1** was dissolved in methanol to achieve a concentration of 10 µM. Agar plates were prepared by mixing 200 µL of stationary phase overnight cell culture with 20 mL of melted solid medium (cooled to 42 °C for 5 min). The seeded agar was poured into a sterile 100-mm round dish (VWR) and allowed to solidify at 25 °C for 10 min. Lipoavitide **1** was directly spotted on the solidified agar. Plates were incubated at temperatures according to **Supplementary Table 4** for 16 h, and the antimicrobial activity was determined by the presence or absence of inhibition zones.

**Hemolytic Assay.** Fresh defibrinated whole bovine blood was purchased from Hemostat Laboratories. Whole blood was washed three times, diluted to a final concentration of 1:25 (*v/v*) using PBS, and then split into 50 µL aliquots in individual 1.7 mL Eppendorf tubes. Next, stock solutions of concentrations at 1 mM, 200 µM, and 40 µM were prepared for **1**, **2**, **3**, and **4** in methanol. Aliquots of 2.5 µL each stock solution were mixed with the blood, yielding final concentrations at 50 µM, 10 µM, and 2 µM. An equal volume of methanol and Triton X-100 were used as negative and positive controls. The mixtures were then incubated in Eppendorf ThermoMixer C with a heated lid for 18 h at 37 °C. After incubation, the samples were centrifugated at 500 × g for 10 min, and the supernatants were measured for hemoglobin absorbance at 410 nm on a NanoDrop Spectrophotometer. Each measurement was performed in biological triplicate, and the percentage of hemolysis was calculated using the following equation:

$$\% \text{ Hemolysis} = \frac{\text{Abs of test sample} - \text{Abs of DMSO}}{\text{Abs of Triton X} - 100 - \text{Abs of DMSO}} \quad (1)$$

**Characterization of stereochemistry by Marfey's assays.** Purified **1** (250 µg) was dried under vacuum in a Schlenk flask. To the dried peptide, 300 µL of a 1:1 mixture 12 M DCl (35% *w/v*, Aldrich) and $D_2O$ (Cambridge Isotope Laboratory) was added. The solution was equilibrated under nitrogen gas and subjected to one freeze-pump-thaw cycle. The sample was then frozen, subjected to vacuum, and sealed. Once equilibrated back to room temperature, the sample was heated to 95 °C for 12 h under reduced pressure. The flask was removed from heat, and once cooled, insoluble debris was removed by centrifugation for 5 min. Supernatant was removed and dried using a Speedvac concentrator (Savant ISS110).

The hydrolysate of **1** was dissolved in 150 µL 1 M aq. $NaHCO_3$. An acetone solution containing 4 mg/mL 1-fluoro-2-4-dinitrophenyl-5-L-alanine amide (L-FDAA, Thermo Scientific) was prepared and 150 uL was added to the reconstituted hydrolysate. Contents were mixed, and the reaction vessel was heated to 60 °C for 2 h without shaking using an Eppendorf ThermoMixer C. The reaction mixture was then quenched using 50 µL 6 M HCl dropwise. Sample was dried using a Speedvac concentrator (Savant ISS110). Amino acid standards were prepared by drying down 100 µL L-amino acid solution (0.5 µmol/mL), followed by dissolution in 1 M aq. $NaHCO_3$ and reaction with L-FDAA as above. D-amino standards were generated by reacting the L-amino acid solution with 1-fluoro-2-4-dinitrophenyl-5-D-alanine amide (D-FDAA, Toronto Research Chemicals). These reaction products are enantiomers to the hypothetical D-amino acid and L-FDAA coupled products, where both are chromatographically identical using an achiral stationary phase.

Derivatized amino acid standards and hydrolysate of **1** were reconstituted in 300 µL 15% aq. acetonitrile + 0.1% formic acid (LC-MS grade). Samples were sonicated for 15 min in a sonication bath and insoluble debris was removed by centrifugation. Amino acid stereochemistry was determined using a Shimadzu LC-MS 2020 with a quadrupole detector and a Macherey-Nagel Nucleodur C18 column (250 × 4.6 mm, 5 µm particle size, 100 Å pore size). Samples were analyzed in batch using the following conditions: solvent A 0.1% aq. formic acid, solvent B acetonitrile + 0.1% formic acid, 1 mL/min flow rate, and the following method: 15% B for 7 min, 15-50% B over 45 min, hold at 50% B for 5 min followed by wash and re-equilibration in starting conditions. Analyte (20 µL) was injected and the $[M+H]^+$ ions of the amino acid-Marfey's reagent adducts were selectively monitored by electrospray ionization in positive mode as follows (*m/z*): Ala, 342; Phe, 418; Ile/Leu, 384; Met, 402; Val, 370.

**Gene disruption and LpvA point mutation by RecET recombineering.** The genetic manipulation of *lpv* was achieved using the RecET recombination system. Donor DNA fragments containing an ampicillin resistance marker flanked by *Nsi*I restriction sites were amplified by PCR with primers listed in **Supplementary Table 8**. The amplified DNA fragment contains homologous arms to the target site that enables recombination. When introducing point mutations to *lpvA*, sites of mutagenesis were also introduced into the donor DNA by primers. The donor DNA and *lpv* were co-transformed into *E. coli* GB05-dir that contains an inducible RecET recombination system as described previously[5]. Colonies were picked and grown overnight in 5 mL LB medium supplemented with 50 µg/mL apramycin at 37 °C. The plasmid DNA was purified from the cultures using Qiaprep Spin Miniprep Kit (Qiagen, Germany), digested by appropriate restriction enzymes, and analyzed by agarose gel electrophoresis. Correct plasmids were digested by NsiI-HF and re-circularized by T4 DNA ligase to remove the ampicillin resistance marker. Each digestion reaction contains 100 ng plasmid, 0.5 µL NsiI-HF (10 U), and 2 µL rCutSmart Buffer within a total volume of 20 µL. The reaction was performed under 37 °C for 1 h, followed by 80 °C for 20 min to inactivate the enzyme. The reaction was then cooled to room temperature, supplemented by 0.2 µL (400 U) T4 DNA ligase, 0.8 µL of 25 mM ATP, and 1 µL of 200 mM DTT, and incubated at 20 °C for another 1 h.

The reaction mixture was then transformed into *E. coli* NEB10β. Plasmids were purified and analyzed by restriction enzyme digestion as described above. Correct plasmids were transformed into *E. coli* WM6026 and conjugated into *S. albus* J1074 for expression.

**Expression and purification of recombinant proteins.** Genes were PCR amplified from the genomic DNA of *Streptomyces* sp. NRRL S-1521 and *Streptomyces* sp. NRRL S-920 using primers listed in **Supplementary Table 7**, which were cloned into pET28a to produce proteins with *N*-terminal His-tag using NEBuilder HiFi DNA assembly master mix. Plasmids were transformed into *E. coli* BL21(DE3) for heterologous expression. Cells were grown overnight on LB agar plates (5 g/L yeast extract, 10 g/L tryptone, 10 g/L NaCl, 20 g/L agar) containing 50 μg/mL kanamycin at 37 °C. Single colonies were used to inoculate 5 mL of LB liquid medium containing 50 μg/mL kanamycin and grown at 37 °C for 14-18 h. This culture was used to inoculate 500 mL of Terrific Broth (TB; 24 g/L yeast extract, 20 g/L tryptone, 0.4% glycerol (*v/v*), 17 mM $KH_2PO_4$, and 72 mM $K_2HPO_4$) containing 50 μg/mL kanamycin and grown to an optical density at 600 nm ($OD_{600}$) of 0.6-0.8. Protein expression was induced by addition of isopropyl β-D-1-thiogalactopyranoside (IPTG) to a final concentration of 0.1 mM for 18 h at 18 °C. Cells were harvested by centrifugation and stored at -80 °C until further use.

Cell pellets were resuspended in 20 mL buffer A (50 mM $NaH_2PO_4$, 300 mM NaCl, 20 mM imidazole, pH 7.5) and treated with 20 mg lysozyme and 5 μL benzolase at 4 °C for 1 h. Cells were then lysed on ice by sonication using a Vibra Cell sonicator with the following settings: 60% amplitude, 5 min total sonication time; alternating between 2 s on/5 s off-pulse. Lysates were centrifuged at 4 °C for 30 min at 30,000 × g, and supernatants were filtered through a syringe filter (0.45 μm). Ni-NTA purification was performed using a 5 mL Ni-NTA HisTrap column (GE Healthcare). After loading, the column was washed with 50 mL of buffer A and eluted using a step gradient with an increasing percentage of buffer B (50 mM $NaH_2PO_4$, 300 mM NaCl, 500 mM imidazole, pH 7.5) in 15 mL volumes: 20%, 40%, 60%, 80% and 100% B. Fractions containing the recombinant protein were analyzed on SDS-PAGE and concentrated with Amicon centrifugal filter units (at suitable MW cut-offs) to around 2 mL. Concentrated proteins were loaded to PD-10 columns and eluted by storage buffer (50 mM Tris-HCl, 5 % glycerol (*v/v*), pH 7.5). Proteins were flash frozen by liquid nitrogen and stored at -80 °C until further use.

**Characterization of the *in vitro* activity of StsG.** The reaction of StsG contains 50 mM Tris-HCl (pH 7.5), 2 mM Tris (2-carboxyethyl) phosphine hydrochloride (TCEP-HCl), 1 mM isobutyryl-CoA (**4**), 1 mM methylmalonyl-CoA (**5**), and 50 μM StsG in a total volume of 20 μL. Boiled StsG was used as negative control. Either 2 mM **4** or **5** were used to set up reactions with substrate molar ratios of **4**:**5** at 2:1 or 1:2, respectively. When using the methylmalonyl-CoA epimerase, the enzyme was added to a final concentration of 50 μM. All reactions were performed at 30 °C for 10 min, quenched by adding 2 μL formic acid, stored at -20 °C, and diluted 5-fold by water right before analysis. A total of 5 μL of each sample was analyzed by HPLC using an Agilent 1260 system equipped with Phenomenex Kinetex SB-C18 column (5 μm, 100 Å, 250 × 10 mm) under the following conditions: mobile phase (A = 10 mM ammonium acetate, B = methanol), 5% to 100 % B over 15 min, 100% to 5% B for 1 min, 5% B for 15 min; flow rate 1.0 mL/min. The UV absorbance was monitored at 260 nm.

**Characterization of the *in vitro* activity of LpvV.** The two-enzyme reaction of StsG/LpvV was set up by adding 50 μM LpvV, along with either 1 mM NADH or 10 mM NADPH, to the StsG reaction in a total volume of 20 μL. Boiled LpvV was used as negative control. The reaction was performed at 30 °C for 10 min and quenched by adding 2 μL formic acid. Subsequent sample preparation and HPLC analysis followed the same methods as described for the StsG reactions.

**Preparation of 2,4-dimethyl-3-oxopentanoyl-CoA.** The 2,4-dimethyl-3-oxopentanoyl-CoA **10** was synthesized using a previously described carbonyldiimidazole (CDI)-activation method[6]. A mixture of CDI (4.2 mg, 26 μmol, 4 eq.) and 2,4-dimethyl-3-oxopentanoic acid (4.0 mg, 31 μmol, 4.8 eq.) in THF (200 μL) was stirred at 25 °C for 1 h. Afterwards, 5 mg CoA (6.4 μmol, 1 eq.) dissolved in 50 μL of 0.5 M $NaHCO_3$ was added to the reaction, which was stirred for another 45 min at 25 °C, flash frozen in liquid nitrogen, and lyophilized to dryness. The sample was then dissolved in 600 μL $H_2O$ and purified by HPLC using an Agilent 1260 system equipped with Phenomenex Kinetex SB-C18 column (5 μm, 100 Å, 150 × 10 mm) under the following conditions: mobile phase (A = 0.1 % *(v/v)* TFA in water, B = 0.1% (*v/v*) in acetonitrile); 5% to 35 % B over 15 min, 35% to 100% B over 3 min, 100% B for 3 min, 100% to 5% B over 1 min, 5% B for 3 min; flow rate 1.0 mL/min. The UV absorbance was monitored at 260 nm. The product was collected, analyzed by MALDI-TOF-MS, flash frozen in liquid nitrogen, lyophilized to dryness, and stored at -80 °C until further use.

**Characterization of the *in vitro* activity of LpvE.** The three-enzyme reaction of StsG, LpvV, and LpvE was set up by adding 50 μM LpvE and 0.5 mM peptide substrate (**2** or **3**) to the described two-enzyme reaction of StsG and LpvV in a total volume of 20 μL. Boiled LpvE was used as negative control. The reaction was performed at 30 °C for 20 h and quenched by adding 2 μL formic acid. The reaction mixture was then diluted 5-fold by water and analyzed by HPLC using an Agilent 1260 system equipped with Phenomenex Kinetex SB-C18 column (5 μm, 100 Å, 250 × 10 mm) under the following conditions: mobile phase (A = 0.1 % (*v/v*) TFA in water, B = 0.1% (*v/v*) in acetonitrile); 30% to 70 % B over 15 min, 70% to 100% B for 3 min, 100% B for 5 min, 100% to 5% B for 1min, 5% B for 5 min; flow rate 1.0 mL/min. The UV absorbance was monitored at 280 nm. The reactions of StsE were performed and analyzed using the same method as described for LpvE. The reaction of LpvE using 2,4-dimethyl-3-oxopentanoyl-CoA as the acyl donor was assayed in 50 mM Tris-HCl (pH 7.5), 2 mM TCEP-HCl, 1 mM 2,4-dimethyl-3-oxopentanoyl-CoA, and 1 mM peptide substrate (**2** or **3**). Subsequent sample preparation and HPLC analysis methods were same as described for the three-enzyme reaction.

**Structure simulation molecular docking of LpvE.** The structures of **2** and **9** were optimized with semi-empirical quantum mechanics MOPAC[7] and energy minimization. AlphaFold2 was used for predicting the three-dimensional structure of LpvE, with each of the five trained model parameters from CASP14 models[8]. The multiple sequence alignment (MSA) generation, AlphaFold predictions, and Amber structural relaxation were run on local server with GPU. A full database (updated to 2021-09-17) was applied for the structure predictions. For molecular docking, a grid box of 5 Å was created around the active site within LpvE. The docking study of 2 was performed after obtaining the LpvE-9 docking complex. Autodock 4.2 plug-in[9] within YASARA (version 20.4.24)[10, 11] was used to perform molecular docking calculations with a fixed protein backbone. The protein residues were treated with AMBER ff99 force field[12], and the ligand atoms were treated using GAFF[13, 14] with AM1-BCC partial charges[15]. A total of 100 docking runs were calculated, and the obtained docking poses were clustered using an RMSD cutoff of 0.5 Å within the YASARA dock_run macro file. Finally, ChimeraX (version 1.4)[16] was used to visualize and analyze the proteins.

**Supplementary Table 1: Pfam annotation of *lpv* proteins.** The top match for each protein is given. Proteins lacking a match are indicated by "N/A". LpvP2 was also analyzed by HHpred[17] and Foldseek[18], with results in **Supplementary Fig. 2**, showing its potential function as a peptidase.

| Name | Accession Code | Protein Family | Functional Annotation |
|---|---|---|---|
| LpvR1 | WP_062773463.1 | PF00072 | Response regulator receiver domain |
| | | PF00196 | Bacterial regulatory proteins, luxR family |
| | | PF04545 | Sigma-70, region 4 |
| | | PF08281 | Sigma-70, region 4 |
| | | PF13384 | Homeodomain-like domain |
| LpvR2 | WP_062773466.1 | PF07730 | Histidine kinase |
| LpvP1 | WP_062773469.1 | PF02517 | CPBP intramembrane metalloprotease |
| LpvV | WP_062773474.1 | PF00106 | short chain dehydrogenase |
| | | PF13561 | Enoyl-(Acyl carrier protein) reductase |
| | | PF08659 | KR domain |
| LpvU3 | WP_062773479.1 | N/A | N/A |
| LpvP2 | WP_062773482.1 | N/A | N/A |
| LpvR3 | WP_062773485.1 | PF01381 | Helix-turn-helix |
| | | PF13560 | Helix-turn-helix domain |
| LpvR4 | WP_079107248.1 | PF00392 | Bacterial regulatory proteins, gntR family |
| | | PF13730 | Helix-turn-helix domain |
| | | PF02082 | Transcriptional regulator |
| | | PF08222 | CodY helix-turn-helix domain |
| LpvA | WP_159043817.1 | N/A | N/A |
| LpvB | WP_062773494.1 | PF01636 | Phosphotransferase enzyme family |
| LpvC | WP_062773498.1 | PF17914 | HopA1 effector protein family |
| LpvD | WP_062773501.1 | PF02441 | Flavoprotein |
| LpvE | WP_136241767.1 | PF13302 | Acetyltransferase (GNAT) domain |
| | | PF13523 | Acetyltransferase (GNAT) domain |
| LpvF | WP_062773508.1 | PF07366 | SnoaL-like polyketide cyclase |
| | | PF12680 | SnoaL-like domain |
| LpvG | WP_062773511.1 | PF08541 | 3-Oxoacyl-[acyl-carrier-protein (ACP)] synthase III *C* terminal |
| | | PF08545 | 3-Oxoacyl-[acyl-carrier-protein (ACP)] synthase III |
| LpvH | WP_159043818.1 | PF00107 | Zinc-binding dehydrogenase |
| LpvM1 | WP_062773518.1 | PF13649 | Methyltransferase domain |
| | | PF08241 | Methyltransferase domain |
| | | PF13847 | Methyltransferase domain |
| | | PF08242 | Methyltransferase domain |
| | | PF13489 | Methyltransferase domain |
| LpvU1 | WP_062773521.1 | N/A | N/A |
| LpvI | WP_062773525.1 | PF06722 | Protein of unknown function (DUF1205) |
| | | PF00201 | UDP-glucoronosyl and UDP-glucosyl transferase |
| | | PF04101 | Glycosyltransferase family 28 C-terminal domain |
| LpvK | WP_062773528.1 | PF05721 | Phytanoyl-CoA dioxygenase (PhyH) |
| LpvJ | WP_062773532.1 | PF00296 | Luciferase-like monooxygenase |
| LpvU2 | WP_062773536.1 | N/A | N/A |
| LpvL | WP_062773540.1 | PF00483 | Nucleotidyl transferase |
| | | PF12804 | MobA-like NTP transferase domain |
| LpvN | WP_062773543.1 | PF16363 | GDP-mannose 4,6 dehydratase |
| | | PF01370 | NAD dependent epimerase/dehydratase family |
| | | PF01073 | 3-beta hydroxysteroid dehydrogenase/isomerase family |

| | | | |
|---|---|---|---|
| | | PF02719 | Polysaccharide biosynthesis protein |
| | | PF04321 | RmlD substrate binding domain |
| LpvO | WP_063802748.1 | PF04321 | RmlD substrate binding domain |
| | | PF01370 | NAD dependent epimerase/dehydratase family |
| | | PF16363 | GDP-mannose 4,6 dehydratase |
| | | PF01073 | 3-beta hydroxysteroid dehydrogenase/isomerase family |
| | | PF02719 | Polysaccharide biosynthesis protein |
| LpvQ | WP_062773549.1 | PF00908 | dTDP-4-dehydrorhamnose 3,5-epimerase |
| LpvM2 | WP_159043819.1 | PF17843 | MycE methyltransferase *N*-terminal |
| | | PF13578 | Methyltransferase domain |
| LpvS | WP_062773556.1 | PF00067 | Cytochrome P450 |

**Supplementary Table 2: InterPro annotation of *lpv* proteins.** Proteins lacking a match are indicated by "N/A".

| Name | Accession Code | InterPro Family | Functional Annotation |
|---|---|---|---|
| LpvR1 | WP_062773463.1 | IPR001789 | Signal transduction response regulator, receiver domain |
| | | IPR000792 | Transcription regulator LuxR, C-terminal |
| | | IPR016032 | Signal transduction response regulator, C-terminal effector |
| | | IPR039420 | Transcriptional regulatory protein WalR-like |
| | | IPR011006 | CheY-like superfamily |
| LpvR2 | WP_062773466.1 | IPR036890 | Histidine kinase/HSP90-like ATPase superfamily |
| | | IPR011712 | Signal transduction histidine kinase, subgroup 3, dimerisation and phosphoacceptor domain |
| LpvP1 | WP_062773469.1 | IPR003675 | Type II CAAX prenyl endopeptidase Rce1-like |
| LpvV | WP_062773474.1 | IPR020904 | Short-chain dehydrogenase/reductase, conserved site |
| | | IPR002347 | Short-chain dehydrogenase/reductase SDR |
| | | IPR036291 | NAD(P)-binding domain superfamily |
| LpvU3 | WP_062773479.1 | IPR046675 | Protein of unknown function DUF6545 |
| LpvP2 | WP_062773482.1 | N/A | N/A |
| LpvR3 | WP_062773485.1 | IPR001387 | Cro/C1-type helix-turn-helix domain |
| | | IPR010982 | Lambda repressor-like, DNA-binding domain superfamily |
| LpvR4 | WP_079107248.1 | IPR000524 | Transcription regulator HTH, GntR |
| | | IPR036388 | Winged helix-like DNA-binding domain superfamily |
| | | IPR036390 | Winged helix DNA-binding domain superfamily |
| LpvA | WP_159043817.1 | N/A | N/A |
| LpvB | WP_062773494.1 | IPR011009 | Protein kinase-like domain superfamily |
| | | IPR002575 | Aminoglycoside phosphotransferase |
| LpvC | WP_062773498.1 | IPR040871 | HopA1 effector protein |
| LpvD | WP_062773501.1 | IPR003382 | Flavoprotein |
| | | IPR036551 | Flavin prenyltransferase-like |
| LpvE | WP_136241767.1 | IPR016181 | Acyl-CoA N-acyltransferase |
| | | IPR000182 | GNAT domain |
| LpvF | WP_062773508.1 | IPR009959 | Polyketide cyclase SnoaL-like |
| | | IPR032710 | NTF2-like domain superfamily |
| LpvG | WP_062773511.1 | IPR016039 | Thiolase-like |
| | | IPR013747 | Beta-ketoacyl-[acyl-carrier-protein] synthase III, C-terminal |
| | | IPR013751 | Beta-ketoacyl-[acyl-carrier-protein] synthase III, N-terminal |
| LpvH | WP_159043818.1 | IPR020843 | Polyketide synthase, enoylreductase domain |
| | | IPR036291 | NAD(P)-binding domain superfamily |
| | | IPR011032 | GroES-like superfamily |
| | | IPR013149 | Alcohol dehydrogenase-like, C-terminal |
| LpvM1 | WP_062773518.1 | IPR029063 | S-adenosyl-L-methionine-dependent methyltransferase superfamily |
| | | IPR041698 | Methyltransferase domain 25 |
| LpvU1 | WP_062773521.1 | IPR046732 | Protein of unknown function DUF6624 |
| LpvI | WP_062773525.1 | IPR002213 | UDP-glucuronosyl/UDP-glucosyltransferase |
| | | IPR010610 | Erythromycin biosynthesis protein CIII-like, central |
| LpvK | WP_062773528.1 | IPR008775 | Phytanoyl-CoA dioxygenase-like |
| LpvJ | WP_062773532.1 | IPR011251 | Luciferase-like domain |
| | | IPR036661 | Luciferase-like domain superfamily |
| | | IPR025660 | Cysteine peptidase, histidine active site |
| LpvU2 | WP_062773536.1 | N/A | N/A |
| LpvL | WP_062773540.1 | IPR005835 | Nucleotidyl transferase domain |
| | | IPR005908 | Glucose-1-phosphate thymidylyltransferase, long form |

| | | | |
|---|---|---|---|
| | | IPR029044 | Nucleotide-diphospho-sugar transferases |
| LpvN | WP_062773543.1 | IPR005888 | dTDP-glucose 4,6-dehydratase |
| | | IPR016040 | NAD(P)-binding domain |
| | | IPR036291 | NAD(P)-binding domain superfamily |
| | | IPR020904 | Short-chain dehydrogenase/reductase, conserved site |
| LpvO | WP_063802748.1 | IPR029903 | RmlD-like substrate binding domain |
| | | IPR005913 | dTDP-4-dehydrorhamnose reductase family |
| | | IPR036291 | NAD(P)-binding domain superfamily |
| LpvQ | WP_062773549.1 | IPR000888 | dTDP-4-dehydrorhamnose 3,5-epimerase-related |
| | | IPR014710 | RmlC-like jelly roll fold |
| | | IPR011051 | RmlC-like cupin domain superfamily |
| LpvM2 | WP_159043819.1 | IPR029063 | S-adenosyl-L-methionine-dependent methyltransferase superfamily |
| | | IPR040800 | Methyltransferase MycE, N-terminal |
| LpvS | WP_062773556.1 | IPR036396 | Cytochrome P450 superfamily |
| | | IPR017972 | Cytochrome P450, conserved site |
| | | IPR002397 | Cytochrome P450, B-class |
| | | IPR001128 | Cytochrome P450 |

**Supplementary Table 3: BLAST-P results of LpvS in the SwissProt database.** The top 10 hits are listed below.

| Description | Scientific Name | Max Score | Total Score | Query Cover | E value | Per. ident | Accession |
|---|---|---|---|---|---|---|---|
| Mycinamicin IV hydroxylase/epoxidase; Cytochrome P450 MycG; Multifunctional P450 enzyme; Mycinamicin biosynthesis protein G | *Micromonospora griseorubida* | 253 | 253 | 94% | 2E-79 | 38.34 | Q59523.1 |
| Cytochrome P450 107B1; Cytochrome P450CVIIB1 | *Saccharopolyspora erythraea* NRRL 2338 | 251 | 251 | 98% | 2E-78 | 38.05 | P33271.1 |
| Vitamin D(3) 25-hydroxylase; Cytochrome P450 | *Pseudonocardia autotrophica* | 247 | 247 | 98% | 6E-77 | 38.14 | C4B644.1 |
| Polyketide biosynthesis cytochrome P450 PksS | *Bacillus subtilis* subsp. *subtilis* str. 168 | 245 | 245 | 97% | 2E-76 | 33.08 | O31785.2 |
| Cytochrome P450 CYP107DY1; Mevastatin hydroxylase | *Priestia megaterium* QM B1551 | 245 | 245 | 89% | 2E-76 | 37.37 | D5E3H2.1 |
| Cytochrome P450 monooxygenase PikC; Cytochrome P450 monooxygenase PicK; Narbomycin C-12 hydroxylase; Pikromycin synthase CYP107L1 | *Streptomyces venezuelae* | 244 | 244 | 99% | 1E-75 | 38.63 | O87605.1 |
| 6-deoxyerythronolide B hydroxylase; 6-DEB hydroxylase; CYPCVIIA1; Cytochrome P450 107A1; Cytochrome P450eryF; Erythromycin A biosynthesis hydroxylase | *Saccharopolyspora erythraea* NRRL 2338 | 235 | 235 | 96% | 1E-72 | 36.95 | Q00441.2 |
| Nocardicin C *N*-oxygenase | *Nocardia uniformis* subsp. *tsuyamanensis* | 221 | 221 | 95% | 5E-67 | 37.12 | Q5J1R4.1 |
| Cytochrome P450 | *Bacillus subtilis* subsp. *subtilis* str. 168 | 213 | 213 | 99% | 5E-64 | 30.66 | O08469.1 |
| Vitamin D3 dihydroxylase; CYP105A1; Cytochrome P450-CVA1; Cytochrome P450-SU1; Vitamin D3 hydroxylase | *Streptomyces griseolus* | 204 | 204 | 94% | 1E-60 | 34.79 | P18326.2 |

**Supplementary Table 4: Representative thioamitides and the corresponding core peptide sequences**. The *N*-terminal pyruvyl moieties in thioholgamide A[19] and thiostreptamide S4[20] are derived from serine dehydration, followed by spontaneous tautomerization after leader peptide removal. Further reduction of the pyruvyl ketone results in the lactyl moieties found in thioalbamide[20].

| Names | Structures | Core peptide sequences |
|---|---|---|
| Prethioviridamide | 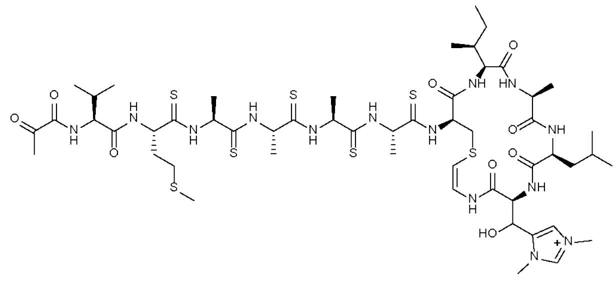 | SVMAAAASIALHC |
| Thioholgamide A | 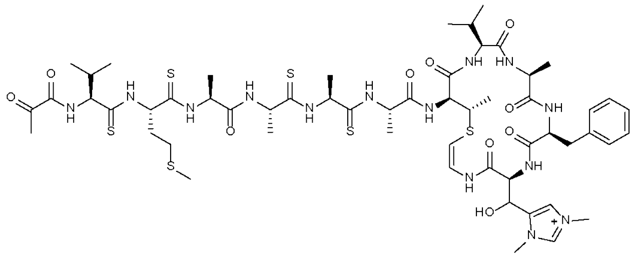 | SVMAAAATVAFHC |
| Thioalbamide | 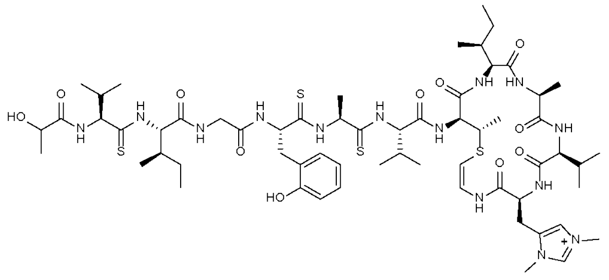 | SVIGFAVTIAVHC |
| Thiostreptamide S4 | 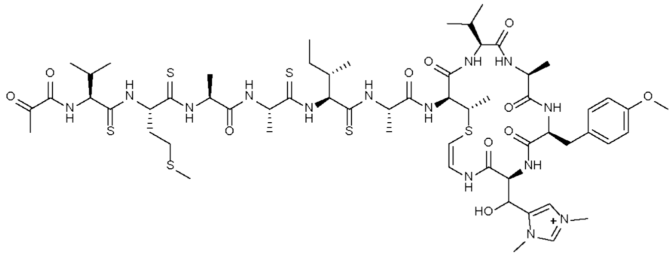 | SVMAAIATVAYHC |

**Supplementary Table 5: $^{1}$H NMR (600 MHz) and $^{13}$C NMR (150 MHz) spectroscopic data for 1**. Data was obtained in acetone acetone-$d_6$ (δ in ppm).

| Residue | $\delta_C$, type | $\delta_H$, multi ($J$ in Hz) | Residue | $\delta_C$, type | $\delta_H$, multi ($J$ in Hz) |
|---|---|---|---|---|---|
| HMP | | | 2 | 62.0, CH | 4.23, m |
| 1 | 180.0, C | | 3 | 36.5, CH | 3.20, overlapped |
| 2 | 36.6, CH | 2.87, m | 4 | 131.5, C | |
| 3 | 47.7, $CH_2$ | 1.31, overlapped | 5/9 | 130.8, CH | 7.29, d (8.5) |
| | | 2.29, m | 6/8 | 117.2, $CH_3$ | 6.93, d (8.5) |
| 4 | 70.7, C | | 7 | 157.6, C | |
| 5 | 27.3, $CH_3$ | 1.31, s | NH | | 8.85, d (3.9) |
| 6 | 21.0, $CH_3$ | 1.13, d (6.8) | 2-*O*-methyl-β-6- | | |
| 7 | 32.4, $CH_3$ | 1.21, s | deoxygulose | | |
| Val | | | 1′ | 99.6, CH | 5.25, d (8.0) |
| 1 | 176.3, C | | 2′ | 78.4, CH | 3.47, dd (8.0, 3.1) |
| 2 | 65.4, CH | 3.60, dd (9.7, 4.8) | 3′ | 71.0, CH | 4.16, dd (3.3, 3.1) |
| 3 | 30.1, CH | 2.05, m | 4′ | 72.5, CH | 3.55, br d (3.3) |
| 4 | 20.9, $CH_3$ | 1.10, d (6.4) | 5′ | 69.7, CH | 4.13, m |
| 5 | 19.6, $CH_3$ | 0.97, d (6.7) | 6′ | 16.3, $CH_3$ | 1.19, d (6.5) |
| NH | | 9.04, d (4.8) | 2′-OMe | 58.7, $CH_3$ | 3.47, s |
| Gly | | | AviMeCys | | |
| 1 | 172.4, C | | 1 | 172.7, C | |
| 2 | 46.3, $CH_2$ | 3.63, dd (15.0, | 2 | 58.6, CH | 4.04, overlapped |
| | | 4.3) | 3 | 49.3, CH | 3.05, m |
| | | 3.72, dd (15.0, | 4 | 23.8, $CH_3$ | 1.51, overlapped |
| | | 6.5) | 5 | 101.8, CH | 5.61, d (6.2) |
| NH | | 9.01, t (5.2) | 6 | 133.7, CH | 7.29, overlapped |
| *D*-Ala | | | 2-NH | | 8.75, d (7.2) |
| 1 | 174.5, C | | 6-NH | | 10.70, d (10.8) |
| 2 | 49.8, CH | 3.93, m | Phe | | |
| 3 | 15.3, $CH_3$ | 1.50, overlapped | 1 | 175.0, C | |
| NH | | 9.08, d (7.0) | 2 | 60.4, CH | 4.36, m |
| Ile 1 | | | 3 | 38.8, $CH_2$ | 3.19, overlapped |
| 1 | 175.8, C | | 4 | 138.7, C | |
| 2 | 64.1, CH | 3.68, dd (10.0, | 5/9 | 130.2, CH | 7.39, d (7.4) |
| | | 4.1) | 6/8 | 129.2, CH | 7.28, overlapped |
| 3 | 36.5, CH | 2.14, m | 7 | 127.4, CH | 7.22, dd (7.2) |
| 4 | 27.6, $CH_2$ | 1.12, m | NH | | 8.37, d (3.6) |
| | | 2.00, m | Met | | |
| 5 | 11.6, $CH_3$ | 0.87, t (7.1) | 1 | 174.1, C | |
| 6 | 15.9, $CH_3$ | 0.99, d (6.7) | 2 | 57.4, CH | 4.06, overlapped |
| NH | | 8.57, d (4.1) | 3 | 31.7, $CH_2$ | 2.02, m |
| Ile 2 | | | | | 2.24, m |
| 1 | 176.1, C | | 4 | 30.9, $CH_2$ | 2.68, m |
| 2 | 64.5, CH | 3.81, dd (10.4, | | | 2.81, m |
| | | 4.4) | *S*-Me | 14.9, $CH_3$ | 2.03, s |
| 3 | 35.8, CH | 2.07, m | NH | | 8.73, br s |
| 4 | 27.5, $CH_2$ | 1.17, m | Ile 3 | | |
| | | 1.99, m | 1 | 172.9, C | |
| 5 | 11.6, $CH_3$ | 0.85, t (7.4) | 2 | 57.3, CH | 4.11, overlapped |
| 6 | 15.6, $CH_3$ | 0.93, d (6.4) | 3 | 36.1, CH | 2.03, m |
| NH | | 8.61, d (2.6) | 4 | 26.2, $CH_2$ | 1.08, m |
| Tyr | | | 5 | 11.8, $CH_3$ | 0.61, t (7.3) |
| 1 | 174.6, C | | 6 | 16.3, $CH_3$ | 0.72, d (6.8) |

| Residue | $\delta_C$, type | $\delta_H$, multi ($J$ in Hz) | Residue | $\delta_C$, type | $\delta_H$, multi ($J$ in Hz) |
|---|---|---|---|---|---|
| NH | | 8.01, d (8.0) | 4 | 134.4, C | |
| hdmHis | | | 5 | 123.8, CH | 7.77, s |
| CO | 168.8, C | | $N^1$-Me | 36.4, $CH_3$ | 3.91, s |
| α | 58.1 | 4.14, overlapped | $N^3$-Me | 35.1, $CH_3$ | 4.04, s |
| β | 63.1, CH | 5.93, d (8.8) | NH | | 7.52, d (7.0) |
| 2 | 138.0, CH | 8.80, s | | | |

**Supplementary Table 6: Indicator strains and corresponding culturing conditions used in the antimicrobial activity screening.** Recipes for media used in the bioassay are as the following: lysogeny broth (LB): 10 g/L tryptone, 10 g/L NaCl, and 5 g/L yeast extract; brain heart infusion broth (BHI): 5 g/L beef heart (infusion from 250 g), 12.5 g/L calf brains (infusion from 200 g), 2.5 g/L $Na_2HPO_4$, 2 g/L glucose, 10 g/L peptone, and 5 g/L NaCl; international *Streptomyces* project-2 medium (ISP2): 10 g/L malt extract, 4 g/L yeast extract, and 4 g/L glucose; ATCC medium 172 (ATCC 172): 10 g/L glucose, 20 g/L soluble starch, 5 g/L yeast extract, 5 g/L N-Z Amine Type A (Sigma C0626), and 1 g/L $CaCO_3$; yeast extract–peptone–dextrose medium (YPD): 10 g/L yeast extract, 20 g/L peptone, and 20 g/L glucose.

| Phylum | Strain | Medium | Temp (°C) |
|---|---|---|---|
| Bacillota | *Bacillus cereus* TZ417 | LB | 30 |
| | *Bacillus subtilis* ATCC 6633 | LB | 30 |
| | *Lactococcus lactis* CNRZ 481 | LB | 30 |
| | *Staphylococcus epidermidis* 15X154 | LB | 37 |
| | *Staphylococcus aureus* USA300 | BHI | 37 |
| | *Streptococcus mutans* ATCC 25175 | BHI | 37 |
| Actinomycetota | *Micrococcus luteus* ATCC 4698 | LB | 30 |
| | *Streptomyces albus* J1074 | ISP2 | 30 |
| | *Streptomyces lividans* TK24 | ISP2 | 30 |
| | *Streptomyces coelicolor* M145 | ISP2 | 30 |
| | *Microbacterium aurum* B-24210 | ATCC 172 | 30 |
| | *Microbacterium esteraromaticum* B-24213 | ATCC 172 | 30 |
| | *Microbacterium hominis* B-24220 | ATCC 172 | 30 |
| | *Microbacterium ketosireducens* B-24221 | ATCC 172 | 30 |
| | *Microbacterium kitamiense* B-24226 | ATCC 172 | 30 |
| Pseudomonadota | *Escherichia coli* DH5a | LB | 37 |
| | *Enterobacter cloacae* | LB | 30 |
| | *Pseudomonas fluorescens* Pf-5 | LB | 30 |
| | *Pseudomonas putida* mt-2 | LB | 30 |
| Ascomycota | *Saccharomyces cerevisiae* YSG50 | YPD | 30 |
| | *Aspergillus terreus* | YPD | 30 |

**Supplementary Table 7: $^1$H NMR (600 MHz) spectroscopic data for the HMP moiety of 10.** Data were obtained in acetone acetone-$d_6$ (δ in ppm).

| Residue | $\delta_H$, multi ($J$ in Hz) |
|---|---|
| HMP | |
| 1 | |
| 2 | 2.83, overlapped |
| 3 | 3.62, m |
| 4 | 1.87, m |
| 5 | 1.03, d (6.8) |
| 6 | 1.07, d (6.3) |
| 7 | 0.85, overlapped |

**Supplementary Table 8: Accession codes of GNAT acyltransferases used for phylogenetic analysis**. Strains sharing the same accession code are listed accordingly.

| Class | Name | Accession code | Strain |
|---|---|---|---|
| GNAT | PA4794 | WP_003111411.1 | *Pseudomonas aeruginosa* |
| | ApxC | WP_009875272.1 | *Actinobacillus pleuropneumoniae* |
| | GAT | WP_003180517.1 | *Bacillus licheniformis* |
| | CBG | WP_003952521.1 | *Streptomyces clavuligerus* |
| | MccE | WP_021528069.1 | *Escherichia coli* |
| | PseH | WP_120818319.1 | *Helicobacter pylori* |
| | WecD | WP_001145166.1 | *Escherichia coli* |
| | MddA | WP_129406907.1 | *Salmonella enterica* |
| | MbtK | WP_003406956.1 | *Mycobacterium tuberculosis* |
| | PhnO | WP_000602316.1 | *Salmonella enterica* |
| | Rv0802c | WP_003404111.1 | *Mycobacterium tuberculosis* |
| | GlmA | WP_010963508.1 | *Clostridium acetobutylicum* |
| | MtAAC(21)-Ic | WP_003899880.1 | *Mycobacterium tuberculosis* |
| | SmAAC(3)-Ia | AAB20441.1 | *Serratia marcescens* |
| | EcAAC(61)-Ib | WP_001749987.1 | *Escherichia coli* |
| | SeAAC(61)-Ib11 | WP_032490438.1 | *Salmonella typhimurium* |
| | SwAAC(61)-Ie | WP_001028144.1 | *Staphylococcus warneri* |
| | EfAAC(61)-Ii | WP_008265821.1 | *Enterococcus faecium* |
| | SeAAC(61)-Iy | WP_000354853.1 | *Salmonella enteritidis* |
| | MtEis | WP_003903886.1 | *Mycobacterium tuberculosis* |
| | hHAT1 | NP_003633.1 | *Homo sapiens* |
| | ScHAT1 | NP_015324.1 | *Saccharomyces cerevisiae* |
| | ScEsa1 | NP_014887.3 | *Saccharomyces cerevisiae* |
| | ScGCN5 | NP_011768.1 | *Saccharomyces cerevisiae* |
| | TtGCN5 | AAB01099.1 | *Tetrahymena thermophila* |
| | hGCN5 | NP_066564.2 | *Homo sapiens* |
| | hPCAF | NP_003875.3 | *Homo sapiens* |
| | ScHpa2 | NP_015519.1 | *Saccharomyces cerevisiae* |
| | aTAT1 | NP_001026892.1 | *Homo sapiens* |
| | MtPAT | WP_003405164.1 | *Mycobacterium tuberculosis* |
| | SsPAT | WP_009989035.1 | *Saccharolobus solfataricus* |
| | hNaa50p | NP_079422.1 | *Homo sapiens* |
| | OaAANAT | NP_001009461.1 | *Ovis aries* |
| | DmAANAT | NP_523839.2 | *Drosophila melanogaster* |
| | AaAANAT | XP_001663122.1 | *Aedes aegypti* |
| | ScGNA1 | NP_116637.1 | *Saccharomyces cerevisiae* |
| | AfGNA1 | XP_747831.1 | *Neosartorya fumigata* |
| | hGNA1 | NP_932332.1 | *Homo sapiens* |

| | | | |
|---|---|---|---|
| | TbGNA1 | XP_829175.1 | *Trypanosoma brucei brucei* |
| | AtGNA1 | NP_197081.1 | *Arabidopsis thaliana* |
| | CeGNA1 | NP_505654.1 | *Caenorhabditis elegans* |
| | HpPseH | WP_000742697.1 | *Helicobacter pylori* |
| | CjPseH | AFU43358.1 | *Campylobacter jejuni* subsp. *jejuni* PT14 |
| | EcWecD | WP_001145189.1 | *Escherichia coli* O6:H1 |
| | PsTTR | WP_002555402.1 | *Pseudomonas amygdali pv. tabaci* |
| | ScMpr1 | BAA95611.1 | *Saccharomyces cerevisiae* |
| | BsPaiA | WP_003244000.1 | *Bacillus subtilis* |
| | TaPaiA | WP_010900802.1 | *Thermoplasma acidophilum* |
| | hSSAT | NP_001307775.1 | *Homo sapiens* |
| | MmSSAT | NP_001278794.1 | *Mus musculus* |
| | VcSpeG | WP_001088091.1 | *Vibrio cholerae* serotype O1 |
| | SeRimI | WP_001092436.1 | *Salmonella typhimurium* |
| | StRimL | WP_000229304.1 | *Salmonella typhimurium* |
| | BsYadF | WP_003246691.1 | *Bacillus subtilis* |
| | SaFemA | WP_000673309.1 | *Staphylococcus aureus* |
| | WvFemX | WP_057745451.1 | *Weissella viridescens* |
| | EcLFT | WP_001241678.1 | *Escherichia coli* |
| | MtMshD | WP_003404307.1 | *Mycobacterium tuberculosis* |
| ***N*-myristoyltransferase** | CaNMT | XP_722713.1 | *Candida albicans* |
| | LdNMT | XP_001467690.1 | *Leishmania infantum* |
| | LmNMT | XP_001685320.1 | *Leishmania major* |
| | ScNMT | NP_013296.1 | *Saccharomyces cerevisiae* |
| | PvNMT | XP_001616826.1 | *Plasmodium vivax* |
| | AfNMT | XP_752019.1 | *Aspergillus fumigatus* |
| | hNMT | NP_004799.1 | *Homo sapiens* |
| **Goadvionin Type A** | SosG | WP_123543502.1 | *Streptomyces ossamyceticus* SAI-001 |
| | SjvG | WP_099964717.1 | *Streptomyces* sp. JV178 |
| | AalG | WP_113692540.1 | *Amycolatopsis albispora* WP1 |
| | AprG | WP_118947408.1 | *Actinosynnema pretiosum* subsp. *pretiosum* |
| | SpaG | WP_080681602.1 | *Salinispora pacifica* CNT08 |
| | AmeG | WP_089330376.1 | *Actinomadura meyerae* DSM 44715 |
| | SfdG | WP_076046519 .1 | *Streptomyces* sp. fd1-xmd |
| | GdvG | BBK08001.1 | *Streptomyces sp. TP-A0584* |
| **Goadvionin Type B** | SalG | WP_027648441.1 | *Salinispora pacifica* CNT001 |
| | SaiG | WP_018221346.1 | *Salinispora pacifica* CNQ768 |
| | SapG | WP_027657831.1 | *Salinispora pacifica* CNR894 |
| **Goadvionin Type C** | SacG | WP_018218316.1 | *Salinispora pacifica* DSM 45548 = CNT-148 |
| **Class I lipolanthin** | FlaB | WP_030318155.1 | *Streptomyces flavochromogenes* |
| | ChrB | WP_017622812.1 | *Nocardiopsis chromatogenes* YIM 90109 |

| | | | |
|---|---|---|---|
| | NatB | WP_030066285.1 | *Streptomyces natalensis* |
| | AngB | WP_263296008.1 | *Streptomyces angustmyceticus* |
| | KatB | WP_228386208.1 | *Streptomyces katsurahamanus* strain T-272 |
| | JumB | WP_228387932.1 | *Streptomyces jumonjinensis* strain NRRL 5741 |
| **Class II lipolanthin** | NocB | KZM69140.1 | *Nocardia terpenica* strain IFM406 |
| | NoxB | WP_098692972.1 | *Nocardia terpenica* strain NC_YFY_NT001 |
| | ClaB | WP_104293653.1 | *Clavibacter michiganensis* strain Z001 |
| | TsuB | WP_148281472.1 | *Tsukamurella* sp. 1534 |
| | NrhB | WP_051823113.1 | *Streptomyces* sp. NRRL S-1448 |
| | NrgB | WP_051798843.1 | *Streptomyces* sp. NRRL S-337 |
| **Class III lipolanthin** | DzeB | WP_053172880.1 | *Streptomyces* sp. 3211 |
| | FssB | WP_030764502.1 | *Streptomyces* sp. NRRL F-2664 |
| | HodB | WP_051735155.1 | *Streptomyces* sp. H036 |
| | IgbB | WP_053632111.1 | *Streptomyces* sp. IGB124 |
| | NffB | WP_045321850.1 | *Streptomyces* sp. NRRL F-4428 |
| | NrsB | WP_051779408.1 | *Streptomyces* sp. NRRL S-241 |
| | RibB | WP_050501170.1 | *Streptomyces rimosus* subsp. *rimosus* strain NRRL WC-3927 |
| | TnfB | WP_075970905.1 | *Streptomyces* sp. TN58 |
| | CfbB | WP_086527277.1 | *Clavibacter michiganensis* strain CFBP7576 |
| | SafB | WP_093416179.1 | *Saccharopolyspora flava* DSM 44771 |
| | LydB | WP_078617180.1 | *Streptomyces lydicus* NRRL ISP-5461 |
| | MoeB | WP_084771993.1 | *Streptomyces* sp. MOE7 |
| | KseB | WP_176956483.1 | *Streptomyces* sp. KS_16 |
| | SpzB | WP_093492296.1 | *Streptomyces* sp. 2112.3 |
| | XinB | WP_095756843.1 | *Streptomyces xinghaiensis* S187 |
| | NrrB | WP_051851820.1 | *Streptomyces sp.* NRRL F-5650 |
| | RocB | ANW61951.1 | *Streptomyces rochei* strain Sal35 |
| | KutB | WP_043715796.1 | *Kutzneria* sp. 744 |
| | AmjB | WP_084145360.1 | *Amycolatopsis jejuensis* NRRL B-24427 |
| | DecB | WP_048828641.1 | *Streptomyces decoyicus* NRRL 2666 |
| | CtdB | WP_052230000.1 | *Streptomyces* sp. CT34 |
| | NrxB | WP_051818380.1 | *Streptomyces* sp. NRRL S-1813 |
| | AlbB | WP_060733178.1 | *Streptomyces albus* subsp. *Albus* NRRL F-4371 |
| | FfsB | WP_053700687.1 | *Streptomyces* sp. NRRL F-5755 |
| | GriB | WP_050507140.1 | *Streptomyces griseoflavus* NRRL B-1830 |
| | PeuB | WP_050508296.1 | *Streptomyces peucetius* NRRL WC-3868 |
| | RimB | WP_050515185.1 | *Streptomyces rimosus* subsp. *rimosus* NRRL WC-3904 |
| **Class IV lipolanthin** | AmrB | WP_076046519.1 | *Streptomyces amritsarensis* MTCC 11845 |
| | ImtB | WP_060180521.1 | *Streptomyces* sp. IMTB 1903 |
| | AmyB | WP_162788449.1 | *Amycolatopsis albispora* WP1 |
| | NeyB | WP_055538078.1 | *Streptomyces neyagawaensis* NRRL B-3092 |

| | | | |
|---|---|---|---|
| | PrcB | WP_074995603.1 | *Streptomyces prasinopilosus* CGMCC 4.3504 |
| **Unique lipolanthin** | RuxB | WP_030361744.1 | *Streptomyces ruber* NRRL ISP-5378 |
| | VwiB | TQJ31121.1 | *Microbacterium* sp. SLBN-146 |
| **Selidamide** | KspN | WP_007358511.1 | *Kamptonema formosum* PCC 6407 |
| | NpuN | WP_012412981.1 | *Nostoc punctiforme* PCC 73102 |
| | PhaN | WP_084300771.1 | *Pseudophaeobacter arcticus* |
| | MaeN | KXS90083.1 | *Microcystis aeruginosa* NIES-88 |
| | NosN | BAY36667.1 | *Nostoc* sp. NIES-2111 |
| | ScyN | WP_017746172.1 | *Scytonema hofmannii* PCC 7110 |
| | NspN | BAZ52079.1 | *Nostoc* sp. NIES-4103 |
| | SnoN | PZV23233.1 | *Snowella* sp. isolate ULC335bin1 |
| | ChaN | PSB57665.1 | *Chamaesiphon polymorphus* CCALA 037 |
| | DbaN | PID77288.1 | *Deltaproteobacteria bacterium* isolate DOLZORAL124_49_6 |
| | PruN | KGE03277.1 | *Pseudohaliea rubra* DSM 19751 |
| | WabN | HBC75360.1 | *Candidatus Wallbacteria bacterium* |
| | HauN | ABX04510.1 | *Herpetosiphon aurantiacus* DSM 785 |
| | DesN | HBV95591.1 | *Desulfotomaculum* sp. |
| | PahN | TNJ64878.1 | *Paenibacillus hemerocallicola* KCTC 33185 |
| | TaqN | WP_038049373.1 | *Thermoanaerobaculum aquaticum* MP-01 |
| | AbaN | RLE26118.1 | *Acidobacteria bacterium* isolate B25_G6 |
| **Lipoavitide** | StsE/SwaE | WP_030786714.1 | *Streptomyces sp. S-920/Streptomyces* sp. WAC 01529 |
| | LpvE/StaE | WP_136241767.1 | *Streptomyces* sp. NRRL S-1521/*Streptomyces* sp. A1499 |
| | SalE | WP_123985585.1 | *Streptomyces alfalfae* XN-04 |
| | SidE | WP_161255695.1 | *Streptomyces* sp. SID685 |
| | SedE | WP_161296757.1 | *Streptomyces* sp. SID161 |
| | PliE | WP_166354528.1 | *Phytoactinopolyspora limicola* HAJB-30 |
| | SmeE | WP_198503968.1 | *Streptomyces alfalfae* Men-myco-93-63 |
| | PcaE | WP_200323252.1 | *Prauserella cavernicola* ASG 168 |
| | StgE/SayE/SxyE/ SfrE/SacE | WP_205034037.1 | *Streptomyces* sp. G44/*Streptomyces alfalfae* XY25/*Streptomyces alfalfae* XY25 507/*Streptomyces fradiae* NKZ-259/*Streptomyces alfalfae* ACCC 40021 |

**Supplementary Table 9: Oligonucleotides used for cloning and site-directed mutagenesis**. The primers are named for the annealing region and whether the primer is for the forward (F) or reverse (R) direction.

| Name | Nucleotide Sequence (5' to 3' direction) |
|---|---|
| pBE44 F | ttaccaatgcttaatcagtgaggcacc |
| pBE45 R | atctttatagtcctgtcgggtttcg |
| pBE44-S-1521-70.1-R | accgcgacgccgttcgtcgacctcgggtacgaggacgtgaacttatatcgtatggggctg |
| pBE45-S-1521-70.1-F | gagcgggacgctttgaccgacgaggccgaggtgatgctggacgctcagtggaacgaaaac |
| S-1521-70.1-g1-F | aattaatacgactcactatagggaatttctactgttgtagatgtcgacctcgggtacgag |
| S-1521-70.1-g1-R | ctcgtacccgaggtcgacatctacaacagtagaaattccctatagtgagtcgtattaatt |
| S-1521-70.1-g2-F | aattaatacgactcactatagggaatttctactgttgtagataccgacgaggccgaggtg |
| S-1521-70.1-g2-R | cacctcggcctcgtcggtatctacaacagtagaaattccctatagtgagtcgtattaatt |
| LpvE KO F | gcgacaccgcctttccgcaaggctacgaacaggcccgcgagcagttgctgatgcattggcgggatcgttgtatatttcttgac |
| LpvE KO R | gcggagcaggcccaccatcaccgcgtccacccgcttgcccgcgggcgtcgatgcatttaccaatgcttaatcagtgaggcacc |
| LpvF KO F | cacgagatctggacgcaggggcacttcgaggccgtgccccgcttcatccaatgcattggcgggatcgttgtatatttcttgac |
| LpvF KO R | ctcctcgatgatcagcccgtcctccagacgggcgatgctcagcccccgatatgcatttaccaatgcttaatcagtgaggcacc |
| LpvG KO F | aacgacagcgtggcggaggccagcggagcggacccggcctggatacaggaatgcattggcgggatcgttgtatatttcttgac |
| LpvG KO R | tgctgtcatgccggcgccgacagcgaccagcgccacgaggtcccctcggcatgcatttaccaatgcttaatcagtgaggcacc |
| LpvH KO F | cccgccgactgcgcgcaggtggccgacatcgaggtacgcggcgaacgcgtatgcattggcgggatcgttgtatatttcttgac |
| LpvH KO R | tcgaggccgccgagaacgaaggggccgacgtcggtcccccggaagggtgtatgcatttaccaatgcttaatcagtgaggcacc |
| LpvI KO F | tccccctggtgccggtgctgtgggccctgcggagtgccgggcacgacgtaatgcattggcgggatcgttgtatatttcttgac |
| LpvI KO R | tgcgccgcgtcgttctcgtcccgcaggaccgccgccgccttcgcgtagccatgcatttaccaatgcttaatcagtgaggcacc |
| LpvJ KO F | actacccgctccagccccgcgatccgcggcagctcgcgcccttcgccagaatgcattggcgggatcgttgtatatttcttgac |
| LpvJ KO R | acctccagcaggtcccgcagggcttccaccgggccttccgccacgagcacatgcatttaccaatgcttaatcagtgaggcacc |
| LpvK KO F | ggttcgtcttccccgtggacgcgctcacacaggaccgggccgcggagctgatgcattggcgggatcgttgtatatttcttgac |
| LpvK KO R | gccatccgcatccccacggcgtcgccgcaccgggccatggcctcgtcgccatgcatttaccaatgcttaatcagtgaggcacc |
| LpvM1 KO F | ccgggatcggcgcgctcgacctgccgaacacggaactcgtcgagcccttcatgcattggcgggatcgttgtatatttcttgac |
| LpvM1 KO R | ctccgcgccggcttccgggtcgcggtgcgcggcgaccacggagaggccgtatgcatttaccaatgcttaatcagtgaggcacc |
| LpvS KO F | cgaggcccacctaccagcagctgcacgacacgggccccgtccaccgggtcatgcattggcgggatcgttgtatatttcttgac |
| LpvS KO R | ccgcgctgccattcgatggaatccgccgctatggcgagggtgaagtccggatgcatttaccaatgcttaatcagtgaggcacc |
| LpvV KO F | gactggccacggtggagaagttctcggcggccggtgcgggcgtcgtcatcatgcattggcgggatcgttgtatatttcttgac |
| LpvV KO R | gcgttctccgcgacgtgcagggccaggtccgcgtactcctccggccggccatgcatttaccaatgcttaatcagtgaggcacc |
| LpvA V1A F | tcgcggccctgacggagctggagacggacccgtcgctcatgccgaccgccgccggttcgatcatctacaccttcatgatccactgctgaatgcattggcgggatcgttgtatatt |
| LpvA V1K F | tcgcggccctgacggagctggagacggacccgtcgctcatgccgaccgccaagggttcgatcatctacaccttcatgatccactgctgaatgcattggcgggatcgttgtatatt |
| LpvA S3A F | tcgcggccctgacggagctggagacggacccgtcgctcatgccgaccgccgtcggtgccatcatctacaccttcatgatccactgctgaatgcattggcgggatcgttgtatatt |
| LpvA S3G F | tcgcggccctgacggagctggagacggacccgtcgctcatgccgaccgccgtcggtggcatcatctacaccttcatgatccactgctgaatgcattggcgggatcgttgtatatt |
| LpvA Y6F F | tcgcggccctgacggagctggagacggacccgtcgctcatgccgaccgccgtcggttcgatcatcttcaccttcatgatccactgctgaatgcattggcgggatcgttgtatatt |

| | |
|---|---|
| LpvA Y6W F | tcgcggccctgacggagctggagacggacccgtcgctcatgccgaccgccgtcggttcgatcatctg gaccttcatgatccactgctgaatgcattggcgggatcgttgtatatt |
| LpvA mutagenesis R | cggcggcatcactttctgccgcgtgccggccgcccgtcactgcgggaagaatgcatttaccaatgct taatcagtgagg |
| pET28a-NdeI-StsG F | gccgcgcggcagccatatgaccaccgcagcgcgtac |
| pET28a-HindIII-StsG R | tcgagtgcggccgcaagctttcagtaccagcgcagtacgg |
| pET28a-NdeI-LpvV F | gccgcgcggcagccatatgcacgtcgcggaaagcgt |
| pET28a-HindIII-LpvV R | tcgagtgcggccgcaagcttctaccccagccggaccgcgc |
| pET28a-NdeI-LpvE F | gccgcgcggcagccatatgtccggcaccgagtccag |
| pET28a-HindIII-LpvE R | tcgagtgcggccgcaagcttcatgactccgccaggagcc |
| pET28a-NdeI-StsE F | gccgcgcggcagccatatgaccggccagaagcgtcg |
| pET28a-HindIII-StsE R | tcgagtgcggccgcaagctttcaggcggtcgccagcttct |

**Supplementary Fig. 1: Predicted functions of enzymes in the *lpv* BGC.**

### Dha and Dhb

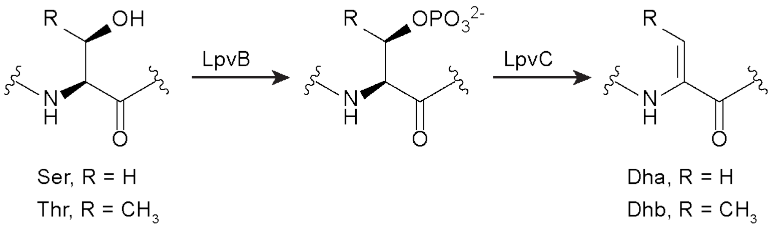

### Avi(Me)Cys

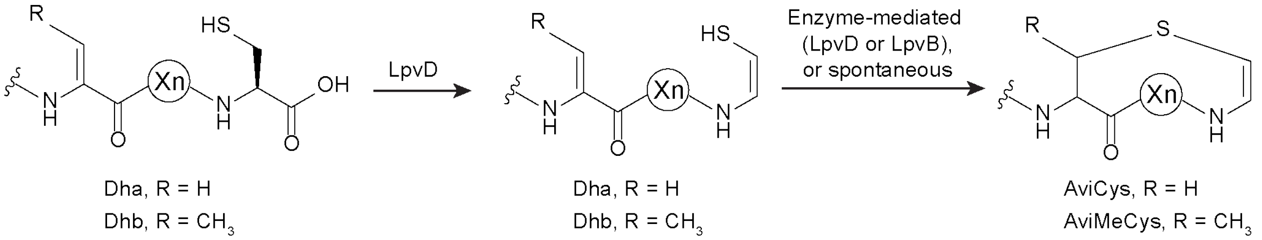

### *D*-alanine

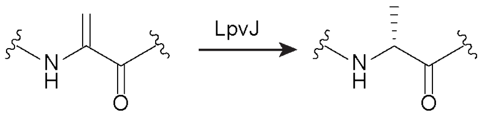

### hdmHis

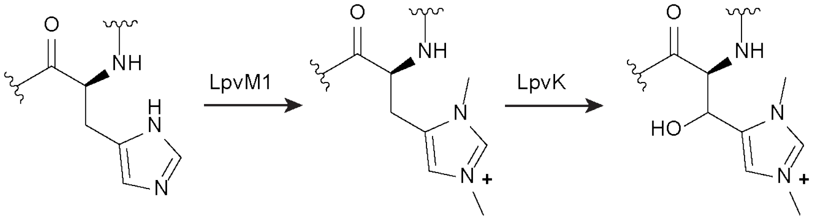

### glycosylated tyrosine

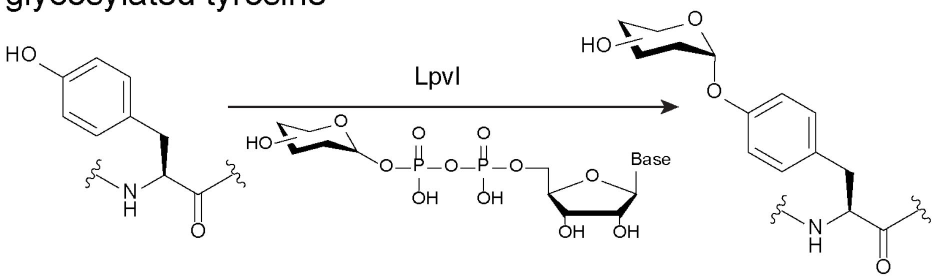

**Supplementary Fig. 2: Functional annotation of LpvP2.** (**a**) HHpred result for LpvP2[17] (RefSeq ID: WP_062773482.1, Uniprot ID: A0A0X3WL74). Analysis was performed using the PDB_mmCIF70_10_Jan database . (**b**) Foldseek[18] results for LpvP2 using AlphaFold/Swissprot database (version 4). Q: LpvP2; T: Metallopeptidase ImmA from *Bacillus subtilis* subsp. *subtilis* 168 (Uniprot ID: P96630). Sequence identity: 16.3%. E-value: 1E-3. In the overlaid structures, LpvP2 and ImmA are shown in cyan and yellow, respectively. Template modeling (Tm)-score: 0.69446. Root mean square deviation (RMSD): 7.31. (**c**) Zoomed view of the aligned HExxH motif. Nitrogen and oxygen atoms are shown in blue and red, respectively.

**a**

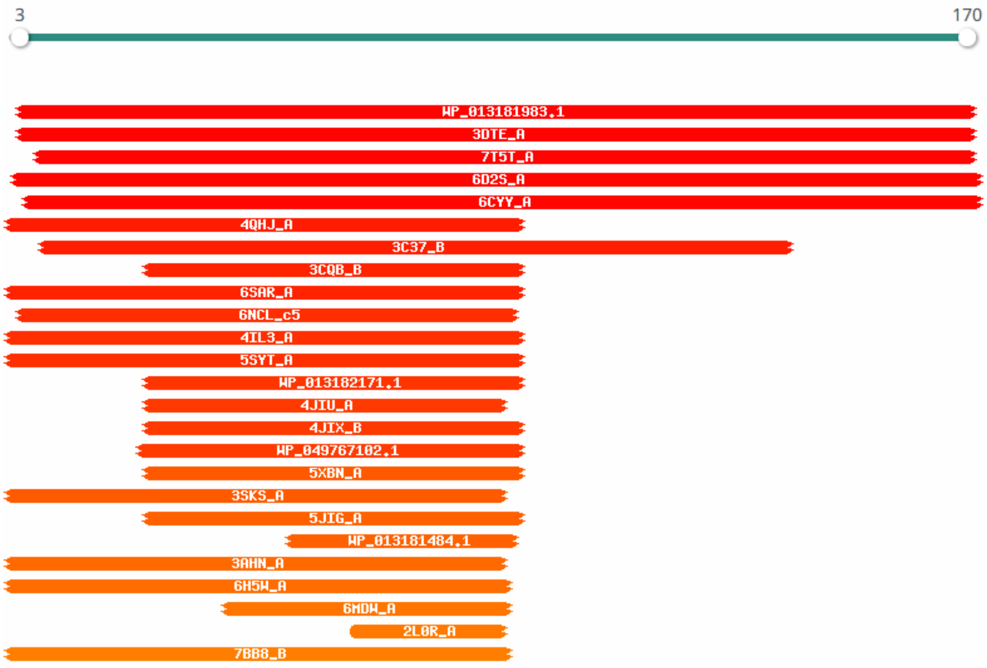

Hitlist

Show 25 Entries Search:

| Nr | Hit | Name | Probability | E-value | Score | SS | Aligned cols | Target Length |
|---|---|---|---|---|---|---|---|---|
| 1 | WP_013181983.1 | ImmA/IrrE family metallo-endopeptidase [Waddlia chondrophila] | 99.74 | 3.9e-17 | 111.05 | 10.9 | 145 | 169 |
| 2 | 3DTE_A | IrrE protein; Deinococcus, Radiotolerance, Gene regulation, Metallopeptidase, IrrE; HET: MSE; 2.6A {Deinococcus deserti} | 99.66 | 1e-15 | 113.56 | 10 | 144 | 301 |
| 3 | 7T5T_A | CapP toxin; Zinc metallopeptidase, IrrE, cysteine switch, HYDROLASE; HET: NHE, SO4; 1.35A {Thauera sp. K11} | 99.62 | 2.4e-15 | 111.12 | 8.3 | 149 | 291 |
| 4 | 6D2S_A | HTH-type transcriptional regulator PrpR; Transcriptional regulator, TRANSCRIPTION; HET: EDO; 1.819A {Mycobacterium tuber | 99.52 | 1.7e-13 | 101.15 | 10.5 | 143 | 289 |

**b**

```
Q   1 MKERELRRHCKRTLRSLGIQPPLRVRELCRLLGEHRGRPIRLVPYSLPVPGPSGLWIATG--KTDYIVFQSETSKAHQDH
         + ++   +  +++ G      V E+C +        I ++     +   +GL           I  +      +
T   5 YTSKGIKHKVQSVIKTHG-TNN--VYEICDIQK------IYILKND--LGQANGLLQHDKATDQYLIHINENLQHQQ--F

Q  79 IILHEIGHMMAGHQSPAAGAELWRTMLPDISPSVITSMLGRTSYDEDREREAELVAT-IIL
       +I HE+GH +  H+            + + S  V+   L         E +A L A+ +IL
T  72 VIAHELGHYFL-HKRLNT------FKVVNCS-KVLKDKL---------EHQASLFASELIL
```

Tm-score: 0.69446
RMSD: 7.31

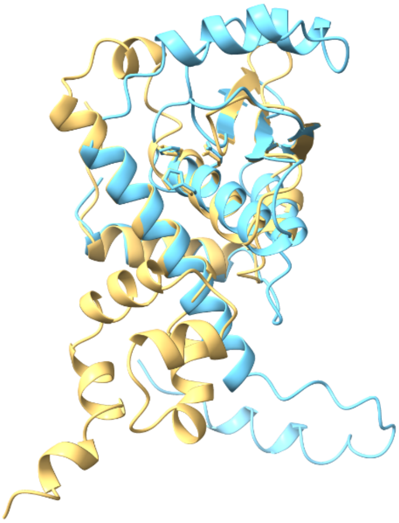

**c**

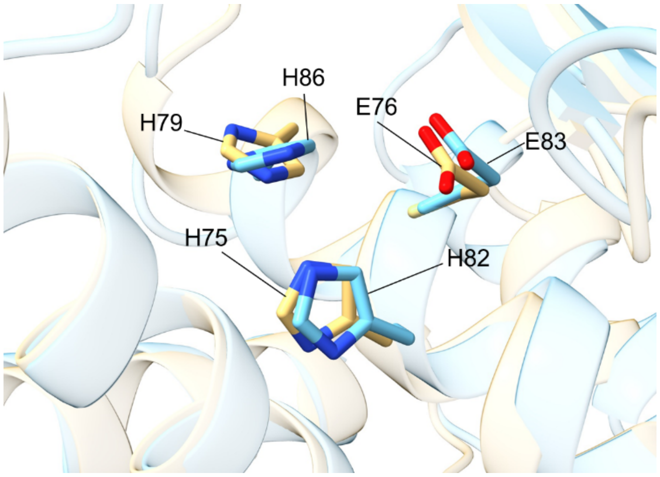

**Supplementary Fig. 3: Enzyme functions in fatty acid biosynthesis**. (**a**) Initiation of type II fatty acid synthesis. FabD catalyzes the transfer of the malonyl group from the malonyl-CoA to an ACP. The malonyl-ACP is then condensed with acetyl-CoA, catalyzed by FabH, to form the substrate for the first elongation cycle, acetoacetyl-ACP. (**b**) Elongation of type II fatty acid synthesis. The β-ketoacyl-ACP is reduced to β-hydroxyacyl-ACP by FabG, which is subsequently dehydrated to (*E*)-2-enoyl-ACP by FabZ (or FabA). The *trans* C2–C3 carbon–carbon double bond is then reduced by FabI, followed by the next round of condensation with malonyl-CoA unit catalyzed by FabB or FabF.

**a**

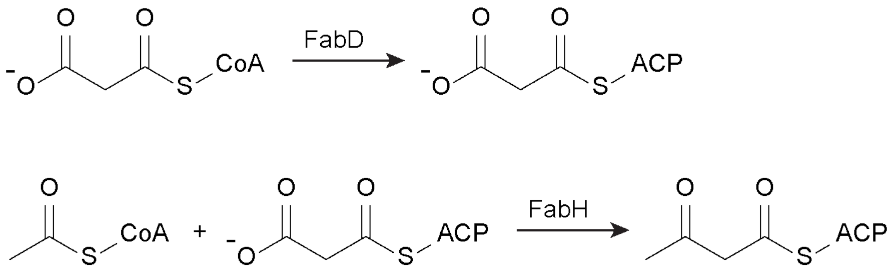

**b**

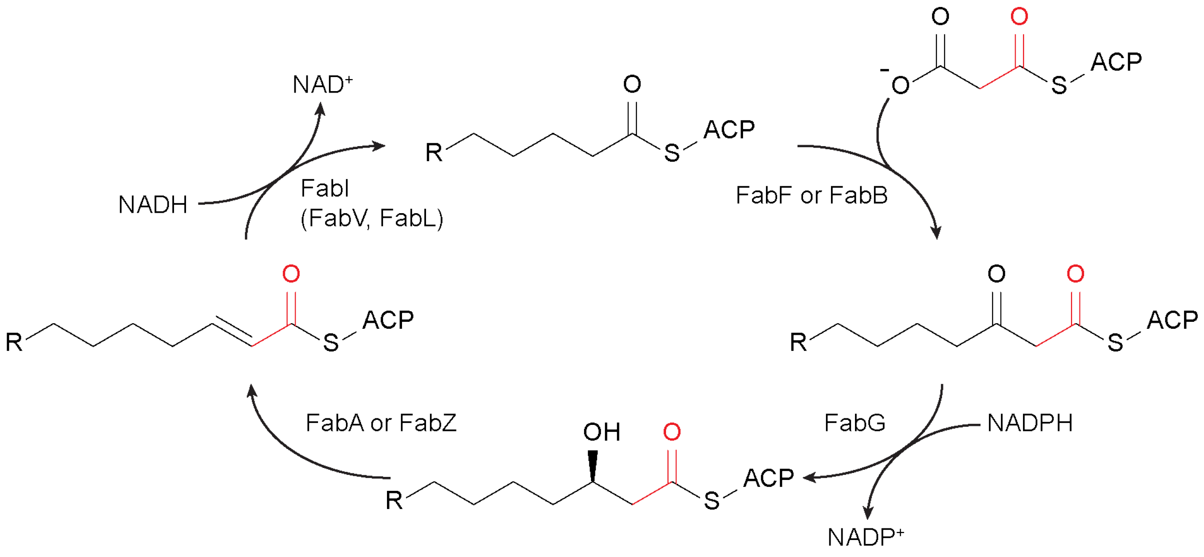

**Supplementary Fig. 4: InterProScan results of LpvV and LpvH**. (**a**) Domains identified in LpvV (RefSeq ID: WP_062773474.1) using InterProScan[21]. LpvV is predicted to function as the FabG in **Supplementary Fig. 3**. Short-chain dehydrogenase/reductase SDR (IPR002347, PR00080, PR00081, PF00106, PTHR43658); NAD(P)-binding domain superfamily (IPR036291, SSF51735, G3DSA:3.40.50.720); Short-chain dehydrogenase/reductase, conserved site (IPR020904, PS00061); 3-hydroxyacyl-CoA dehydrogenase type-2 (G3DSA:3.40.50.720:FF:000215), PKS_KR (SM00822). (**b**) Domains identified in LpvH (RefSeq ID: WP_159043818.1) using InterProScan. LpvH is predicted to function as the FabI in **Supplementary Fig. 3**. Alcohol dehydrogenase-like, C-terminal (IPR013149, PF00107); Polyketide synthase, enoylreductase domain (IPR020843, SM00829); NAD(P)-binding domain superfamily (IPR036291, SSF51735, G3DSA:3.40.50.720); GroES-like superfamily (IPR011032, SSF50129); Medium chain reductase/dehydrogenase (MDR)/zinc-dependent alcohol dehydrogenase-like family (cd05188, G3DSA:3.90.180.10).

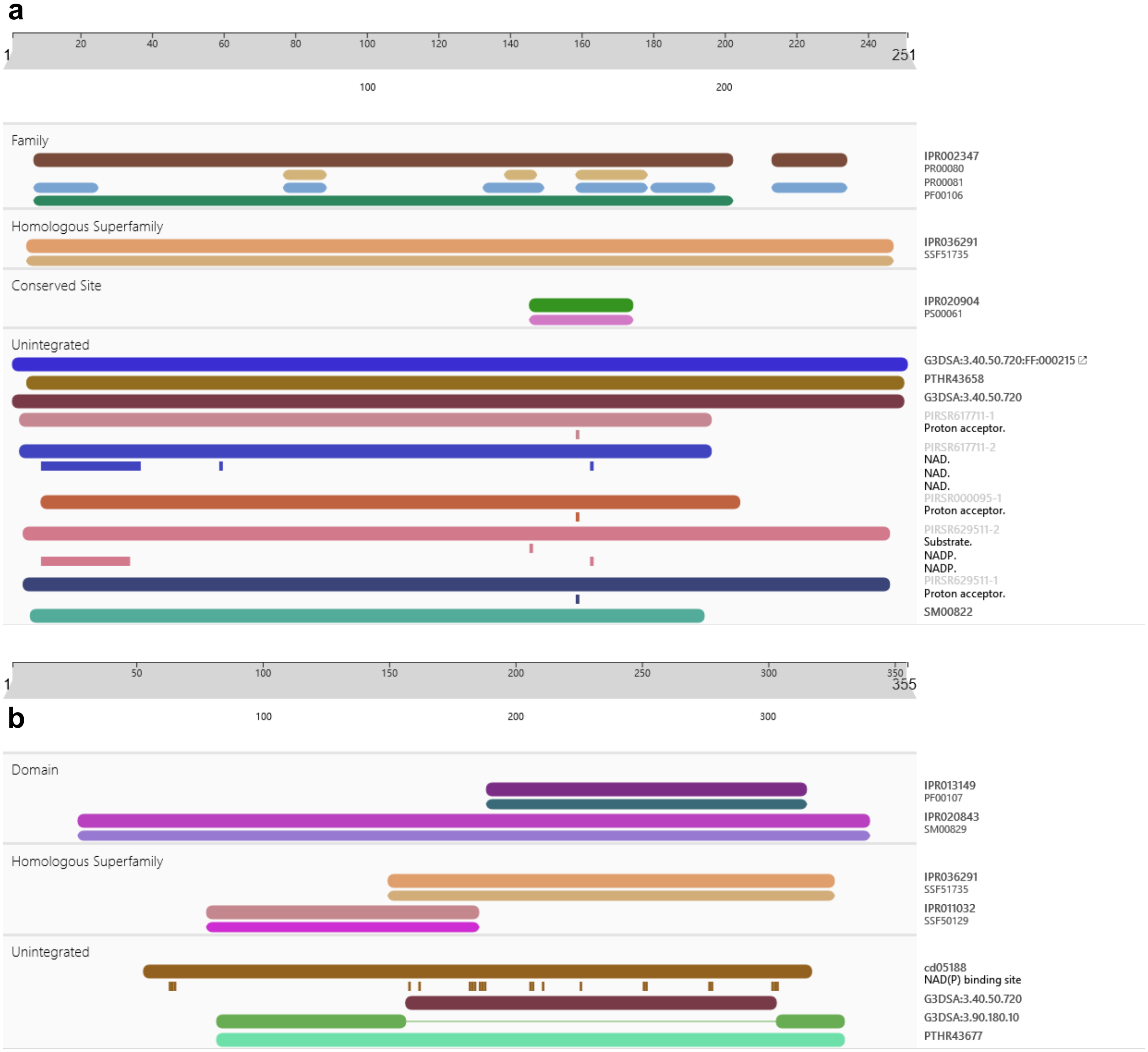

**Supplementary Fig. 5: Homologous *lpv* BGCs identified in bioinformatic analysis**. (**a**) Gene organization of homologous *lpv* BGCs. (**b**) Precursor peptide sequences of homologous *lpv* BGCs. The host species, NCBI accession identifer of nucleotide sequences, and the range in which the BGC was identified, are provided in the heading for each BGC. Multiple headings shown above one BGC indicate that BGCs with the same gene arrangement and precursor peptide sequence are found in more than one host strain. Genes are named according to their functional counterparts in the *lpv* BGC. Annotations of genes that have no counterparts in the *lpv* BGC: *R*, regulator; *Deh*, dTDP-4-dehydro-6-deoxy-α-*D*-glucopyranose 2,3-dehydratase (homologous to OleV, Uniprot ID: Q9RR31, coverage/identity 90%/50%); *M*: S-adenosylmethionine-dependent methyltransferase (homologous to NovU, Uniprot ID: Q9L9E7, coverage/identity 97%/34%); *Ami*: dTDP-3-amino-3,4,6-trideoxy-α-*D*-glucose transaminase (homologous to MegDII, Uniprot ID Q9F837, coverage/identity 99%/66%); *Epi*: UDP-glucose 4-epimerase (MJ_RS01105, Uniprot ID: Q57664, coverage/identity 94%/21%).

**a**

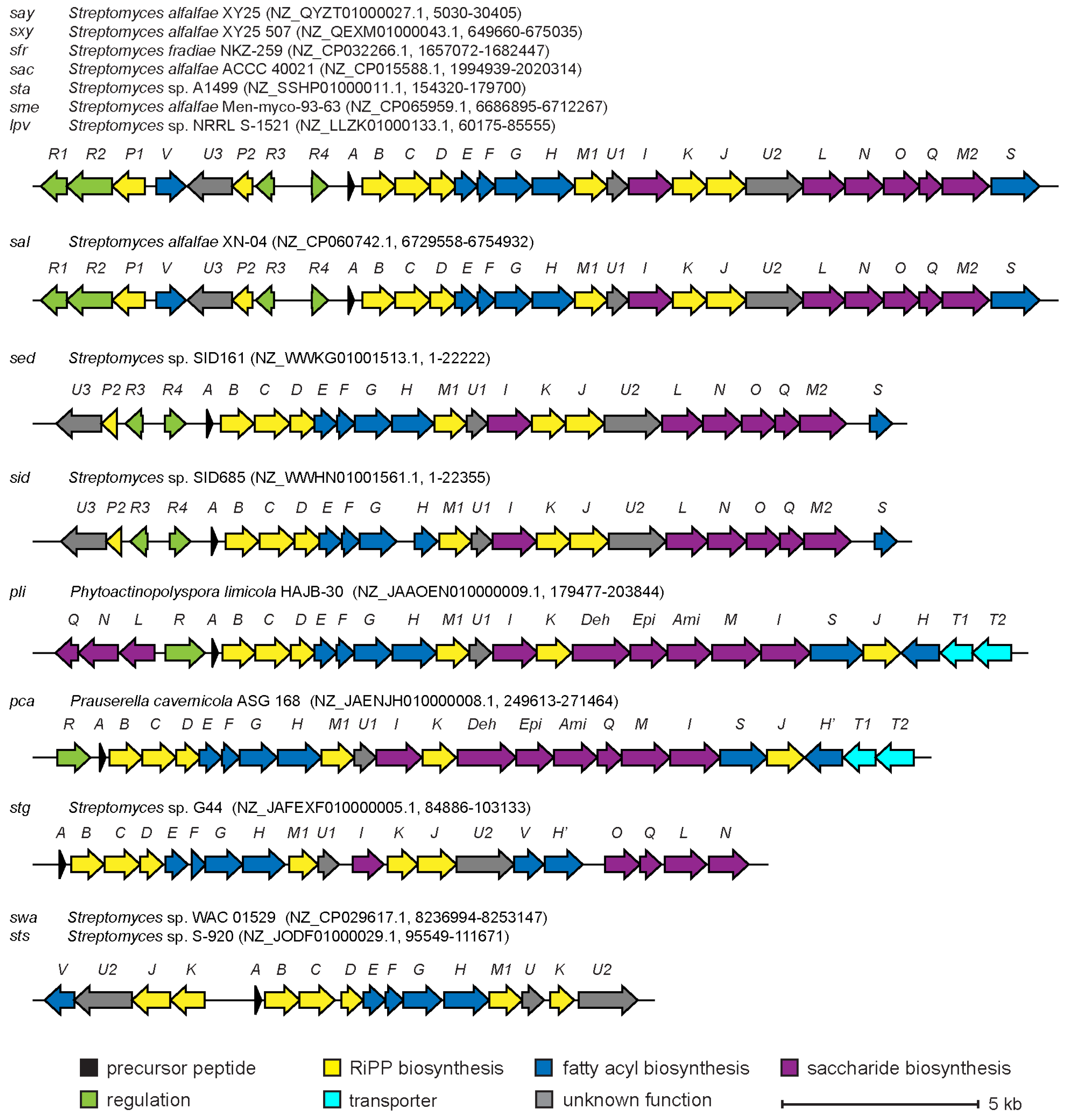

**b** SayA, SxyA, SfrA, SacA, StaA, SmeA, LpvA

MSATEIVVEDVLAELLDGDDVAALTELETDPSLMPTAVGSIIYTFMIHC

SalA

MSATEIVVEDVLAELLDGDDVAALTELETDPSLMPTSVGSIIYTFMIHC

SedA

MSATEIVVEDVLAELLDGDDVAALTELETDPSLMPTAVGSVIYTVMIHC

SidA

MSATEIVVEDVLAELLDGDDVAALTELETDPSLMPTAVGSVIYTVMIHC

PliA

MVTKEMELEDVLGTLLEGETTESIADLEIDPIQMPVAKSSITYTIVIHC

PcaA

MATKEYEIEDVLGTLLEAEAAESIADLEIDPVQMPVAKSSVTYTVAVHC

StgA

MPNAELTVEDLLADLLEADEVDGLADLEIDPTEMPTAKGSVIYTTMLHC

SwaA
StsA

MENTTELAVEDILSTLLDDETAGDLAELEIDPSQMPVAKSSVVWTVTAHC

**Supplementary Fig. 6: Media and host screening for *lpv* BGC expression**. MALDI-TOF mass spectra of methanolic extracts of *S. lividans* TK24 and *S. albus* J1074 colonies that were transformed with *lpv* or the pBE45 empty vector and cultivated on MS and ISP4 agar media.

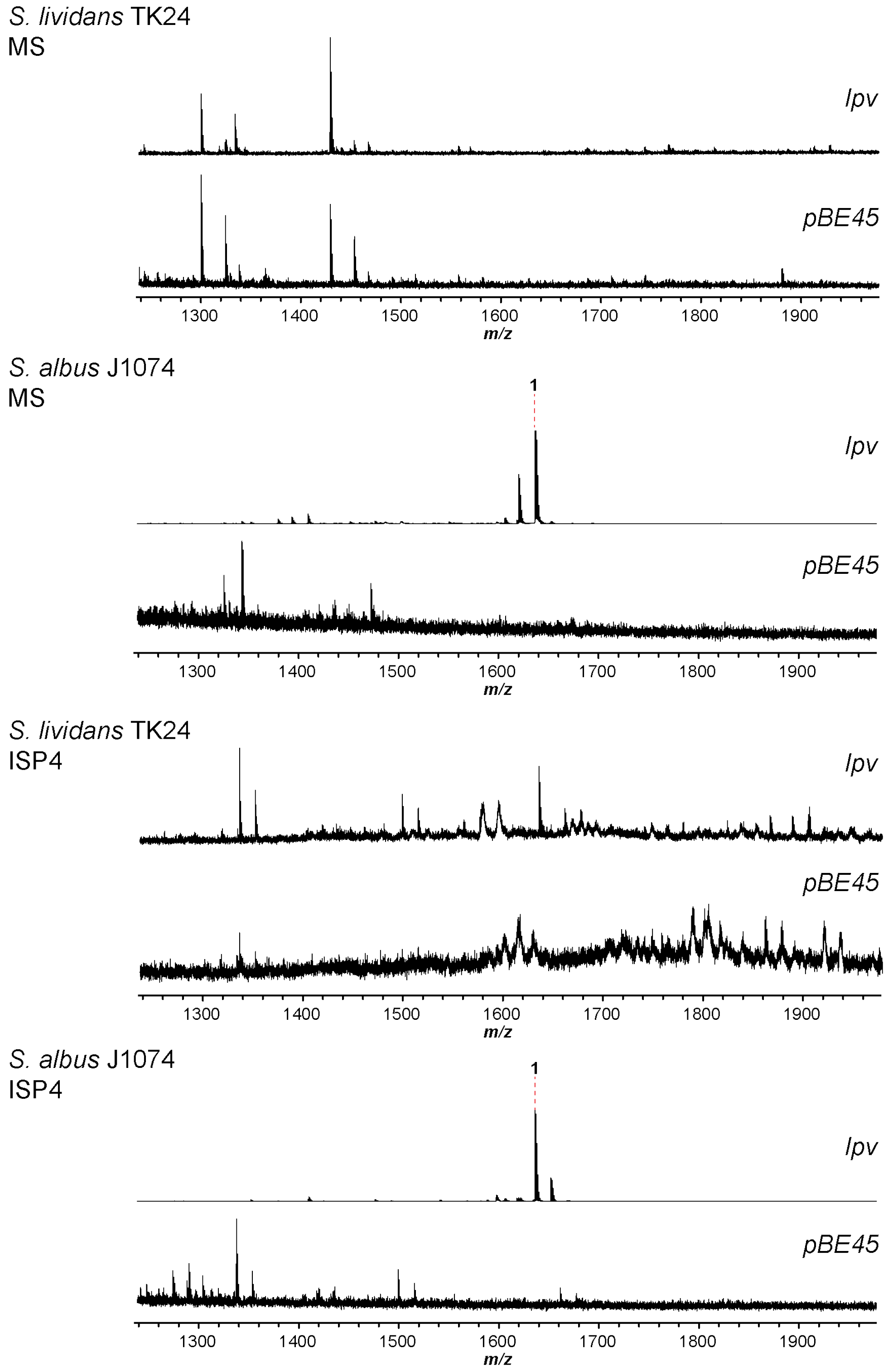

**Supplementary Fig. 7: Purification of lipoavitide 1 by preparative HPLC**. (**a**) Reverse phase (C5) HPLC chromatograms with UV absorbance (220 nm) monitoring of solid phase extraction eluents using 60% acetonitrile. (**b**) MALDI-TOF-MS analysis of the collected fractions.

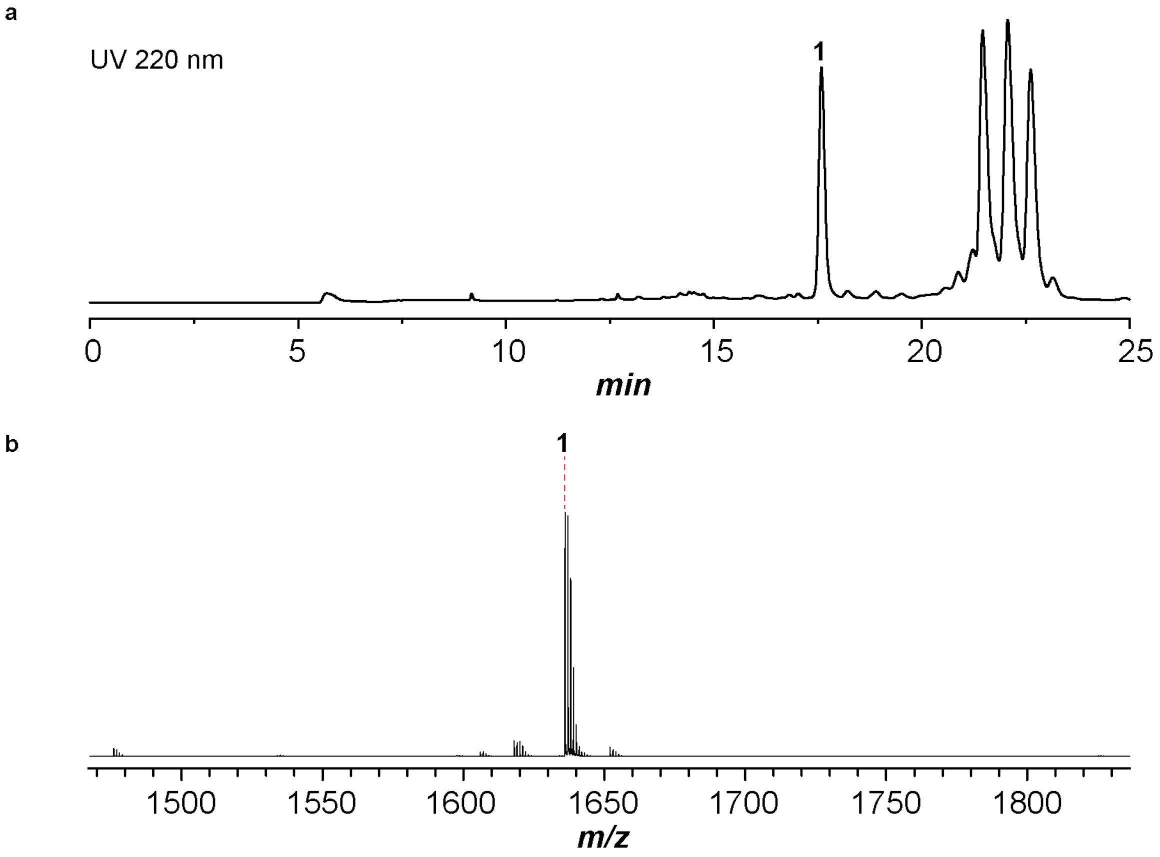

**Supplementary Fig. 8: HR-MS/MS analysis of 1**. (**a**) Broad band high-resolution mass spectrum of compound **1**. (**b**) Summary of collision-induced dissociation (CID) daughter ions for **1**. All ions are in the +1 charge state. "Retro aldol" indicates the formation of 4-formyl-1,3-dimethyl-1*H*-imidazolium through retro aldol reaction during MS/MS analysis. (**c**) CID spectrum of **1**. (**d**) Enlarged CID spectrum of **1** in the dashed box region. (**e**) Table of daughter ion assignments for **1**.

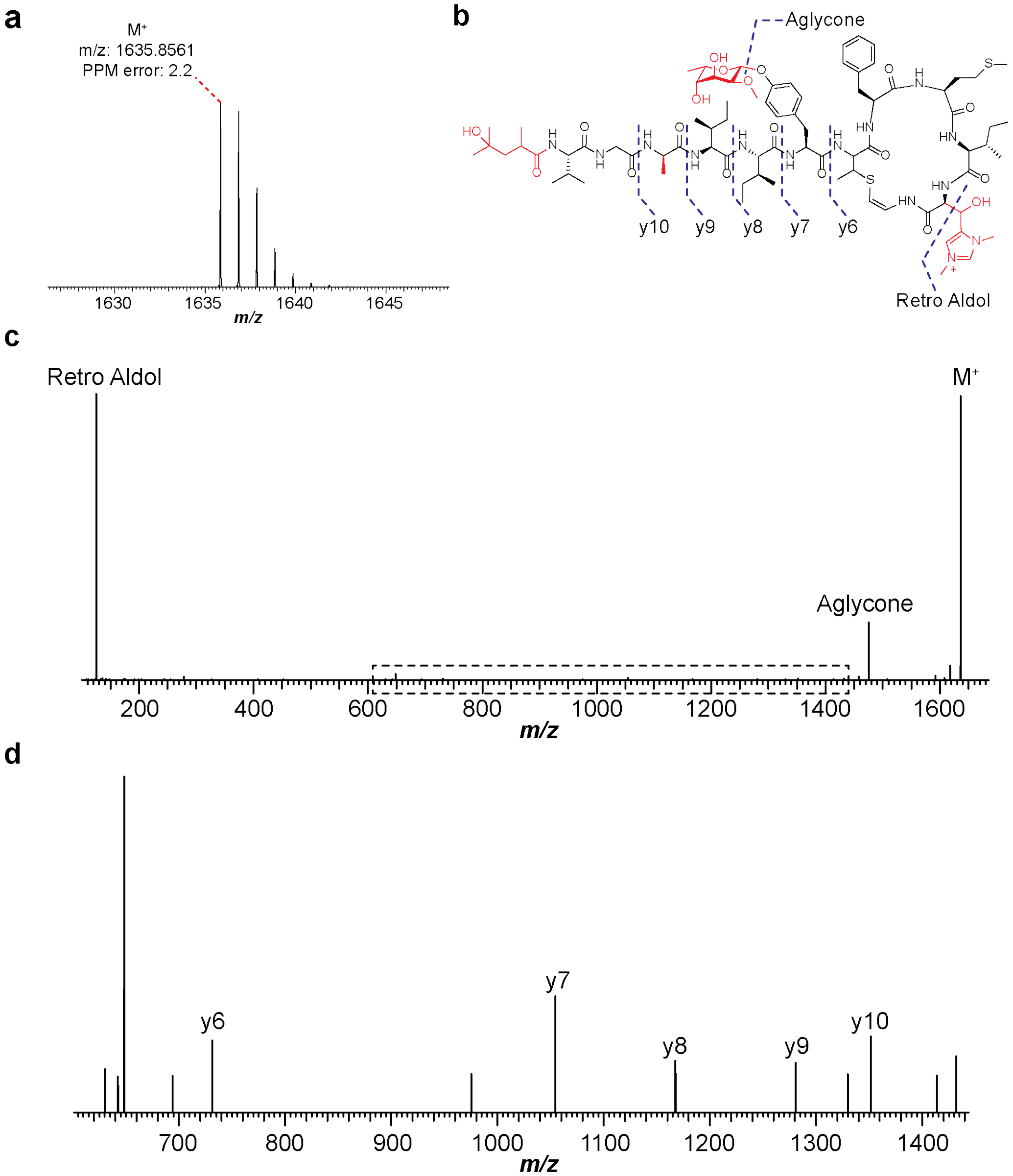

e

| Ion | Calc. Mass | Obs. Mass | Ppm Error |
|---|---|---|---|
| **M⁺** | 1635.8525 | 1635.8583 | 3.5 |
| **Aglycone** | 1475.7789 | 1475.7838 | 3.3 |
| **Retro Aldol** | 125.0709 | 125.0716 | 5.6 |
| **y6** | 731.3368 | 731.3397 | 4.0 |
| **y7** | 1054.4736 | 1054.4780 | 4.2 |
| **y8** | 1167.5577 | 1167.5594 | 1.5 |
| **y9** | 1280.6418 | 1280.6458 | 3.1 |
| **y10** | 1351.6789 | 1351.6790 | 0.1 |

**Supplementary Fig. 9: $^{1}$H NMR of lipoavitide 1 in acetone-$d_6$**. The spectrum was obtained in acetone-$d_6$. (**a**) Full spectrum; (**b-d**) Enlarged views of the spectrum.

**a**

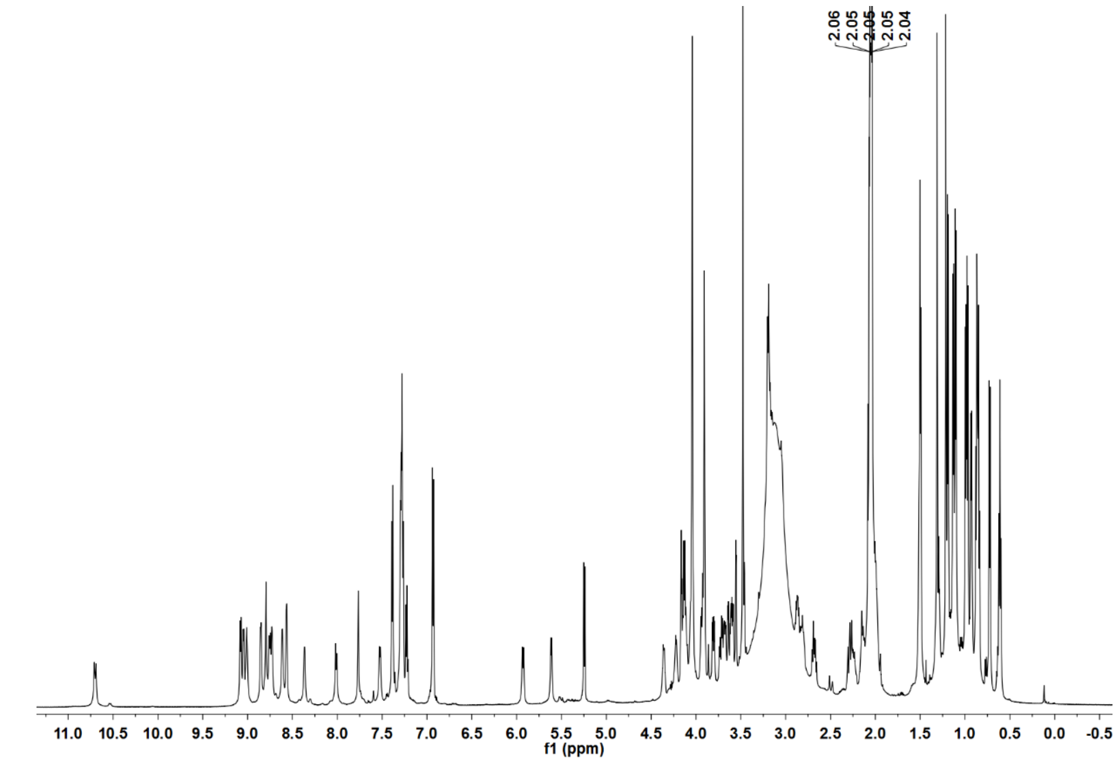

**b**

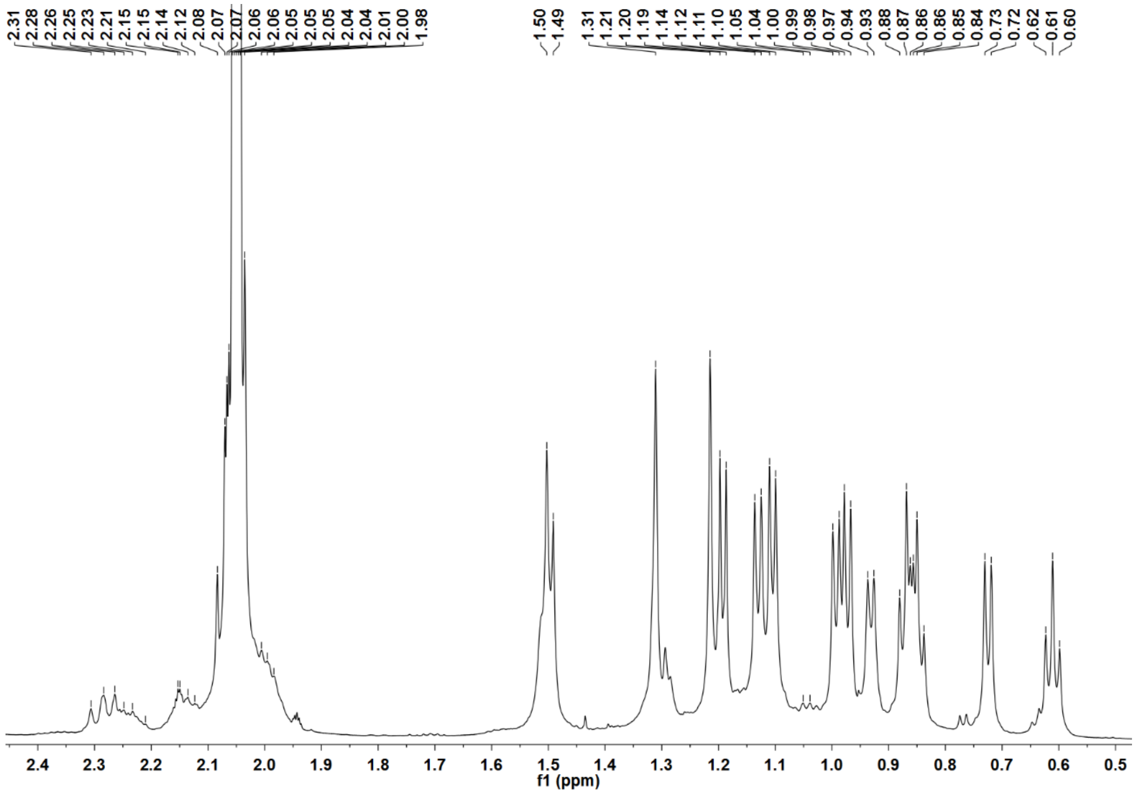

c

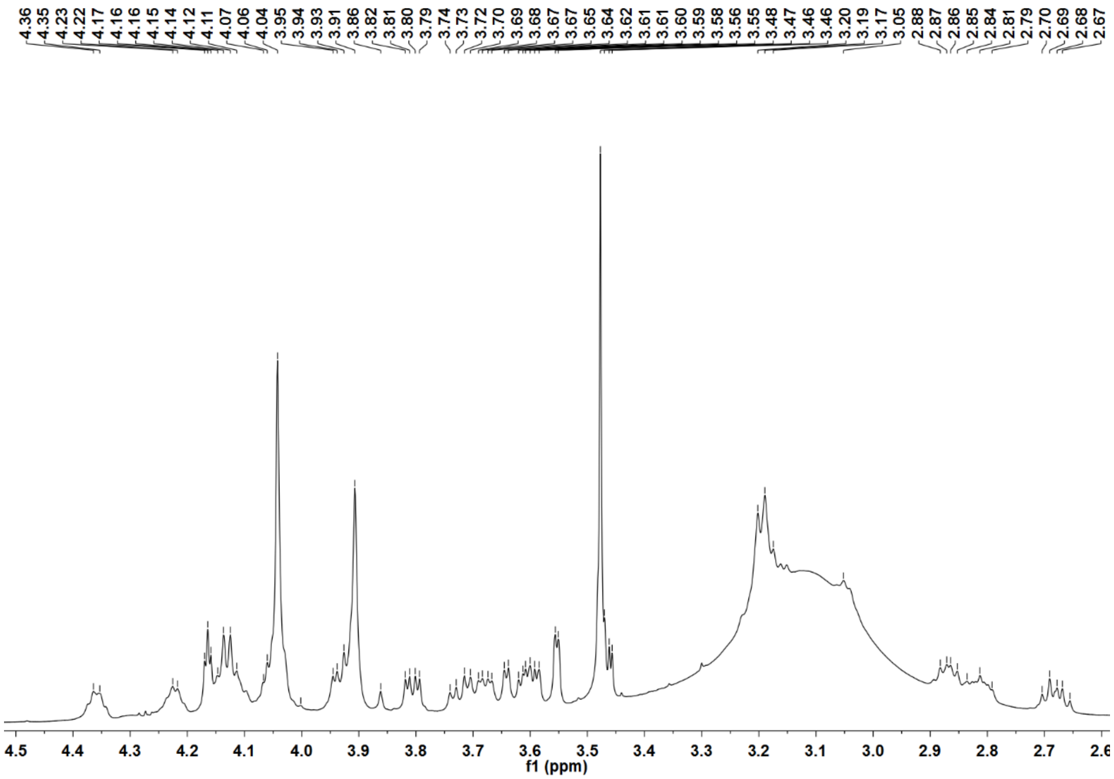
4.36
4.35
4.23
4.22
4.17
4.16
4.16
4.15
4.14
4.12
4.11
4.07
4.06
4.04
3.95
3.94
3.93
3.91
3.86
3.82
3.81
3.80
3.79
3.74
3.73
3.72
3.70
3.69
3.68
3.67
3.67
3.65
3.64
3.62
3.61
3.61
3.60
3.59
3.58
3.56
3.55
3.48
3.47
3.46
3.46
3.20
3.19
3.17
3.05
2.88
2.87
2.86
2.85
2.84
2.81
2.79
2.70
2.69
2.68
2.67
f1 (ppm)

d

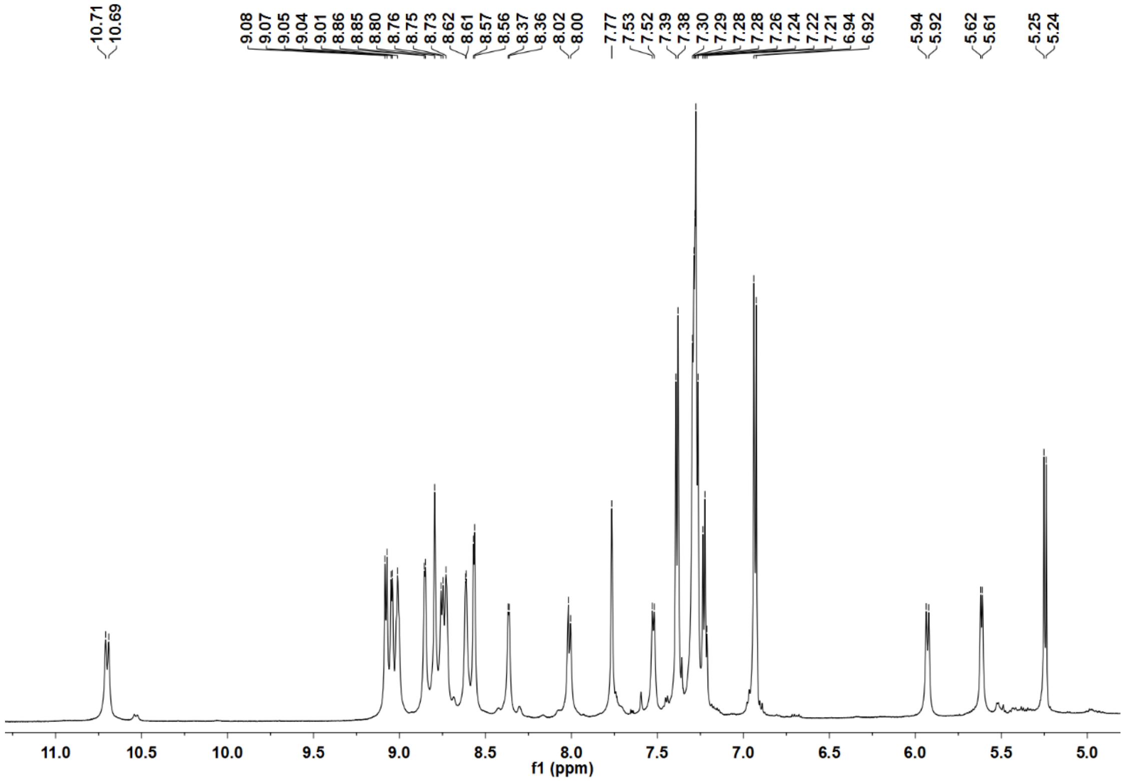
10.71
10.69
9.08
9.07
9.05
9.04
9.01
8.86
8.85
8.80
8.76
8.75
8.73
8.62
8.61
8.57
8.56
8.37
8.36
8.02
8.00
7.77
7.53
7.52
7.39
7.38
7.30
7.29
7.28
7.26
7.24
7.22
7.21
6.94
6.92
5.94
5.92
5.62
5.61
5.25
5.24
f1 (ppm)

**Supplementary Fig. 10: $^{13}$C NMR of lipoavitide 1 in acetone-$d_6$.**

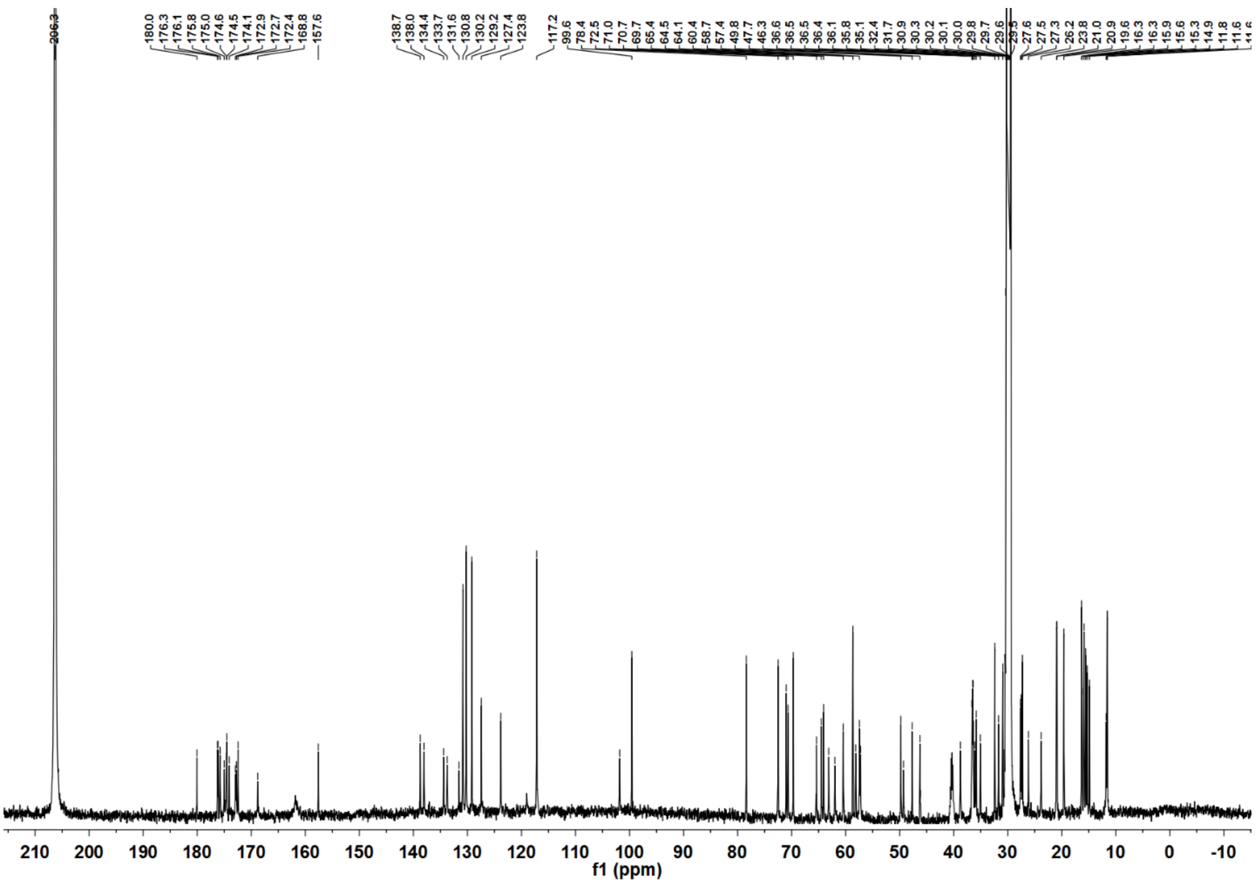

**Supplementary Fig. 11: $^1$H-$^1$H COSY of lipoavitide 1 in acetone-*d*$_6$.** (**a**) Full spectrum; (**b**, **c**) Enlarged views of the spectrum.

**a**

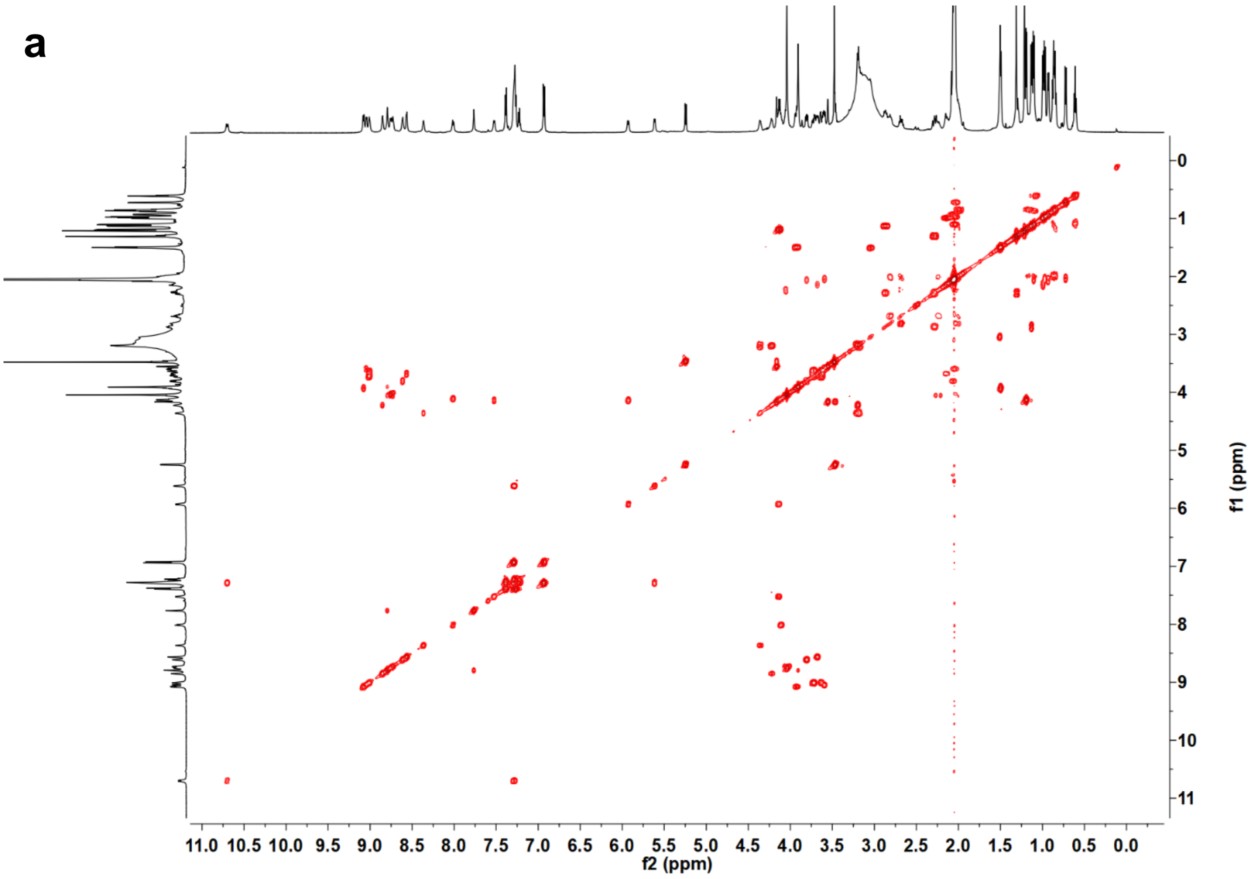

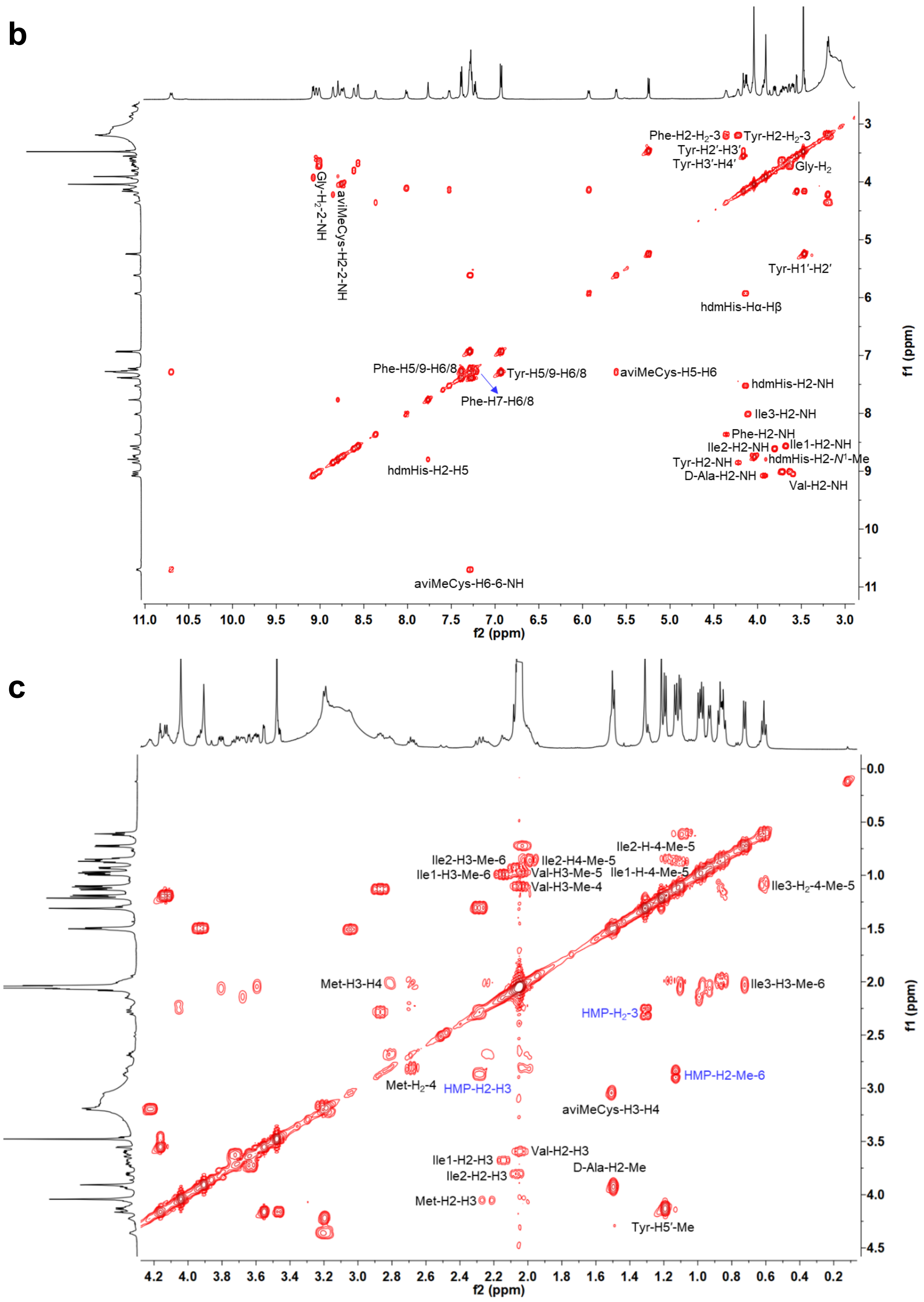

b
Phe-H2-H$_2$-3
Tyr-H2-H$_2$-3
Tyr-H2′-H3′
Tyr-H3′-H4′
Gly-H$_2$
Gly-H$_2$-2-NH
aviMeCys-H2-2-NH
Tyr-H1′-H2′
hdmHis-Hα-Hβ
Phe-H5/9-H6/8
Tyr-H5/9-H6/8
aviMeCys-H5-H6
Phe-H7-H6/8
hdmHis-H2-NH
Ile3-H2-NH
Phe-H2-NH
Ile2-H2-NH
Ile1-H2-NH
hdmHis-H2-$N^1$-Me
Tyr-H2-NH
D-Ala-H2-NH
Val-H2-NH
hdmHis-H2-H5
aviMeCys-H6-6-NH
f2 (ppm)
f1 (ppm)
c
Ile2-H-4-Me-5
Ile2-H3-Me-6
Ile2-H4-Me-5
Val-H3-Me-5
Ile1-H3-Me-6
Ile1-H-4-Me-5
Val-H3-Me-4
Ile3-H$_2$-4-Me-5
Met-H3-H4
Ile3-H3-Me-6
HMP-H$_2$-3
HMP-H2-Me-6
Met-H$_2$-4
HMP-H2-H3
aviMeCys-H3-H4
Val-H2-H3
Ile1-H2-H3
D-Ala-H2-Me
Ile2-H2-H3
Met-H2-H3
Tyr-H5′-Me
f2 (ppm)
f1 (ppm)

**Supplementary Fig. 12: $^{1}H$-$^{13}C$ HSQC of lipoavitide 1 in acetone-$d_6$.** Multiplicity-edited HSQC was performed. Signals of CH2 and CH/CH3 are shown in blue and red, respectively.

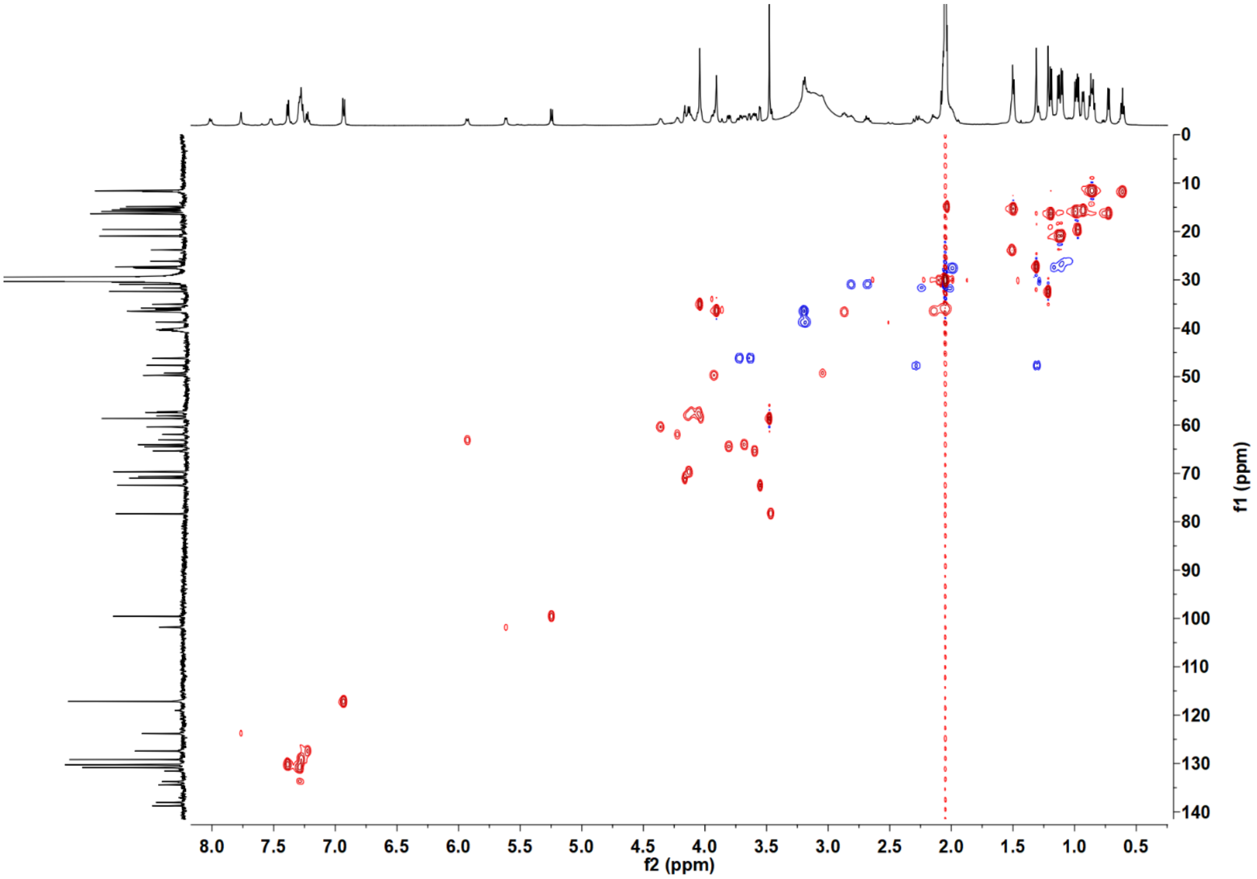

**Supplementary Fig. 13: $^{1}$H-$^{13}$C HMBC of lipoavitide 1 in acetone-$d_6$.**

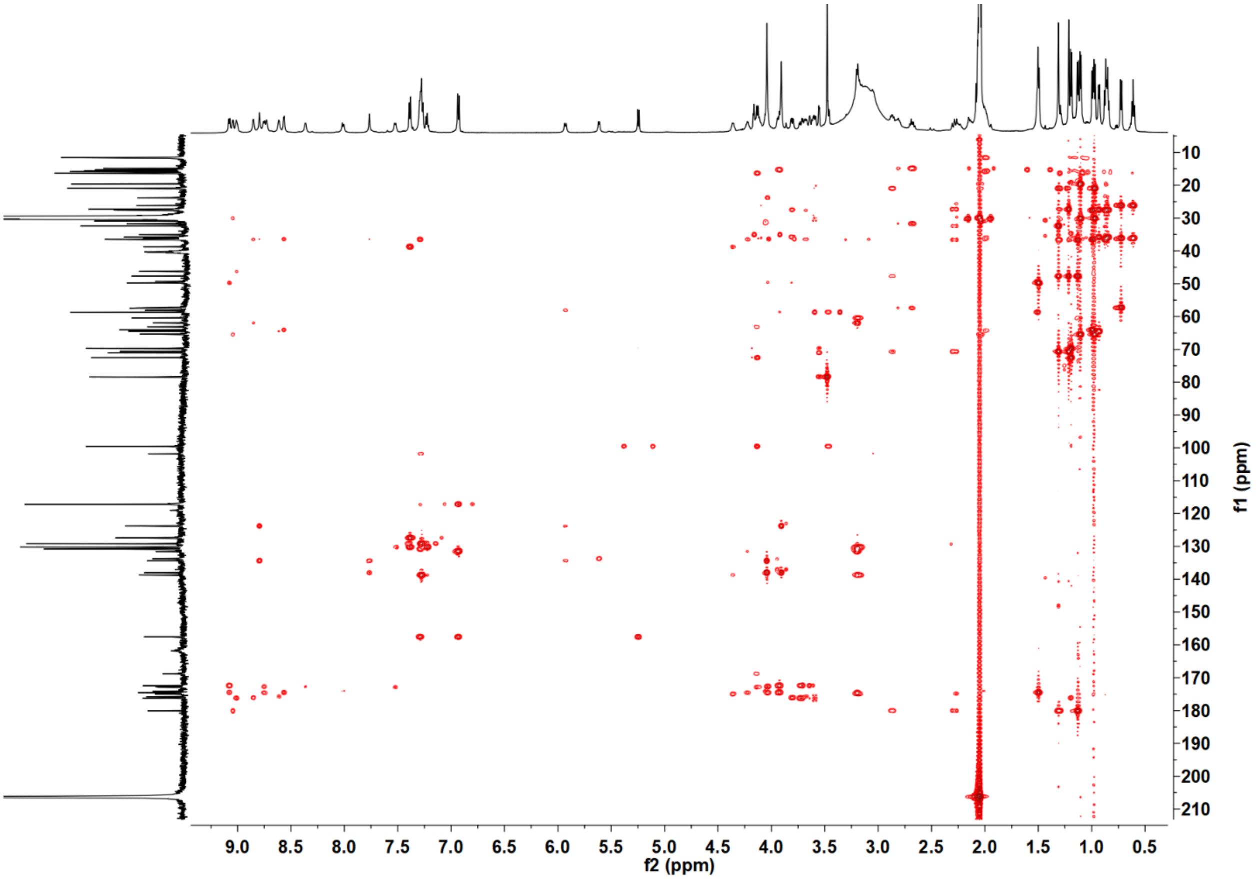

**Supplementary Fig. 14: $^{1}H$-$^{13}C$ HMBC of lipoavitide 1 in acetone-$d_6$ focusing on the HMP region.**

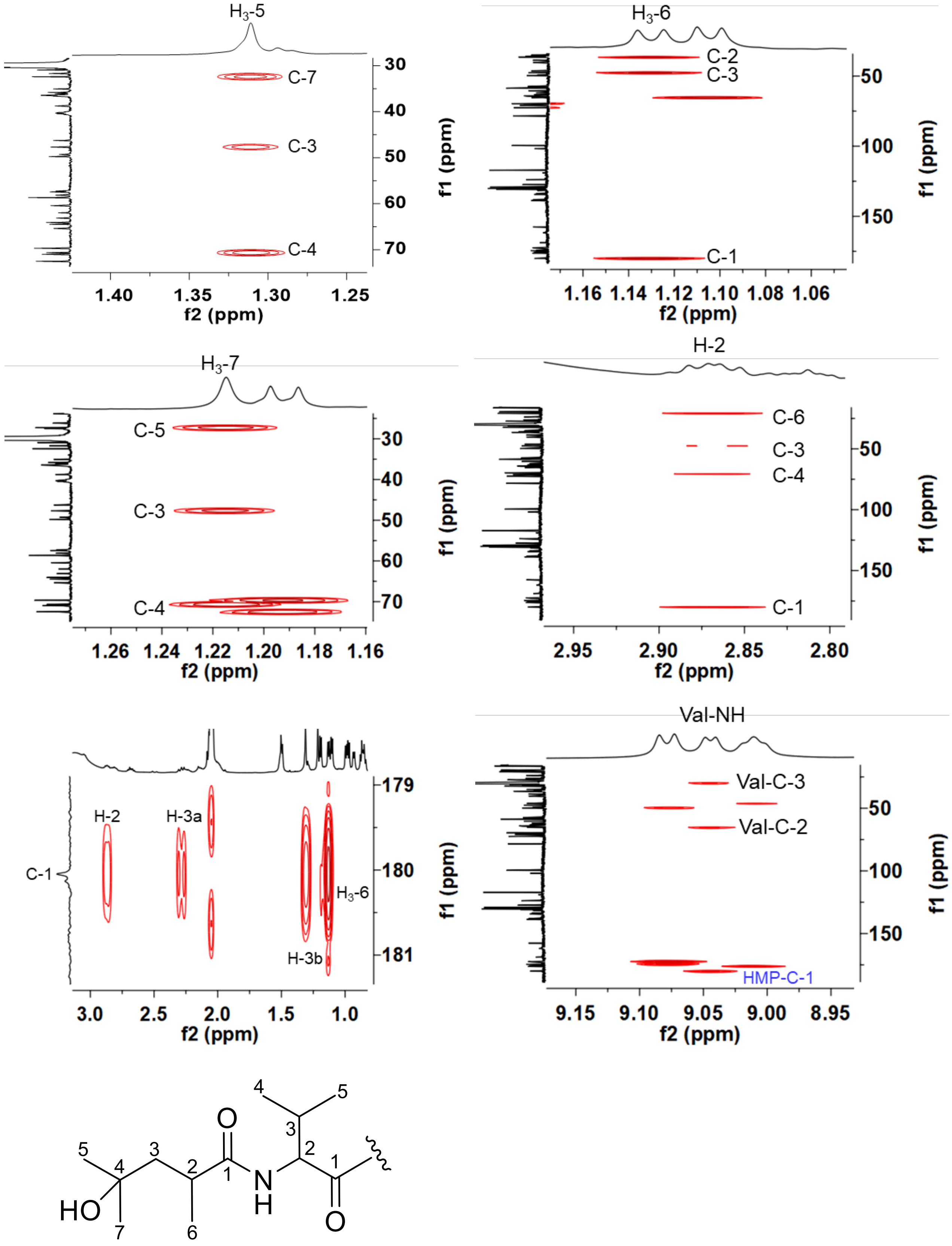

**Supplementary Fig. 15: $^{1}H$-$^{13}C$ HMBC of lipoavitide 1 in acetone-$d_6$ focusing on the *D*-alanine region**.

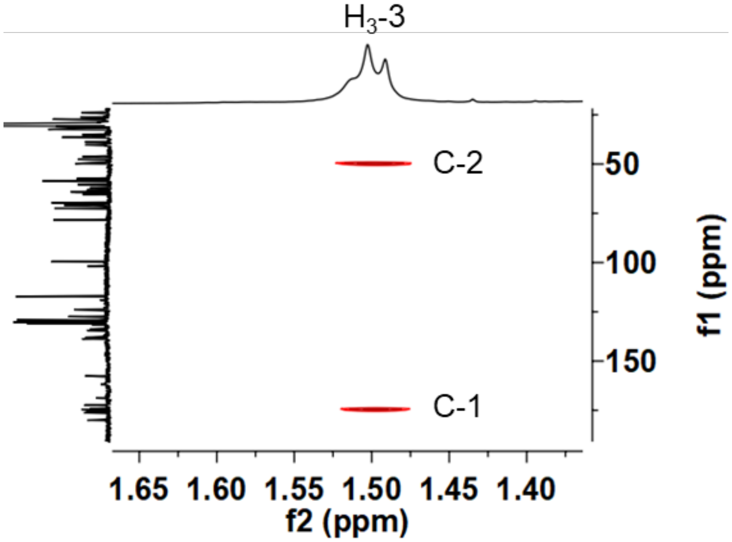

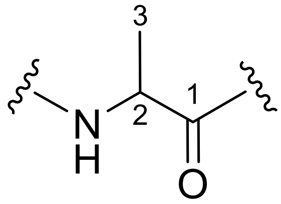

**Supplementary Fig. 16: $^{1}H$-$^{13}C$ HMBC of lipoavitide 1 in acetone-$d_6$ focusing on the 2-*O*-methyl-β-6-deoxygulose region**.

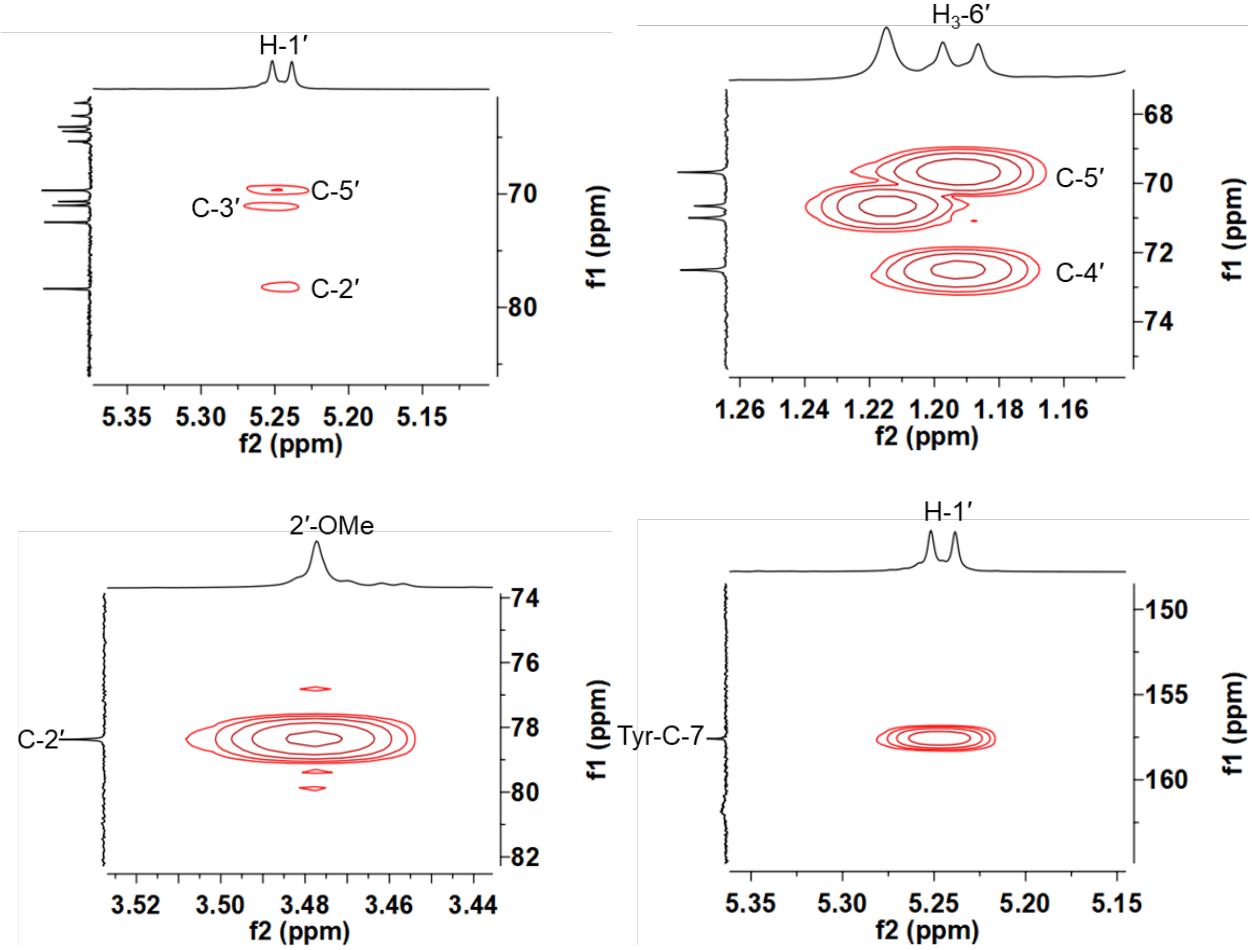

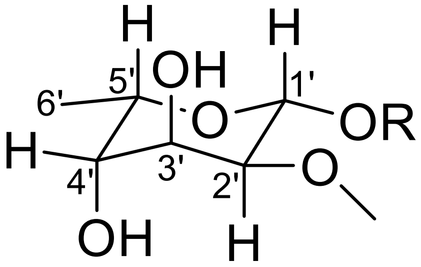

**Supplementary Fig. 17: [1]H-[13]C HMBC of lipoavitide 1 in acetone-$d_6$ focusing on the AviMeCys region**.

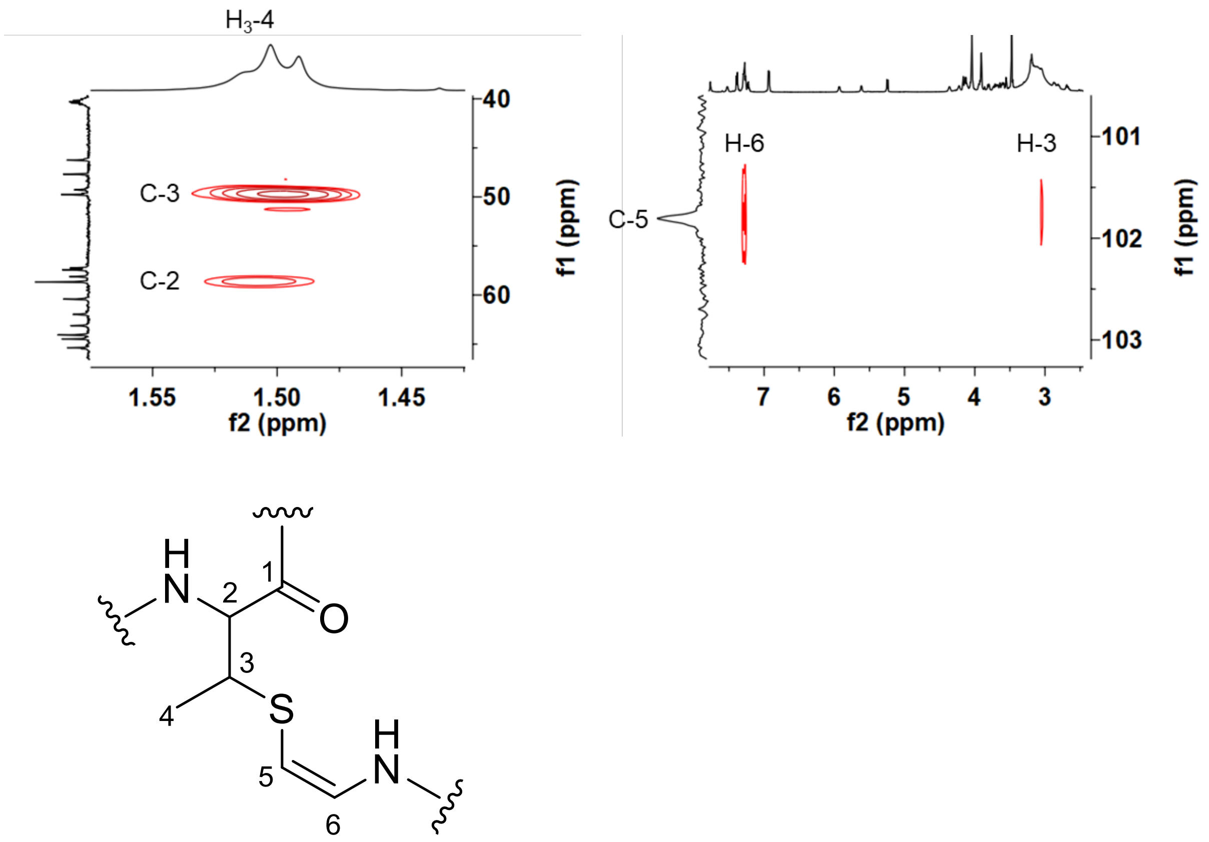

**Supplementary Fig. 18: $^{1}$H-$^{13}$C HMBC of lipoavitide 1 in acetone-$d_6$ focusing on the hdmHis region.**

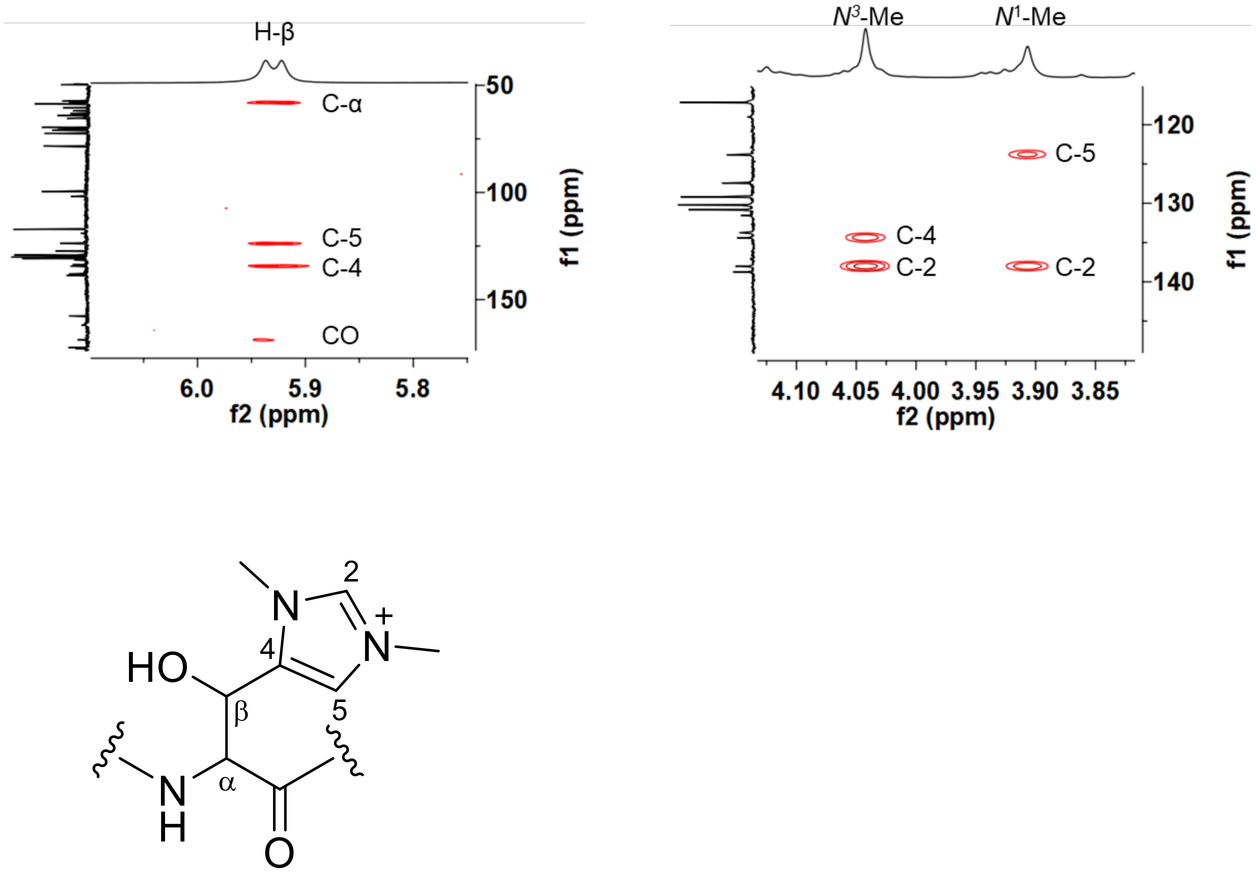

**Supplementary Fig. 19: $^{1}$H-$^{1}$H TOCSY of lipoavitide 1**.

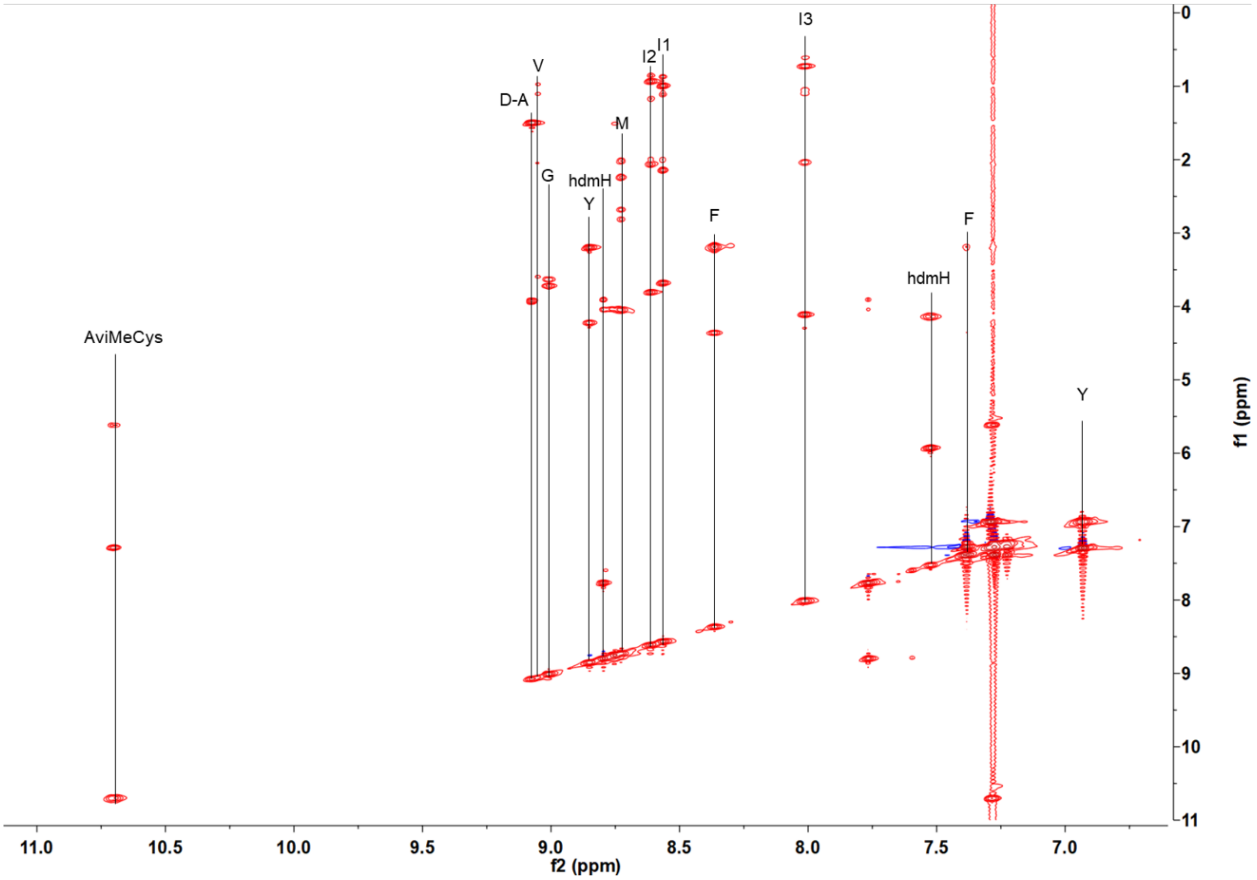

**Supplementary Fig. 20: Determination of amino acid stereochemistry in 1 using LC-MS**. Ion count-normalized LC-MS chromatograms are shown for derivatized amino acid standards and for 1-fluoro-2-4-dinitrophenyl-5-alanine amide (FDAA)-derivatized hydrolysate of **1**. *L*- and *D*-amino acid standards were run individually.

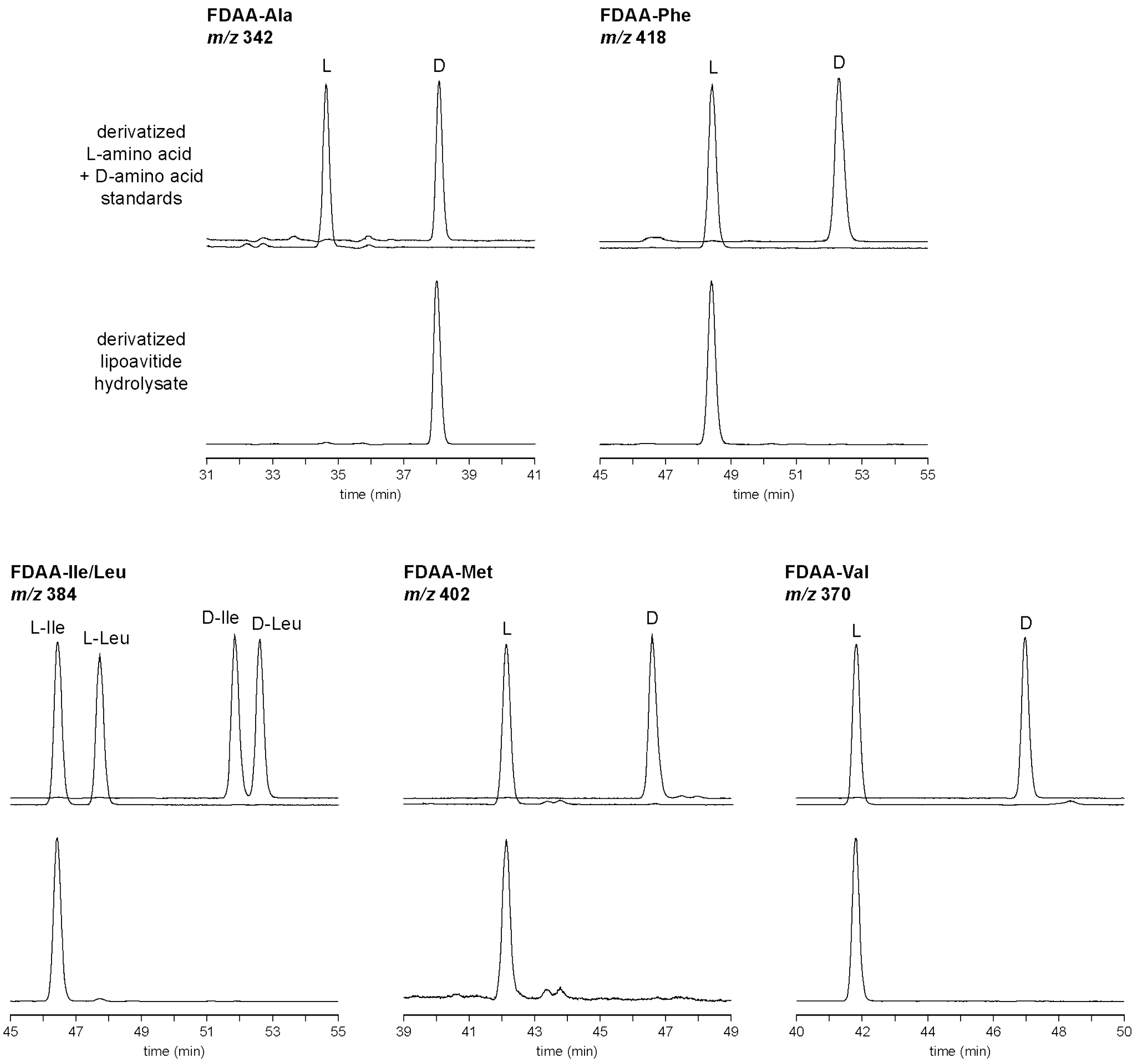

**Supplementary Fig. 21: Mild acid hydrolysis of 1**. (**a**) HPLC chromatogram of the reaction mixture after 20 h at 80 °C. (**b**) MALDI-TOF mass spectrometry analysis of the hydrolytic products. (**c**) Proposed structures of **2** and **3**.

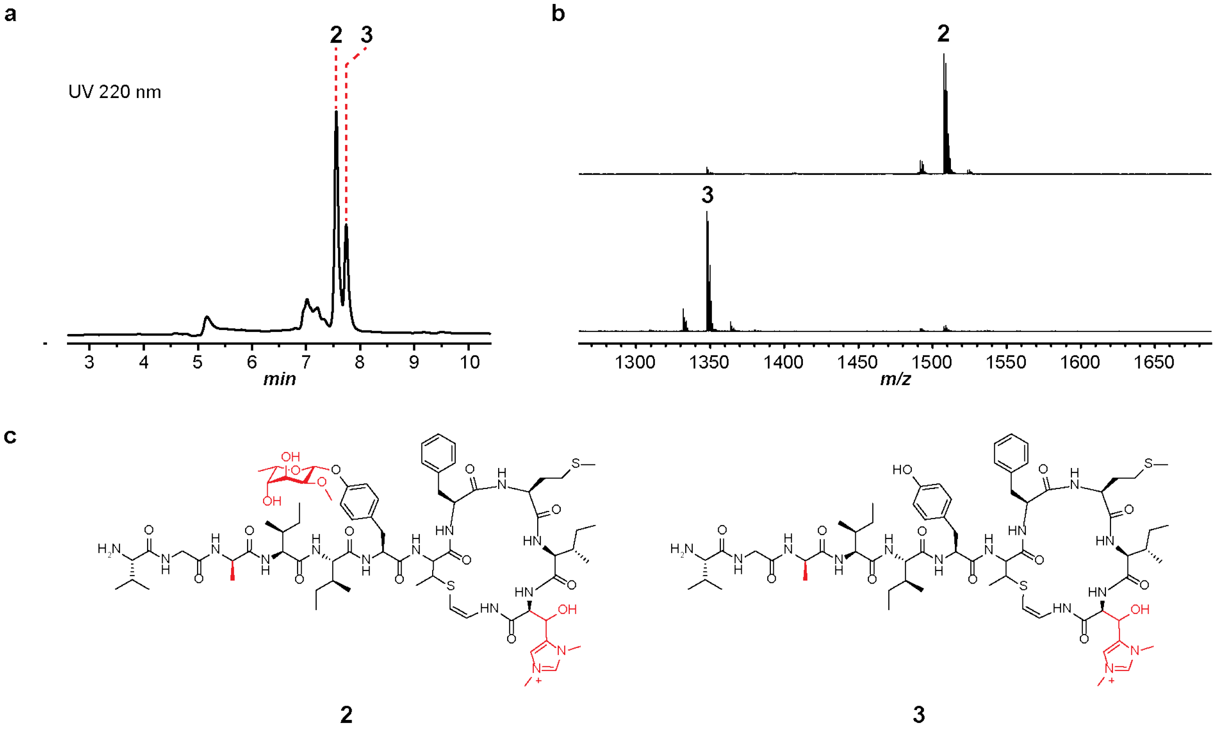

**Supplementary Fig. 22: HR-MS of product 2 and 3**. (**a**) Broad band high-resolution mass spectra of peptide **2** in +1 and +2 charged states. (**b**) Broad band high-resolution mass spectra of peptide **3** in +1 and +2 charged states.

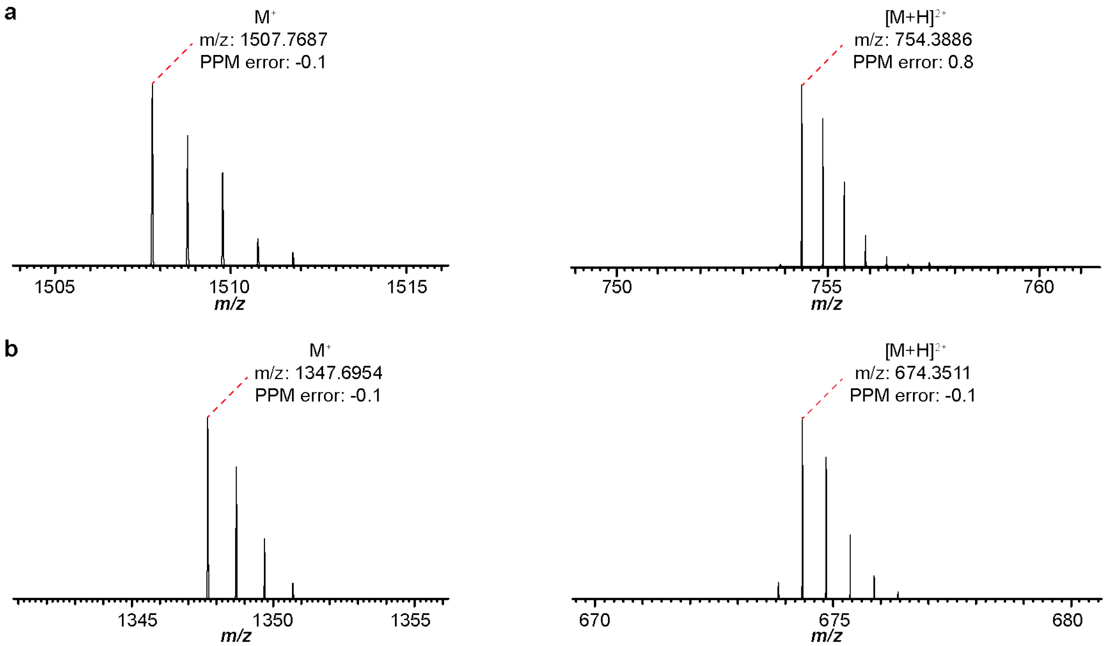

**Supplementary Fig. 23: $^1$H NMR spectrum of product 2 in acetone-$d_6$.** (**a**) $^1$H NMR spectrum of product **2** in comparison with **1**. (**b**) $^1$H NMR spectrum of product **2** in comparison with **1** focusing on the HMP region. Signals of Me-5 ($\delta_H$ = 1,31 ppm) and Me-7 ($\delta_H$ = 1.21 ppm) are absent in the spectrum of **2**.

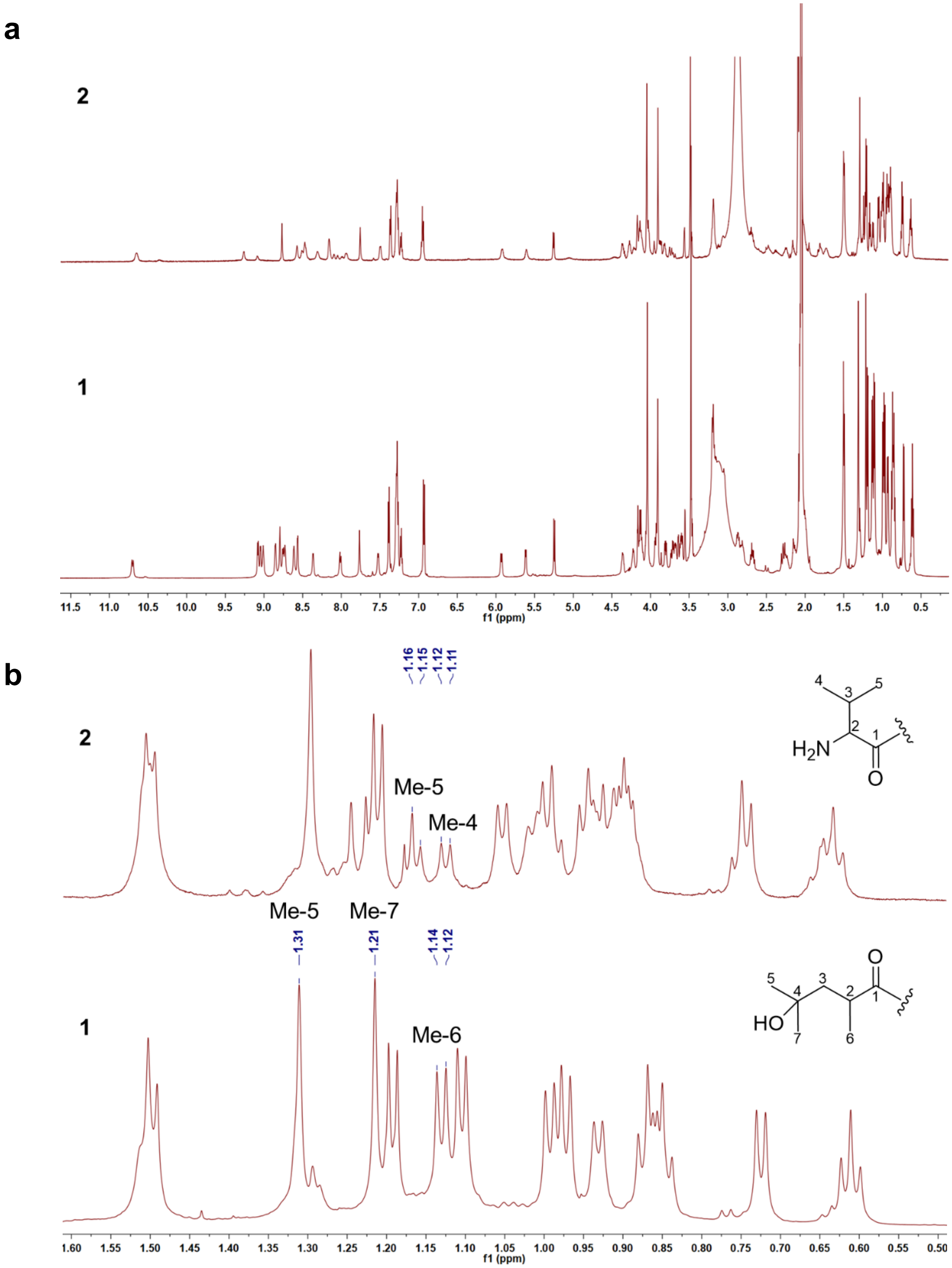

**Supplementary Fig. 24: $^1$H NMR spectrum of product 3 in acetone-$d_6$.** The $^1$H NMR spectrum of **1** is shown below for comparison. The signal of H-1’ in the 2-*O*-methyl-β-6-deoxygulosyl moiety ($\delta_H$ = 5.25 ppm) is absent in the spectrum of **3**.

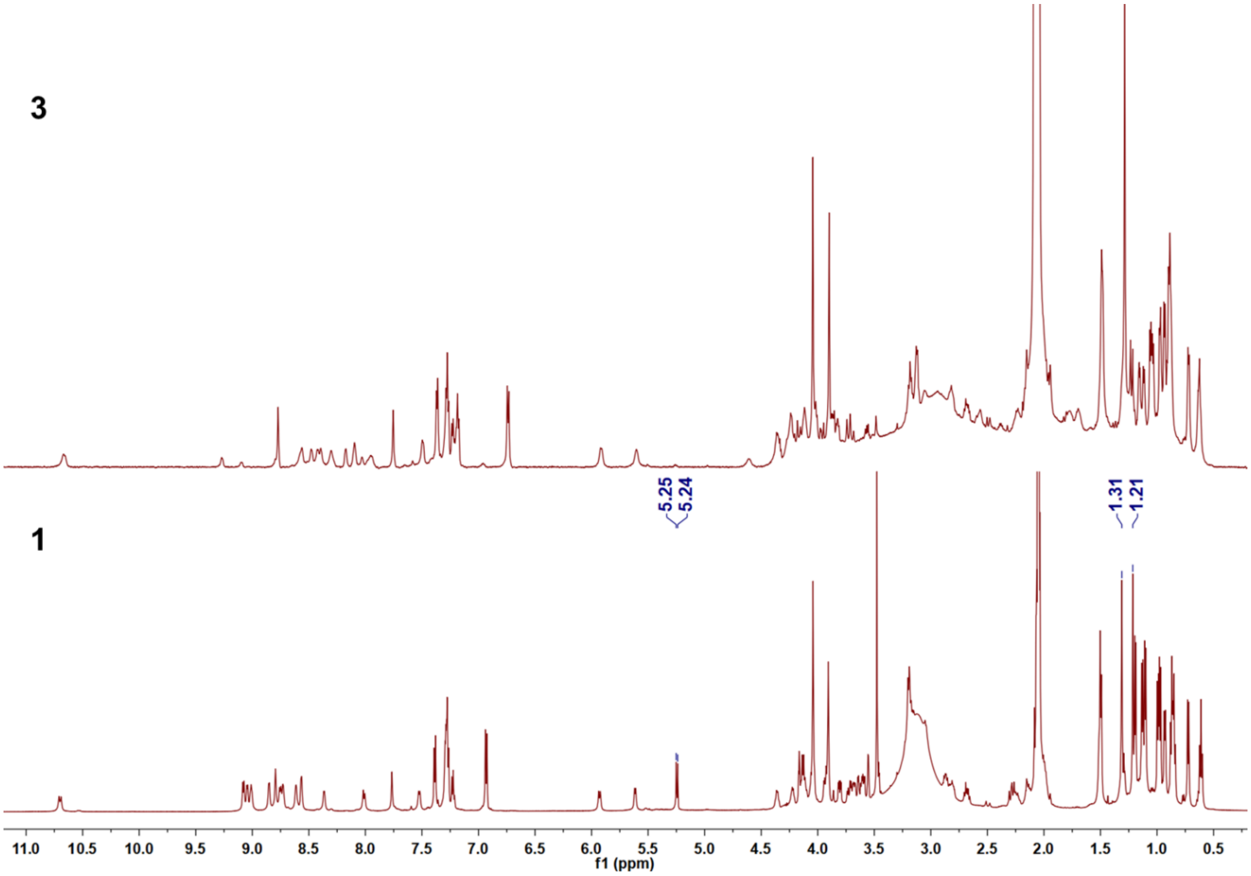

**Supplementary Fig. 25: Metabolism of valine and propionate in *S. albus*.** (**a**) Pathway that converts *L*-valine into isobutyryl-CoA. The isotopes in *L*-valine and its metabolites in the feeding study are highlighted by cyan circles. (**b**) Pathway that converts propionate into methylmalonyl-CoA. The isotopes in propionate and its metabolites in the feeding study are highlighted by cyan rectangles.

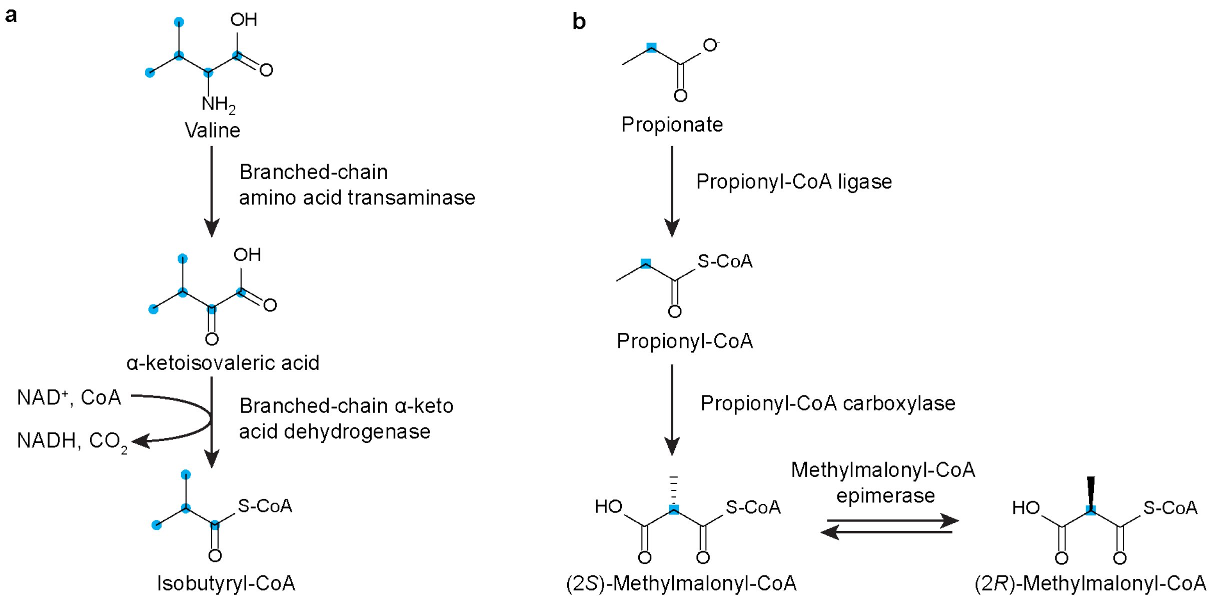

**Supplementary Fig. 26: MALDI-TOF mass spectrometry results of 1 purified from the isotopic labeling study**. (**a**) Product **1** produced in ISP4 solid medium and supplemented by [$^{13}C_5$] *L*-valine. Incorporation of carbon isotopes to the HMP, valine residue, and both positions result in +4 Da, +5 Da, and +9 Da shifts, respectively. (**b**) Product **1** produced in ISP4 solid medium supplemented by [$^{13}$C] propionate. Incorporation of carbon isotopes to the HMP results in +1 Da shift.

a

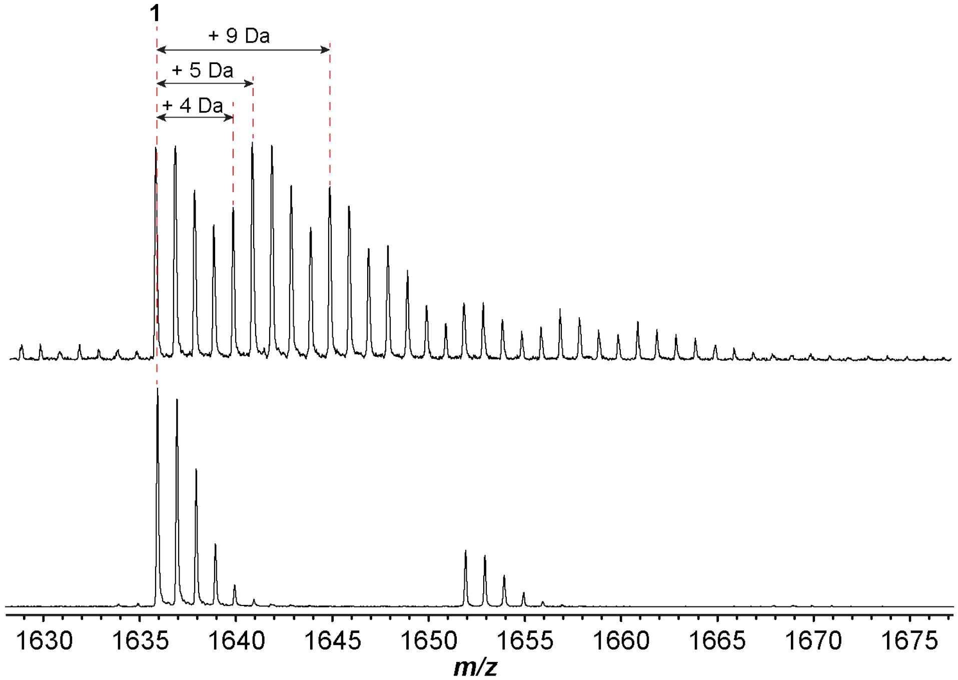

b

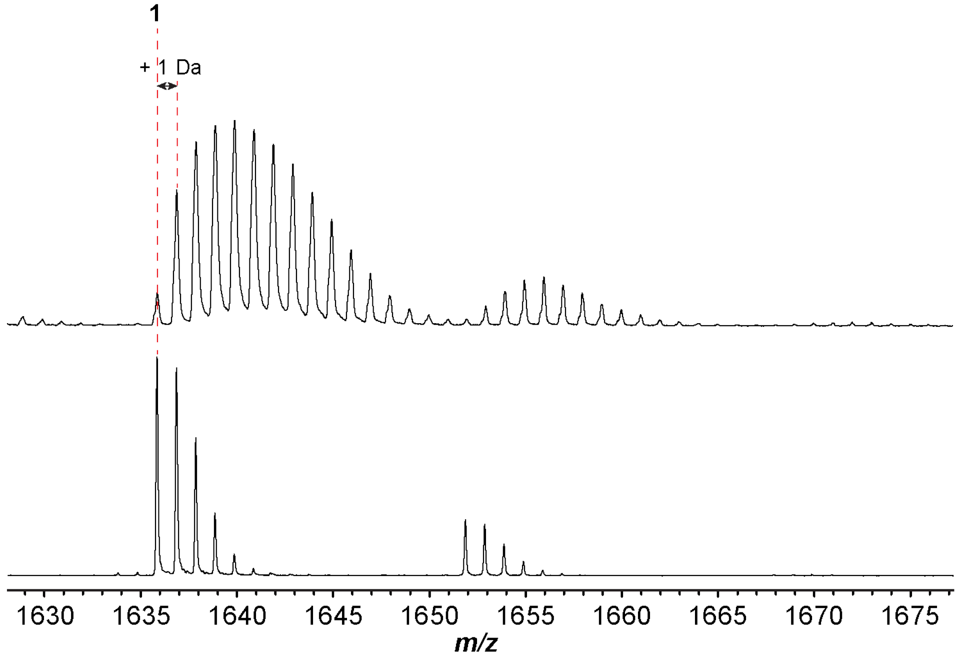

**Supplementary Fig. 27: Isotopic labeling of lipoavitide 1.** (**a**) $^{13}C$ NMR spectra of **1** labeled by [$^{13}C$] *L*-valine. (**b**) $^{13}C$ NMR spectra of **1** labeled by [$^{13}C$] propionate. Spectra of unlabeled **1** are shown in black for comparison. Enriched $^{13}C$ signals are indicated in the spectra and assigned to the positions in HMP and the valine residue.

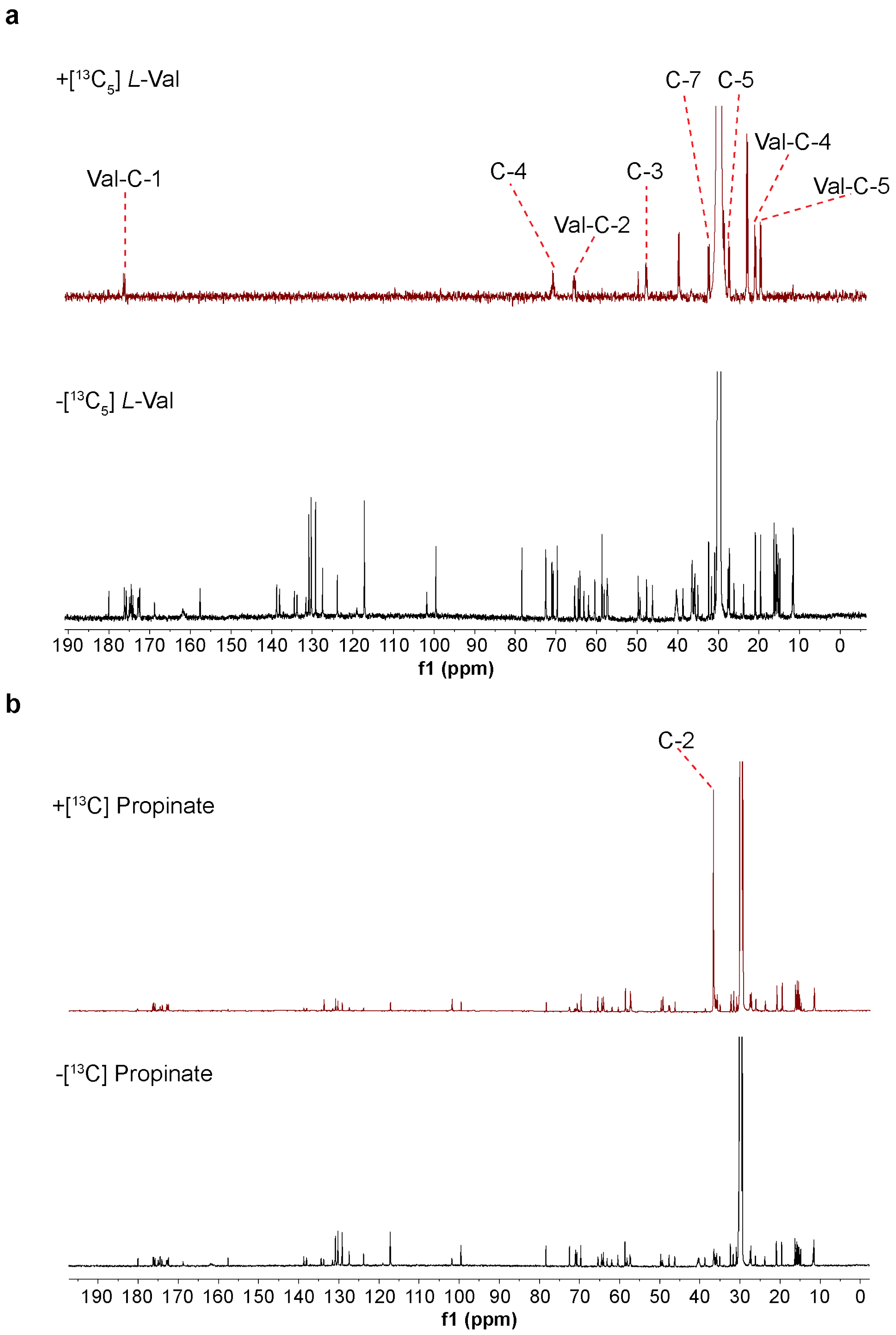

**Supplementary Fig. 28: SDS-PAGE analysis of native Ni-NTA purified proteins used in this study.** (**a**) StsG. (**b**) LpvV. (**c**) LpvE. (**d**) StsE. Composition of buffers for **a**-**c**: buffer A: 50 mM Sodium phosphate, 300 mM NaCl, 25 mM imidazole, pH 7.5; buffer B: 50 mM Sodium phosphate, 300 mM NaCl, 250 mM imidazole, pH 7.5. Composition of buffers for preparing **d**: buffer A: 50 mM Sodium phosphate, 300 mM NaCl, 25 mM imidazole, pH 7.5; buffer B: 50 mM Sodium phosphate, 300 mM NaCl, 500 mM imidazole, pH 7.5. A step gradient with an increasing percentage of buffer B was used for elution in 15 mL volume. Eluted fractions are denoted by the percentage of buffer B. Fractions collected for *in vitro* reactions are denoted by '*'. The ladder lanes are denoted by "#".

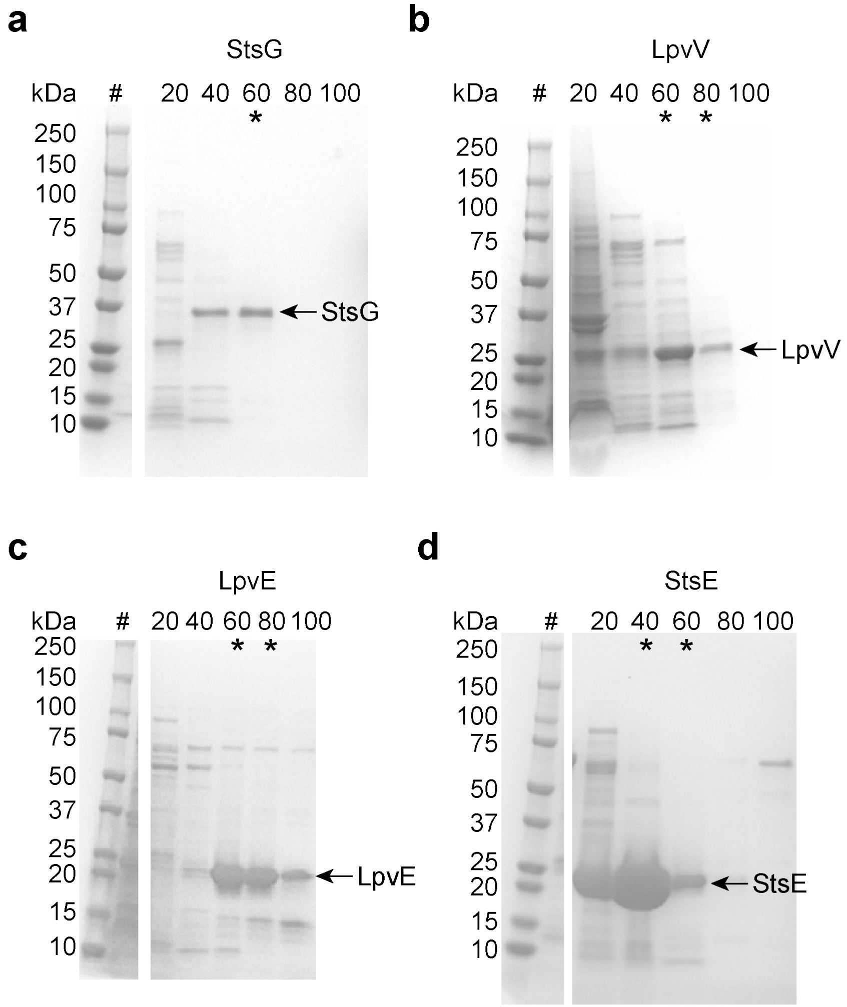

**Supplementary Fig. 29: HPLC chromatograms of the StsG-catalyzed condensation reaction under different substrate ratios**. (**a**) isobutyryl-CoA **5** : methylmalonyl-CoA **6** = 1:2. (**b**) isobutyryl-CoA : methylmalonyl-CoA = 1:1. (**c**) isobutyryl-CoA : methylmalonyl-CoA = 2:1. Absorbance was monitored at UV 260 nm.

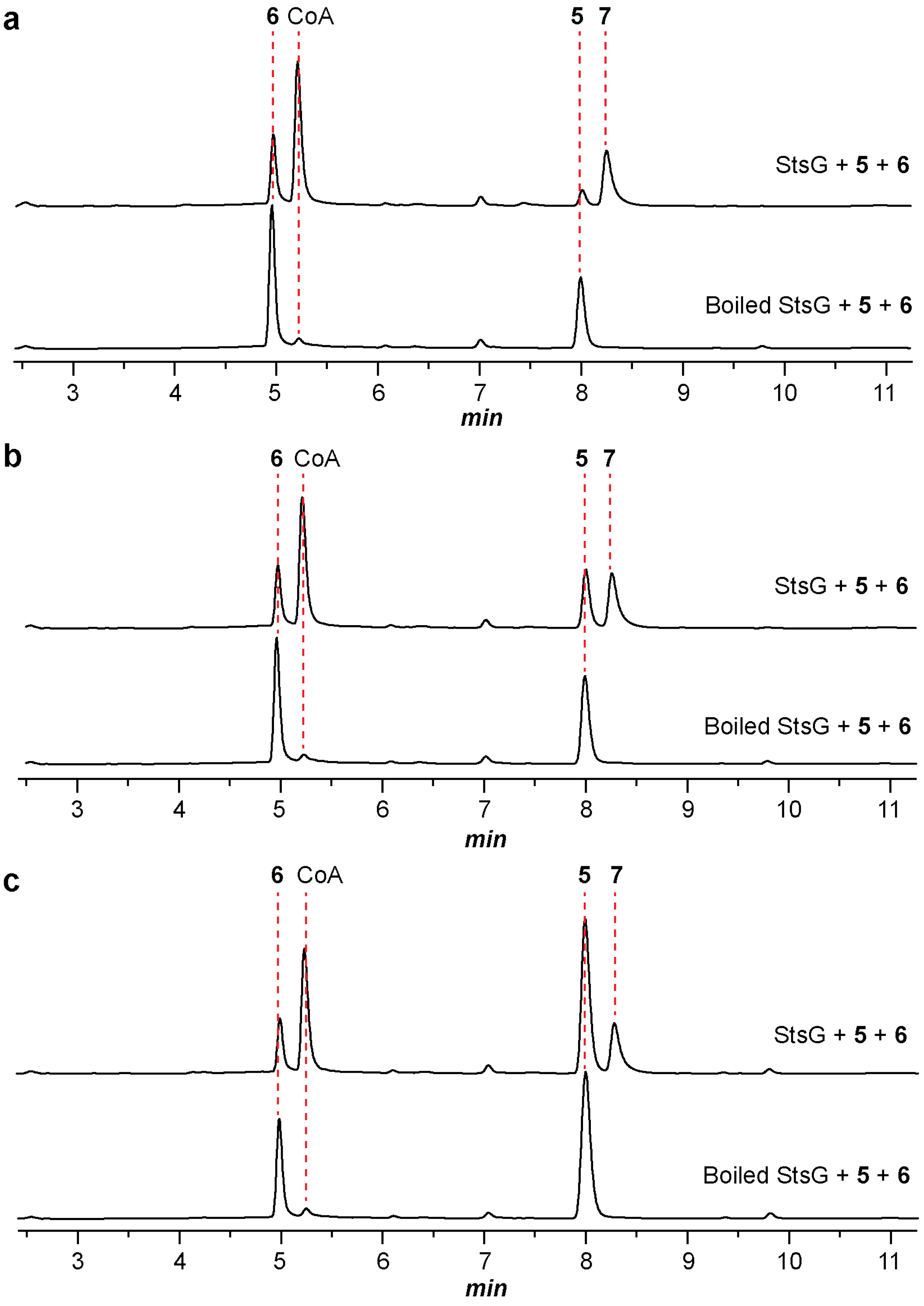

**Supplementary Fig. 30: HR-MS of compound 7**.

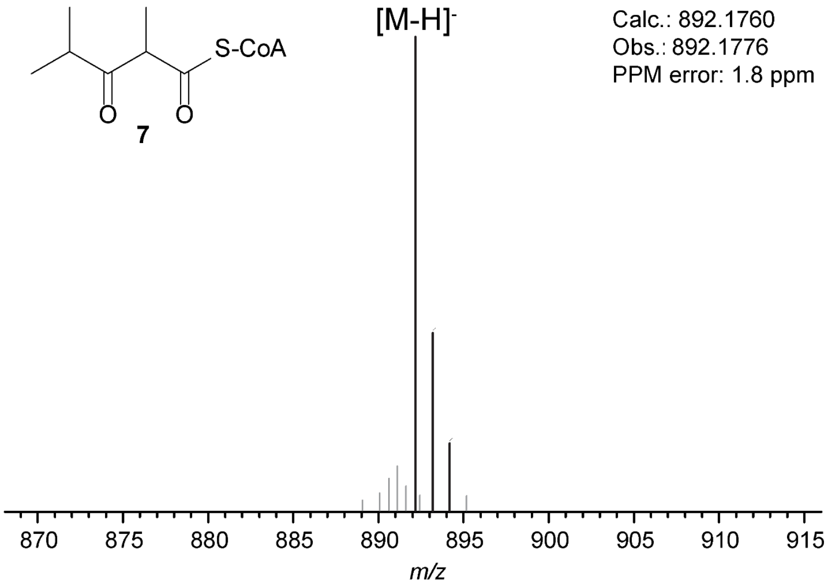

**Supplementary Fig. 31: HPLC chromatograms of the StsG-catalyzed condensation reaction supplemented with methylmalonyl-CoA epimerase**. Absorbance was monitored at UV 260 nm.

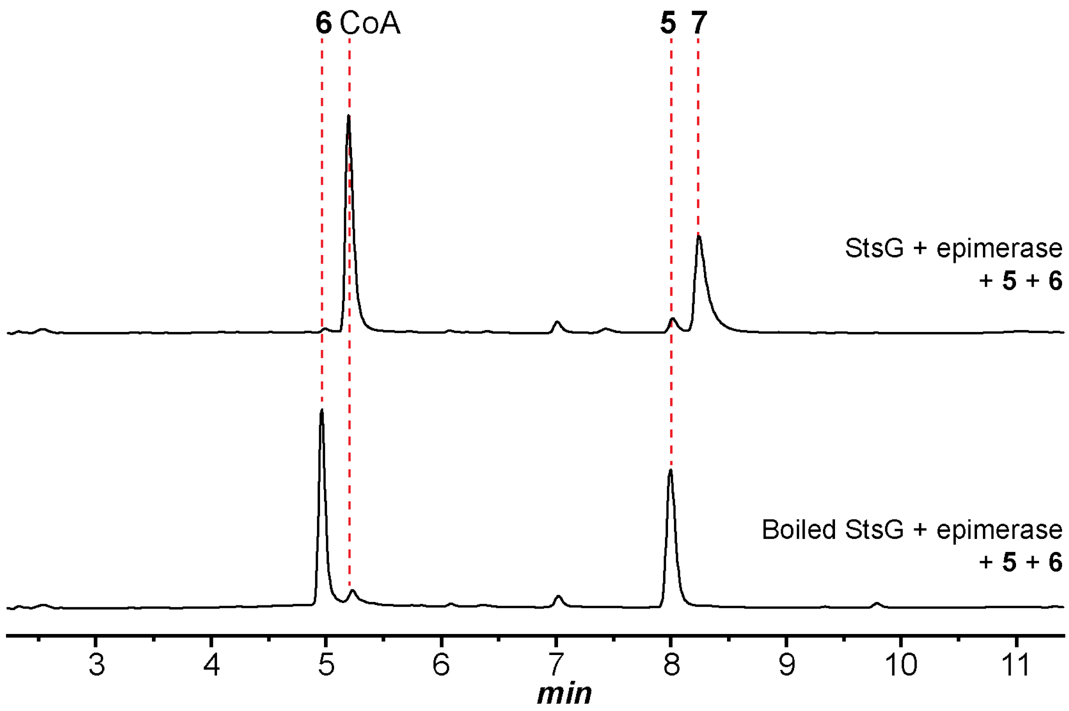

**Supplementary Fig. 32: Product analysis of *S. albus* J1074 containing the *lpvV*-deleted BGC**. (**a**) HPLC chromatogram of fractionated methanol extract. Absorbance was monitored at UV 220 nm. Results of the wild type *lpv* BGC is shown for comparison. (**b**) MALDI-TOF mass spectrum of the methanol extract. Observed *m/z* = 1636.

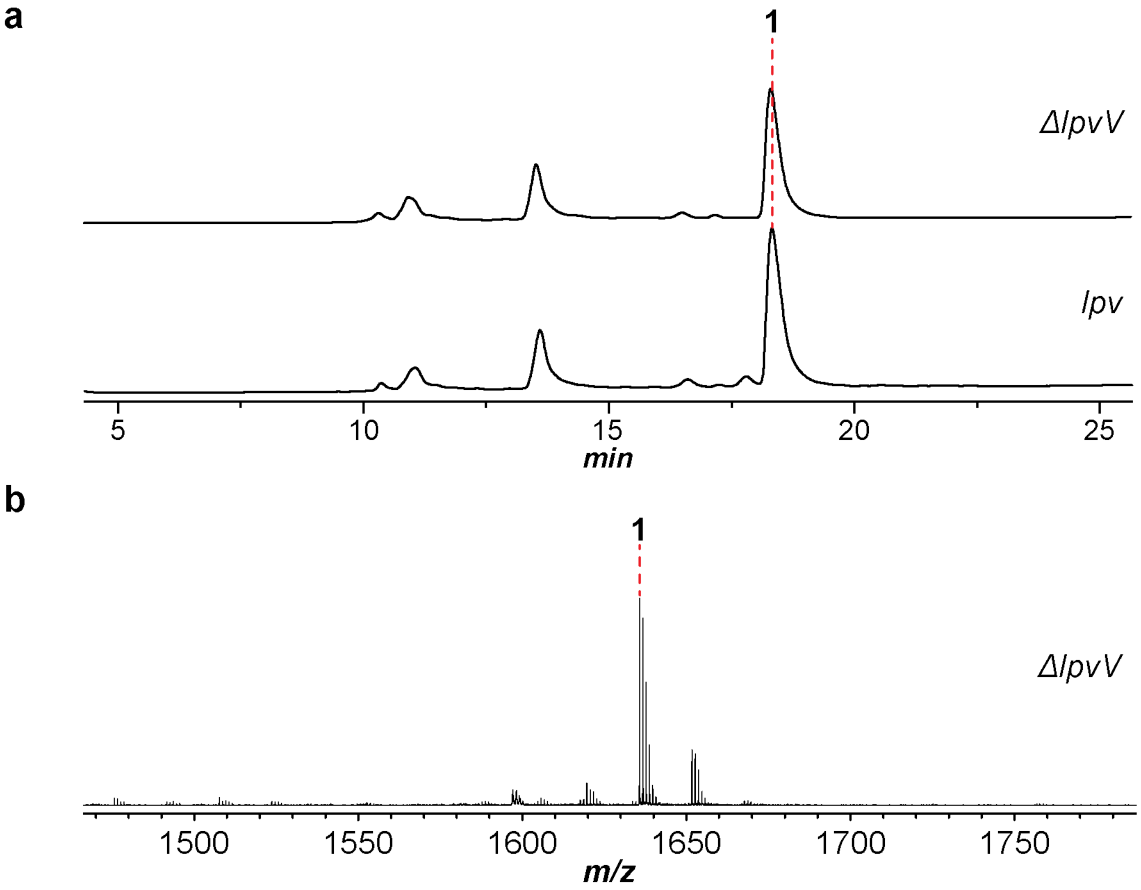

**Supplementary Fig. 33: HR-MS of compound 8**.

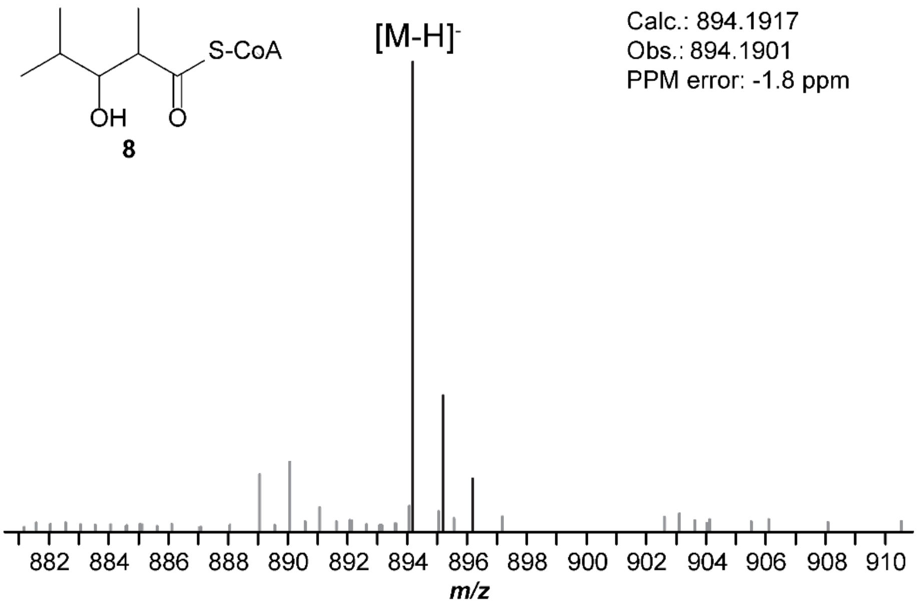

**Supplementary Fig. 34: Proposed pathway that converts 8 to 9**. LpvF was predicted to catalyze the dehydration reaction, which contains a SnoaL-like domain (PF12680), a family of proteins that exhibit diverse activity. BaiE (Uniprot ID: P19412), a characterized member in this family, is a bile acid 7α-dehydratase in the bile acid 7α-dehydroxylation pathway. LpvH was predicted as an enoylreductase. LpvS was predicted as a P450 monooxygenase to catalyze γ-hydroxylation.

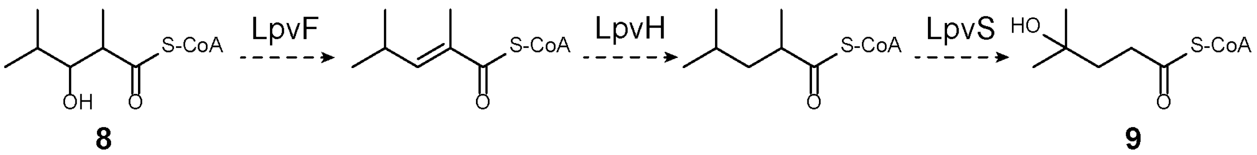

**Supplementary Fig. 35: Product analysis of *S. albus* J1074 containing the *lpvF*-, *lpvH*-, or *lpvS*-deleted BGC**. (**a**) HPLC chromatograms of fractionated methanol extracts. Absorbance was monitored at UV 220 nm. Results of the wild type *lpv* BGC is shown for comparison. (**b**) MALDI-TOF mass spectra of the methanol extracts. Observed *m/z* = 1636.

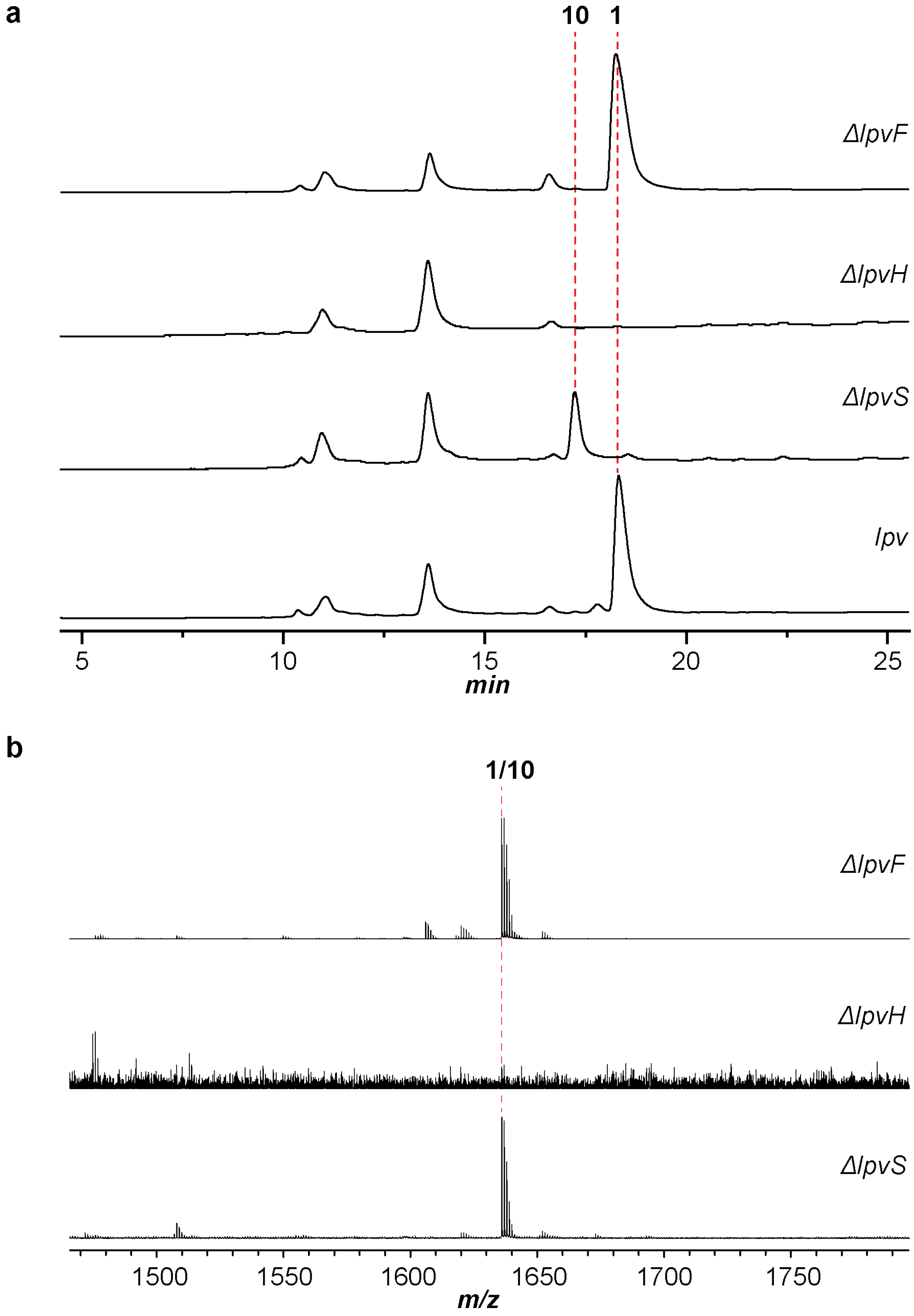

**Supplementary Fig. 36: HR-MS of peptide 10**. *Left*, +1 charge state. *Right*, +2 charge state.

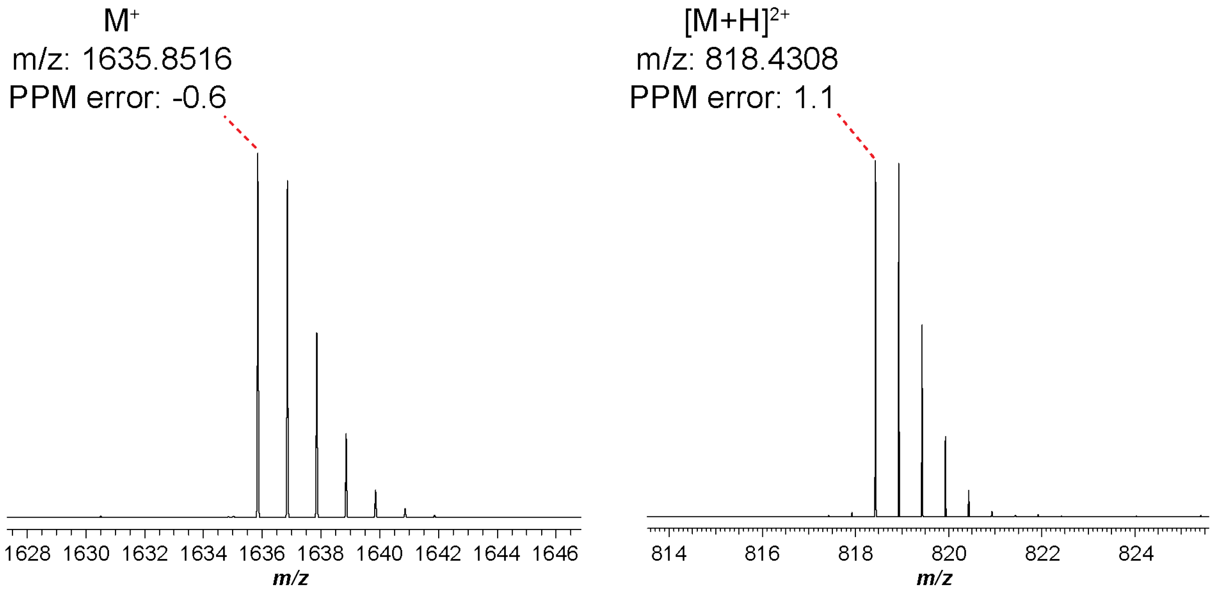

**Supplementary Fig. 37: NMR spectra of peptide 10**. (**a**) $^{1}$H NMR spectrum of peptide **10** in acetone-$d_6$. The $^{1}$H NMR spectrum of **1** is shown below for comparison. (**b**) Chemical shifts of Me-5, Me-6, and Me-7 in **10** and **1**. Of note, compared with Me-5 in **1** which is a singlet, Me-5 in **10** is a doublet due to the existence of H-4. (**c**) $^{1}$H-$^{1}$H COSY of **10**. (**d**) $^{1}$H-$^{1}$H COSY of **10** focusing on the 3-HMP region.

**a**

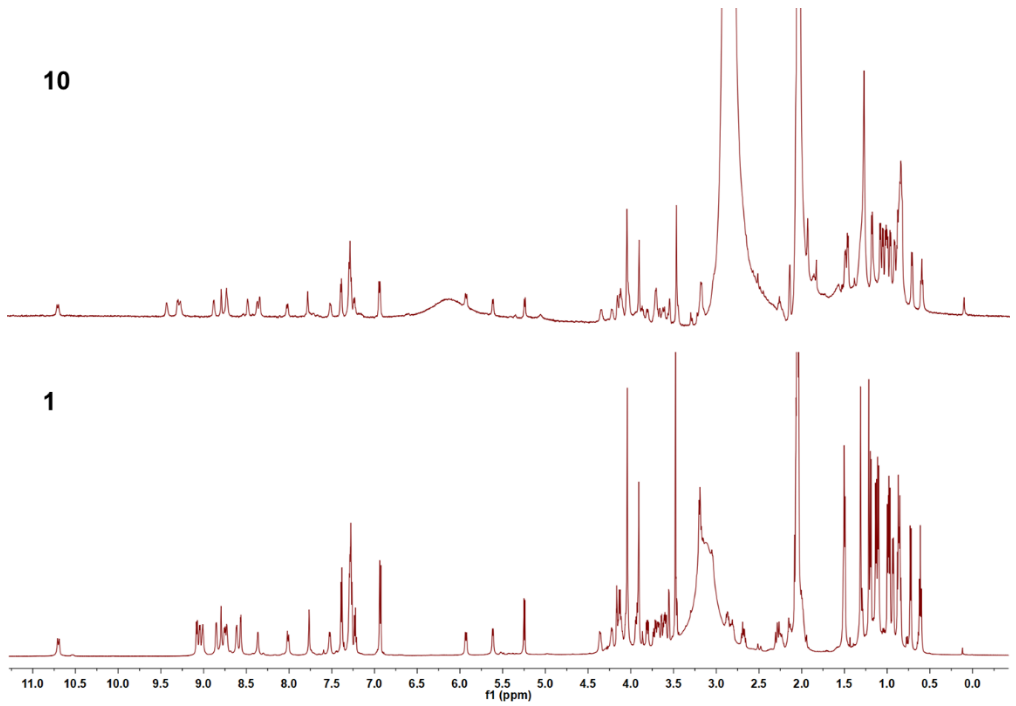

**b**

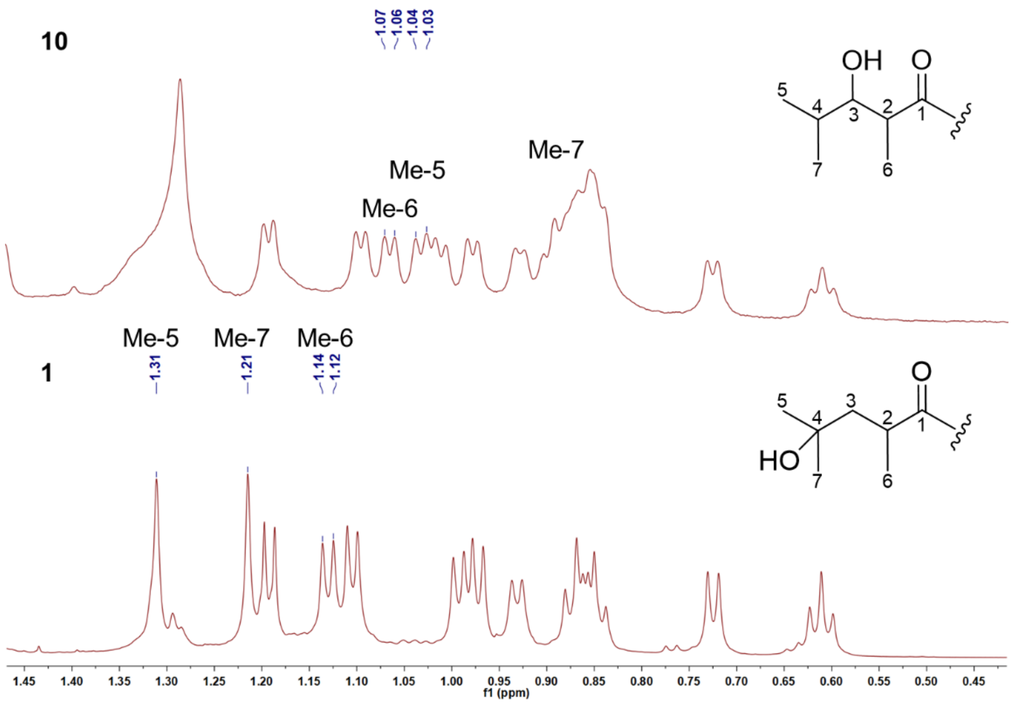

**c**

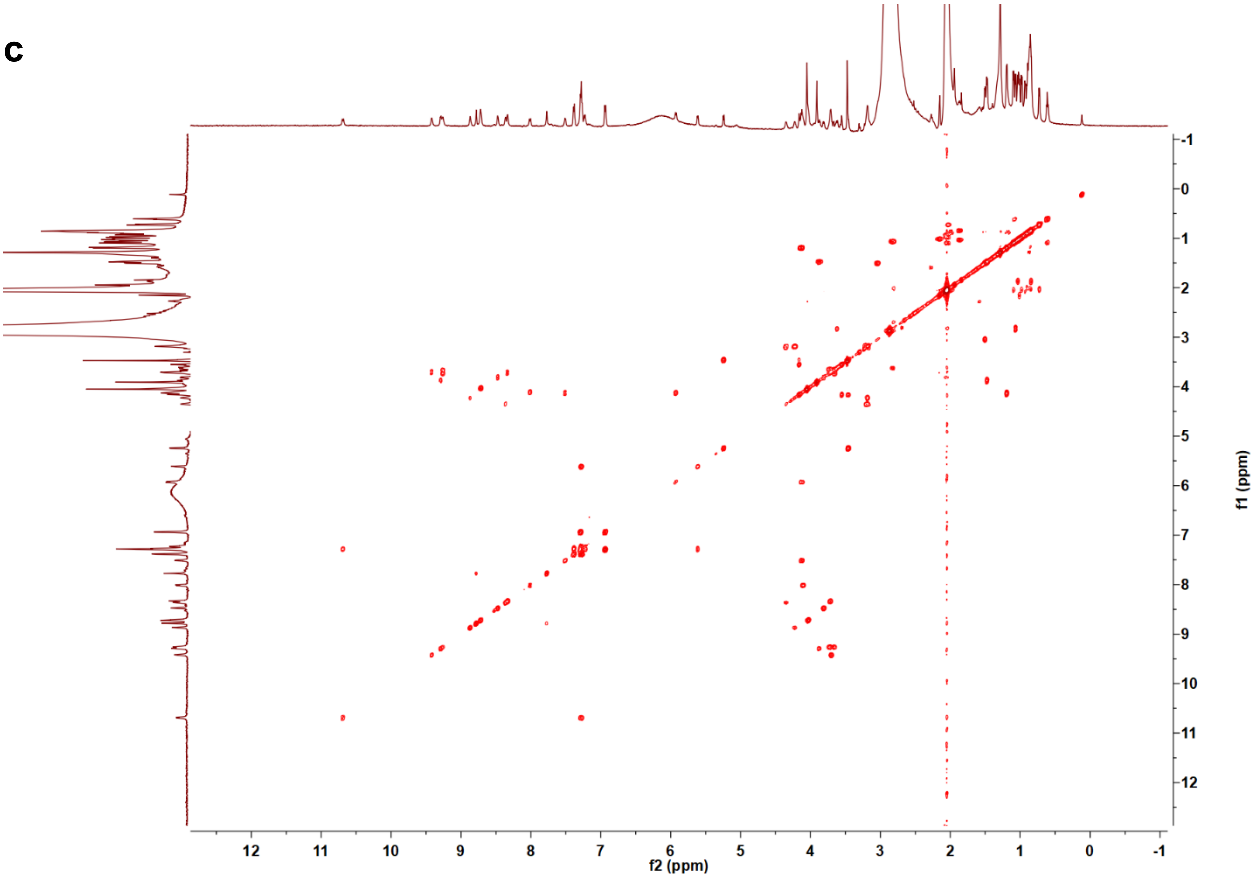
f2 (ppm)
f1 (ppm)
12
11
10
9
8
7
6
5
4
3
2
1
0
-1

**d**

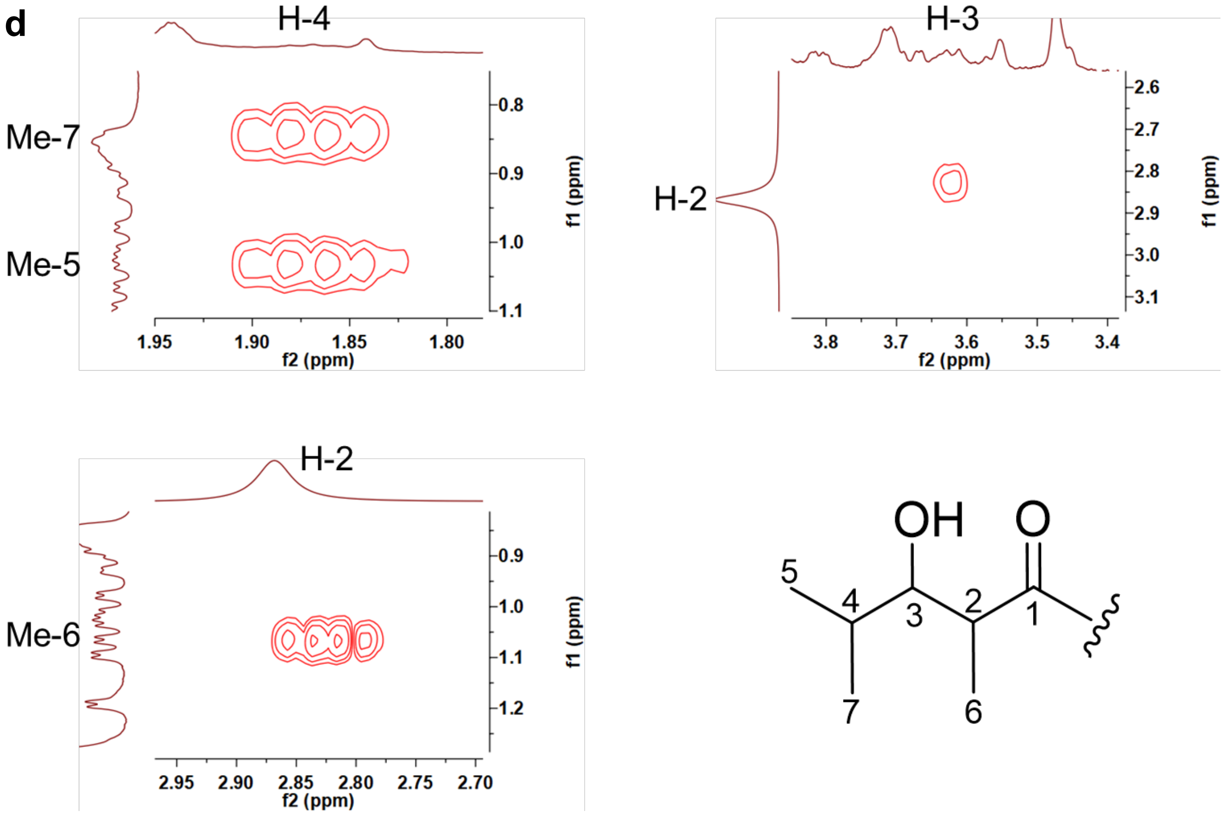
H-4
Me-7
Me-5
1.95
1.90
1.85
1.80
0.8
0.9
1.0
1.1
f2 (ppm)
f1 (ppm)
H-3
H-2
3.8
3.7
3.6
3.5
3.4
2.6
2.7
2.8
2.9
3.0
3.1
H-2
Me-6
2.95
2.90
2.85
2.80
2.75
2.70
0.9
1.0
1.1
1.2
OH
O
5
4
3
2
1
7
6

**Supplementary Fig. 38: HR-MS/MS analysis of 10 formed by *in vitro* reaction**. (**a**) Broad band high-resolution mass spectrum of compound **10**. (**b**) Summary of collision-induced dissociation (CID) daughter ions for **1**. All ions are in the +1 charge state. "Retro aldol" indicates the formation of 4-formyl-1,3-dimethyl-1*H*-imidazolium through retro aldol reaction during MS/MS analysis. (**c**) CID spectrum of **1**. (**d**) Enlarged CID spectrum of **1** in the dashed box region. (**e**) Table of daughter ion assignments for **10**.

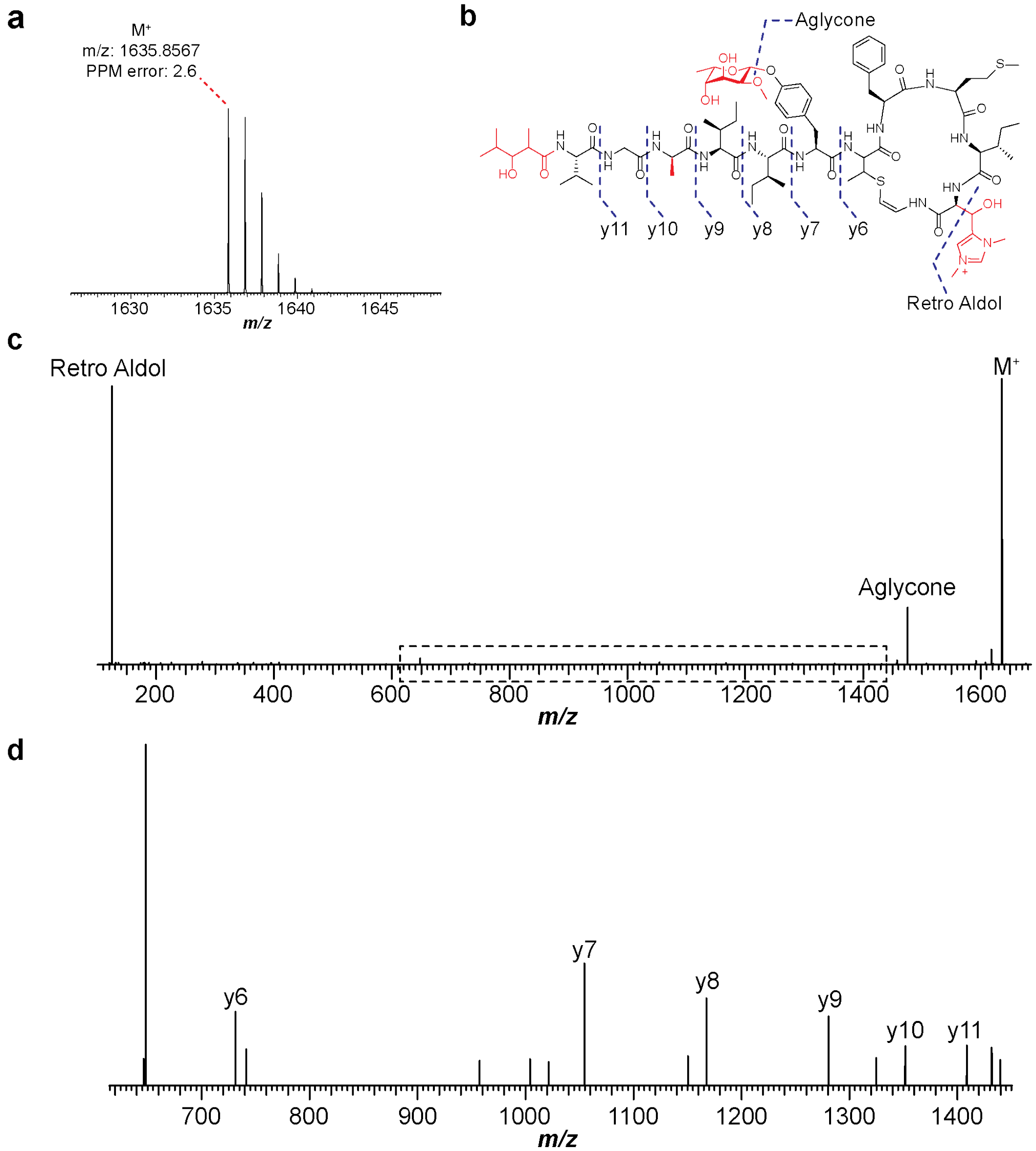

e

| Ion | Calc. Mass | Obs. Mass | Ppm Error |
|---|---|---|---|
| **$M^+$** | 1635.8525 | 1635.8585 | 3.7 |
| **Aglycone** | 1475.7789 | 1475.7841 | 3.5 |
| **Retro Aldol** | 125.0709 | 125.0716 | 5.6 |
| **y6** | 731.3368 | 731.3418 | 6.8 |
| **y7** | 1054.4736 | 1054.4797 | 5.8 |
| **y8** | 1167.5577 | 1167.5627 | 4.3 |
| **y9** | 1280.6418 | 1280.6431 | 1.0 |
| **y10** | 1351.6789 | 1351.6833 | 3.3 |
| **y11** | 1408.7003 | 1408.7042 | 2.8 |

**Supplementary Fig. 39: Enzymatic reconstitution of StsG/LpvV/LpvE**. MALDI-TOF mass spectra of reactions with all the three enzymes (StsG: FabH; LpvV: short-chain dehydrogenase/reductase; LpvE: acyltransferase) and each of them omitted. The isobutylated product (**11**) was observed at *m/z* 1578. The 3-oxo-2,4-dimethylpentanoylated product (**12**) was observed at *m/z* 1634. The region in the dashed box is enlarged for comparing the difference between **12** and **10**.

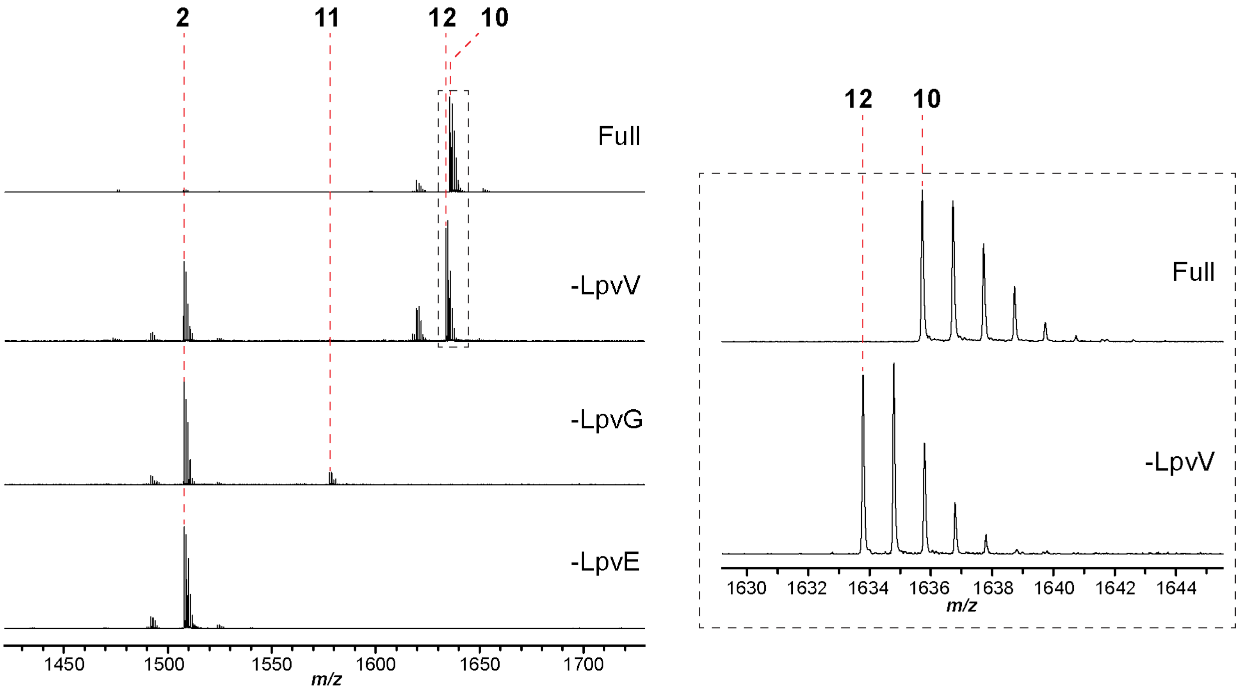

**Supplementary Fig. 40: Synthesis and purification of 2,4-dimethylpentanoyl-CoA using the CDI activation approach**. (**a**) Scheme of the two-step synthesis. (**b**) HPLC traces of the reaction mixtures. Absorbance was monitored at UV 260 nm. The result of the control reaction without CDI is shown below for comparison. (**c**) HR-MS of 2,4-dimethylpentanoyl-CoA.

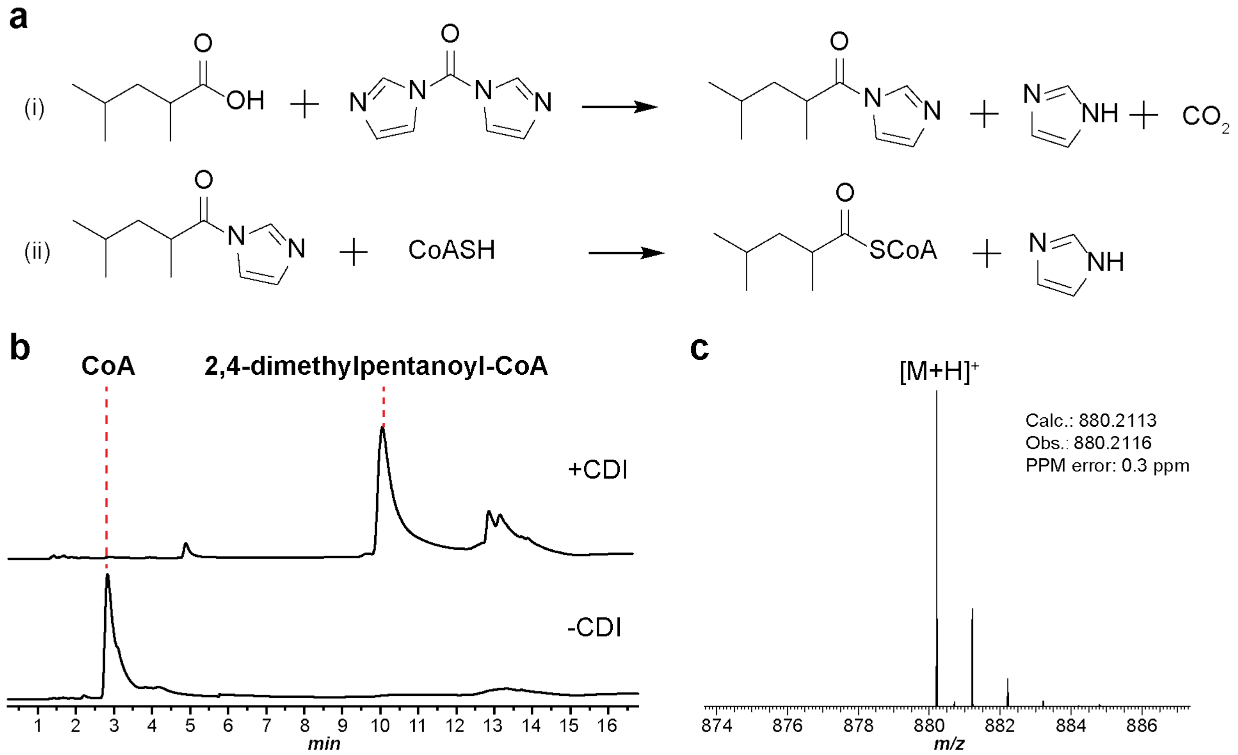

**Supplementary Fig. 41: MALDI-TOF mass spectra of the LpvE-catalyzed condensation reaction of 2,4-dimethylpentanoyl-CoA and peptide 2**. The acylated product **13** was observed at *m/z* 1620. Heat-inactivated LpvE did not yield a detectable quantity of **13**.

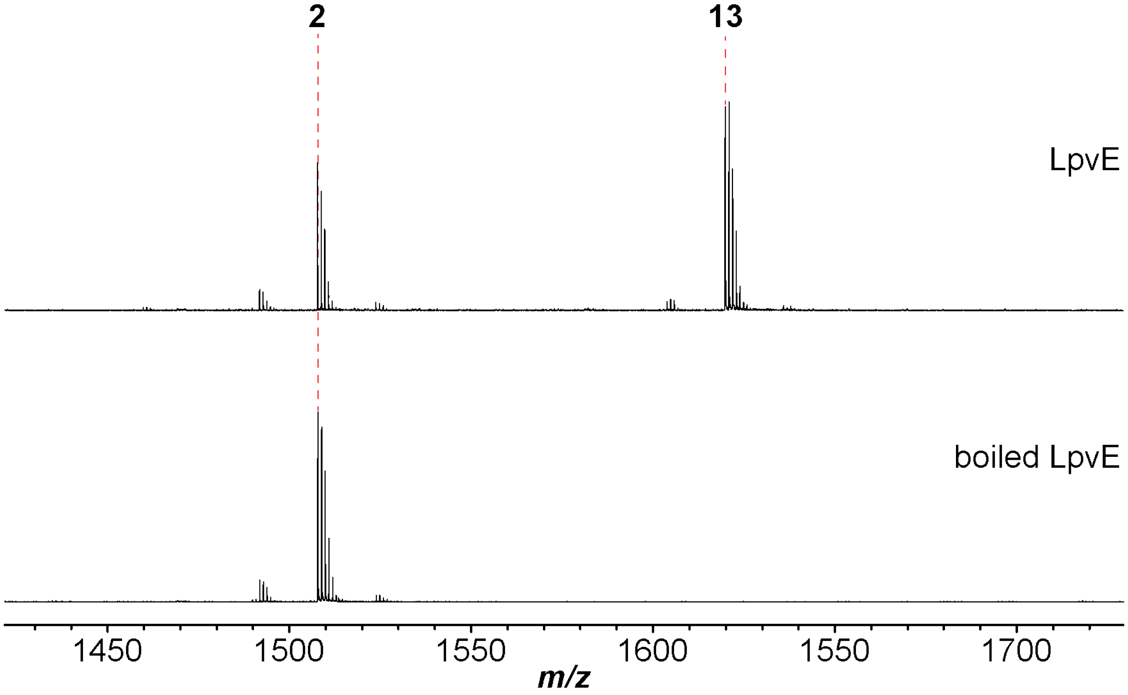

**Supplementary Fig. 42: MALDI-TOF mass spectra of methanol extracts of *S. albus* J1074 containing *lpv* substituted by LpvA variants V1A and V1K**. The processed V1A variant (**14**) was observed at *m/z* 1608. The V1K variant (expected at *m/z* 1693) was not observed.

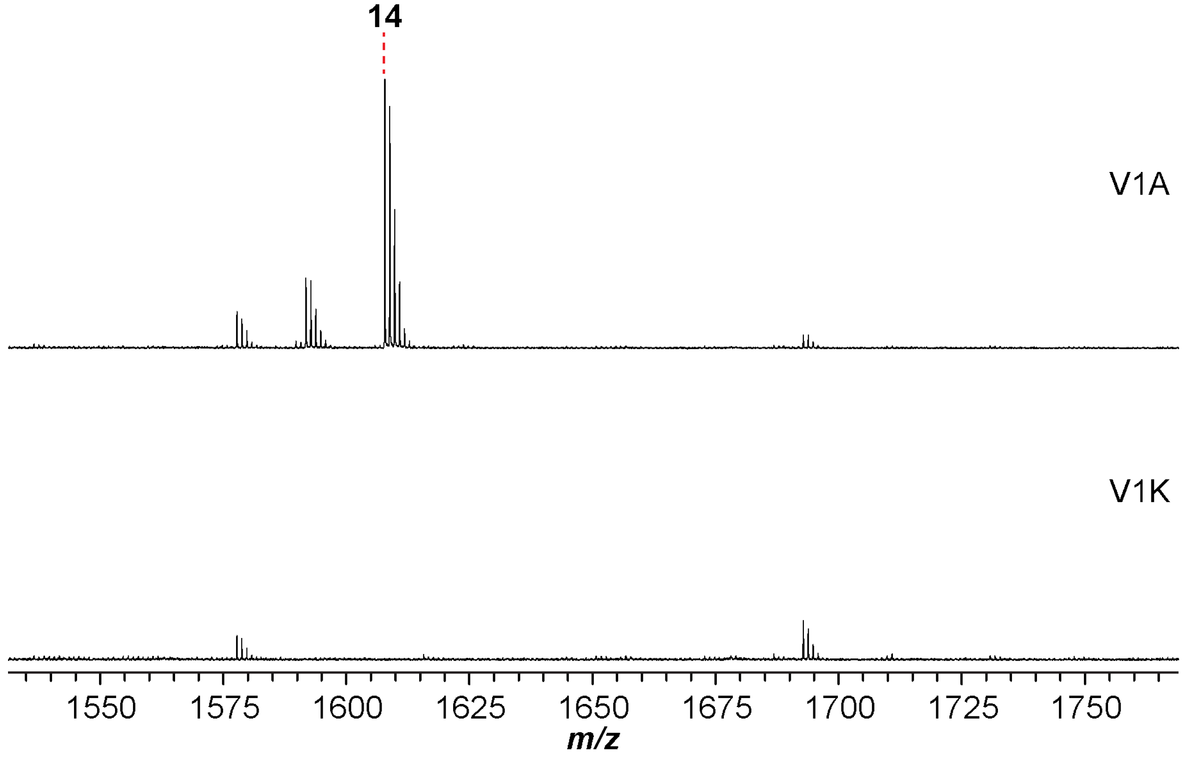

**Supplementary Fig. 43: MALDI-TOF mass spectra of methanol extracts of *S. albus* J1074 containing *lpv* with *lpvJ* deleted, *lpvA* substituted by S3A and S3G variants**. Products were observed at *m/z* 1634 (**15**), 1636 (**16**), and 1622 (**17**), respectively.

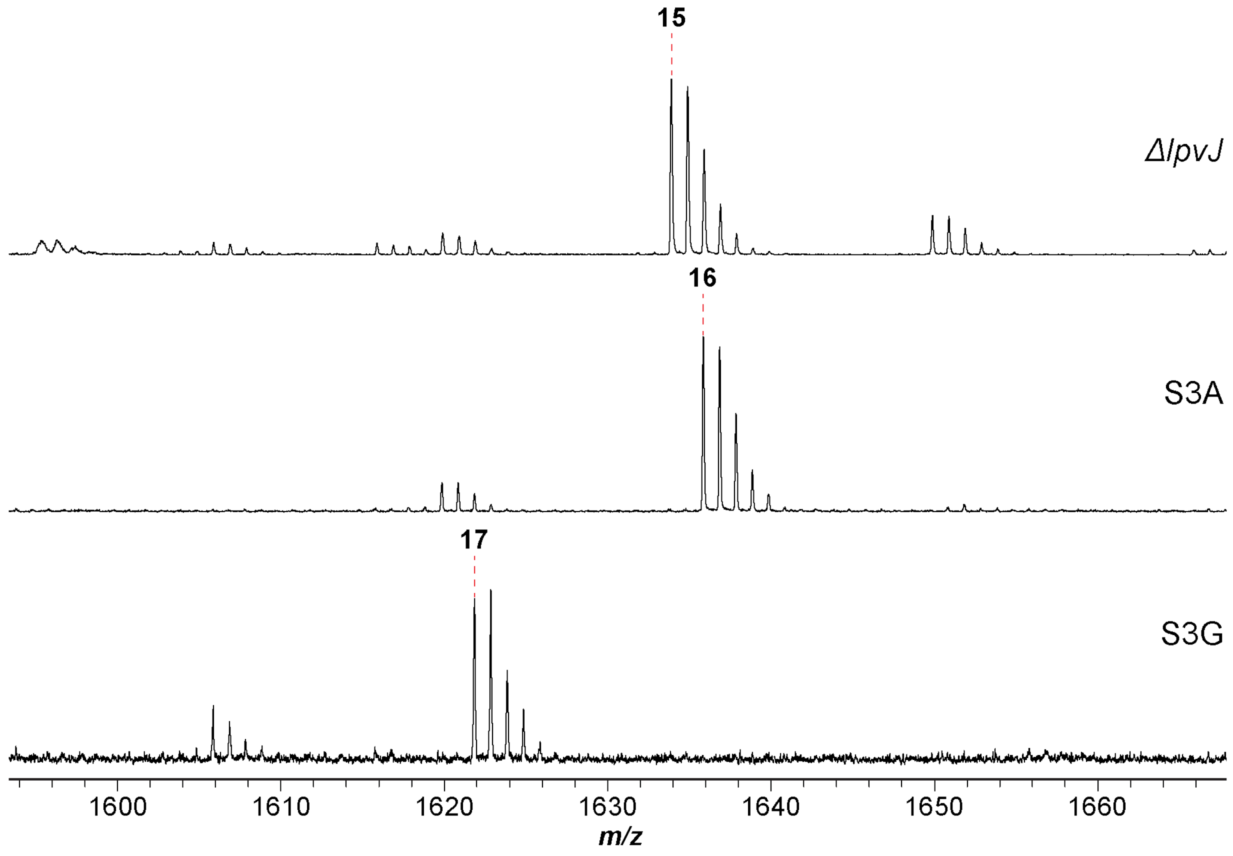

**Supplementary Fig. 44: MALDI-TOF mass spectra of methanol extracts of *S. albus* J1074 containing containing *lpv* with *lpvI* deleted**. The product of *lpvI*-deleted construct was observed at *m/z* 1476 (**18**). Products of *lpv* substituted by *lpvA* variants Y43F and Y43W were expected with *m/z* 1460 and 1499.

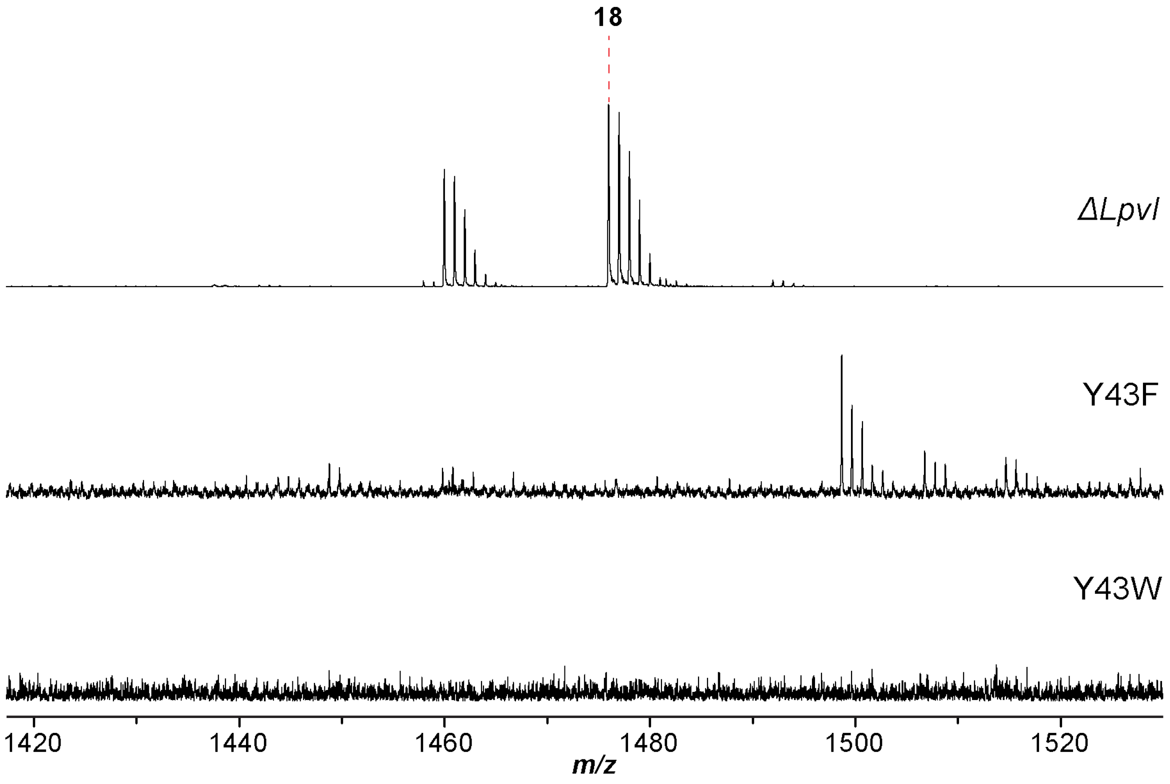

**Supplementary Fig. 45: Enzymatic reconstitution of StsG/LpvV/LpvE using 3 as the peptide substrate**. The acylated peptide (3-HMP-modified **3**) was observed at *m/z* 1476.

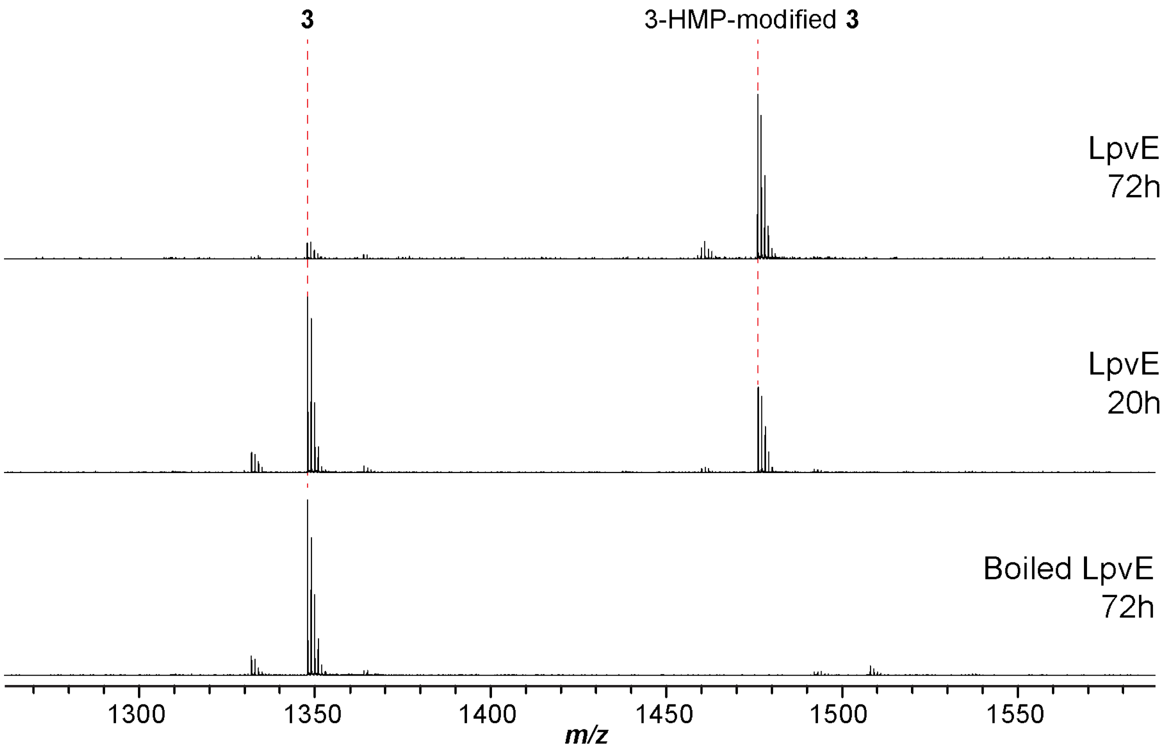

**Supplementary Fig. 46: MALDI-TOF mass spectra of methanol extracts of *S. albus* J1074 containing *lpvM1*- and *lpvK*-deleted BGC**. Products were observed at *m/z* 1620 (**19**) and 1592 (**20**), respectively.

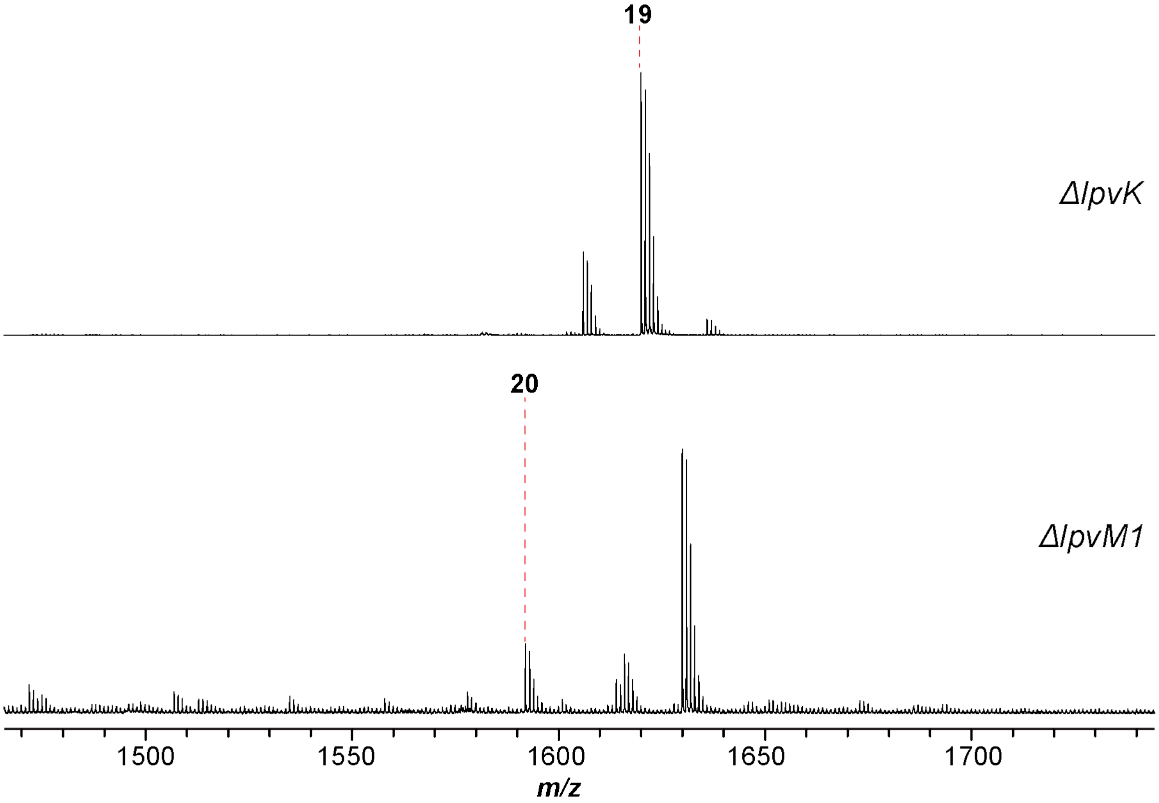

**Supplementary Fig. 47: Enzymatic reconstitution of StsG/LpvV/LpvE using linear peptide VG$^{D}$AIIYTFMIHC as the substrate**. The peptide substrate was observed at *m/z* 1368. The acylated product was expected at *m/z* 1496.

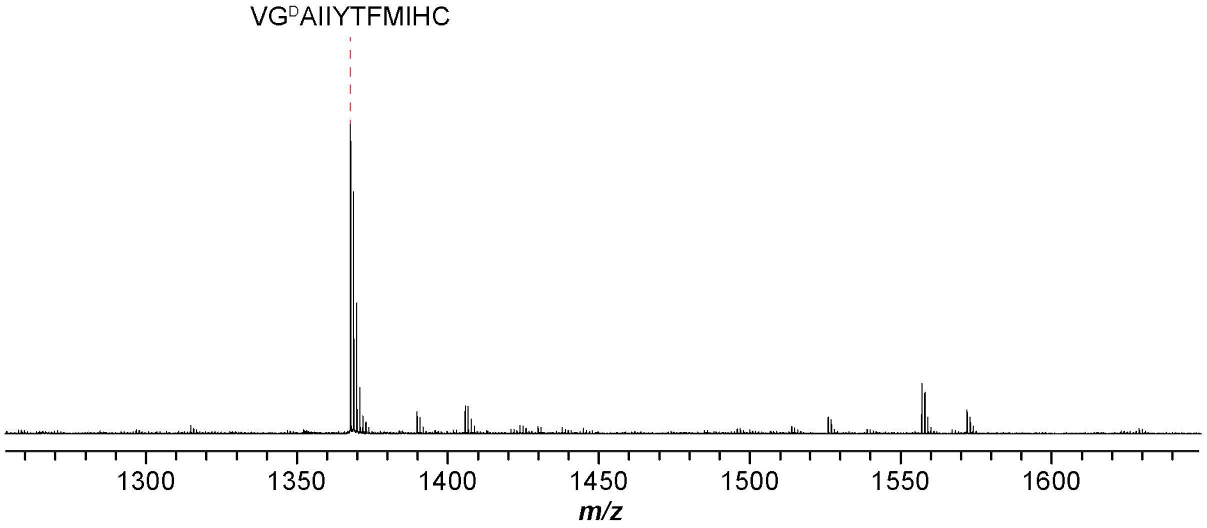

**Supplementary Fig. 48: AlphaFold2 simulation and molecular docking study of LpvE**. (**a**) The overall structure of LpvE (Uniprot ID: A0A4S3G691) in complex with **2** and **9**. The carbon atoms in **2** and **9** are colored in purple and grey, respectively. (**b**) Enlarged view focusing on the amino and HMP moieties in **2** and **9**, respectively. The distance between the *N*-terminal nitrogen in **2** and the carbonyl carbon of the thioester in **9** is 4.508 Å. (**c**) View of the substrate-binding pocket. The protein surface is colored by hydrophobicity (hydrophilic, blue; hydrophobic, red). (**d**) Structure superimposition of LpvE and GNAT acetyltransferase PA2578 (UniProt ID: Q9I0Q8; PDB ID: 3OWC). LpvE and PA2578 are shown in yellow and light blue, respectively. The stick representation of CoA in PA2578 overlaid with **9** of LpvE obtained in docking study. (**e**) The AlphaFold2 confidence measures for LpvE. The predicted Local Distance Difference Test (pLDDT) value is shown per-residue to indicate the quality of obtained structure.

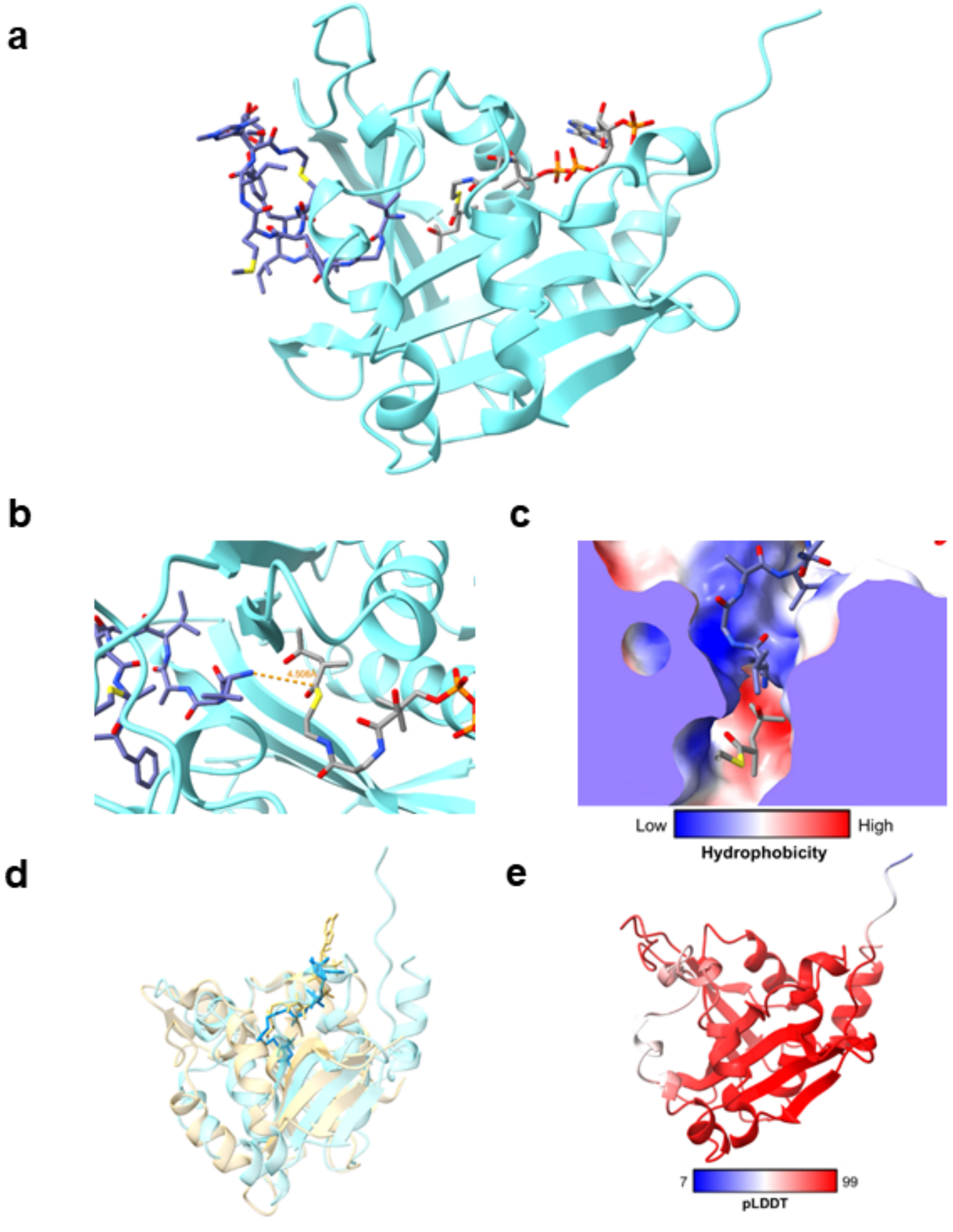

**Supplementary Fig. 49: Maximum-likelihood phylogenetic analysis of lipoavitide acyltransferases within the GNAT superfamily.** Acyltransferases for lipoavitide, lipolanthin, goadvionin, and selidamide biosynthesis are colored in cyan, blue, brown, and purple, respectively. For identical acyltransferases that were identified in multiple BGCs, only one representative is shown for clarity. The tree is rooted using (*S*)-1-pyrroline-5-carboxylate acetyltransferase (ScMpr1, UniProt/GenBank CDS identifiers: E9P8D2/ BAA95611.1).

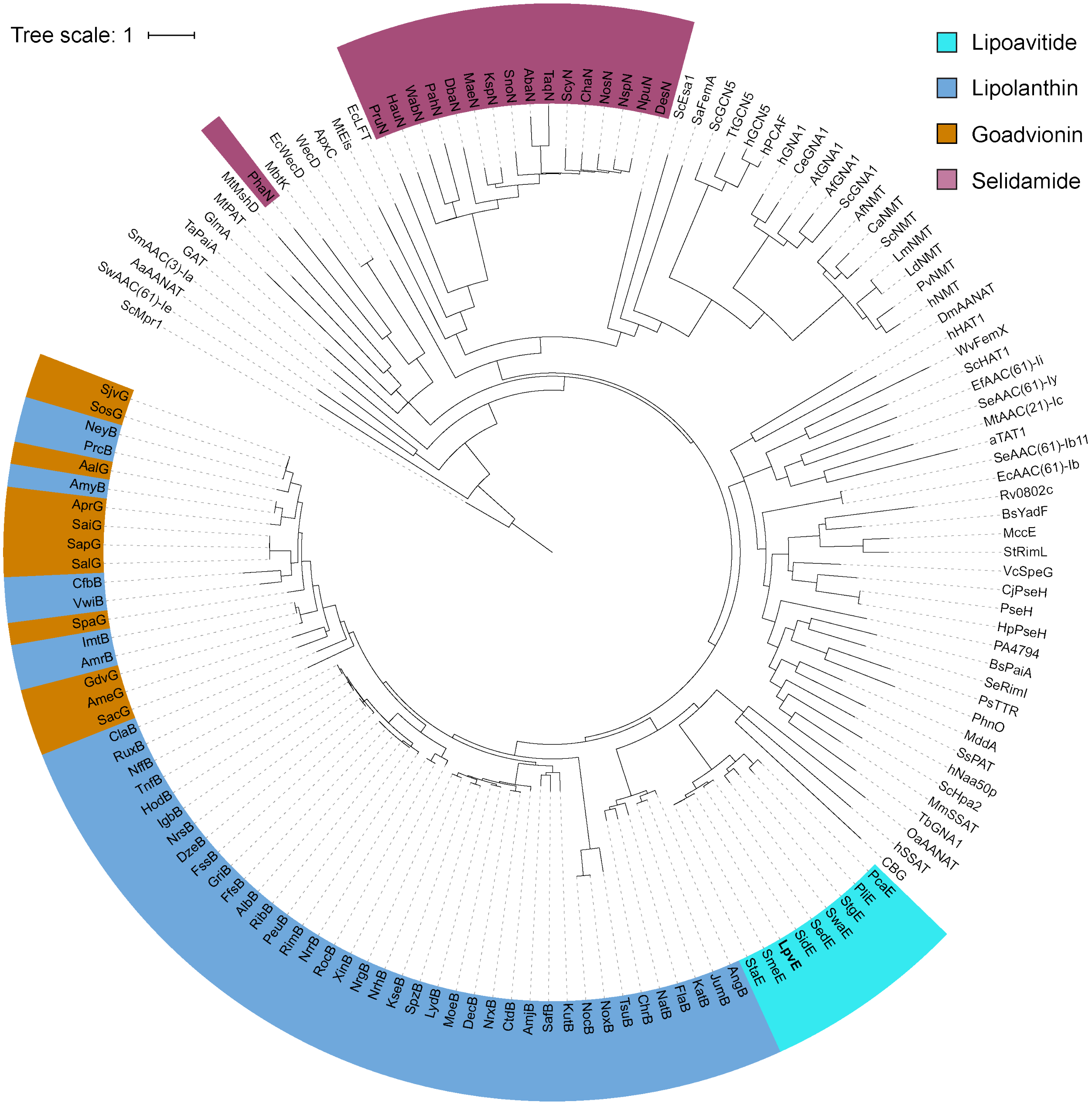

**Supplementary Fig. 50: Enzymatic reconstitution of StsG/LpvV/StsE**. (**a**) MALDI-TOF mass spectra of the reaction using **2** as the peptide substrate. The 3-HMP-modified **2** was observed at *m/z* 1636. (**b**) MALDI-TOF mass spectra of the reaction using **3** as the peptide substrate. The 3-HMP-modified **3** was observed at *m/z* 1476.

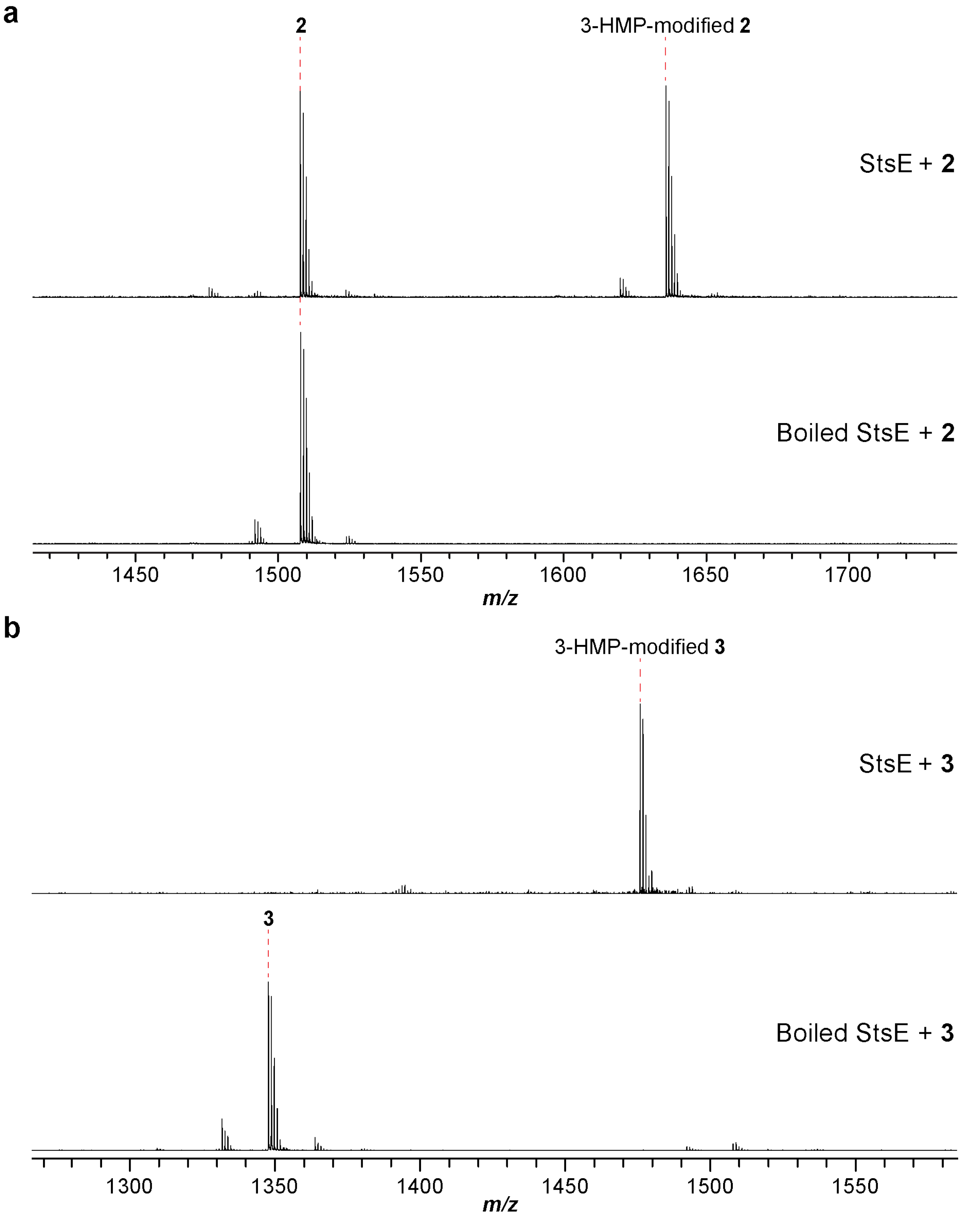

### References

1. Blodgett, J.A. et al. Unusual transformations in the biosynthesis of the antibiotic phosphinothricin tripeptide. *Nat Chem Biol* **3**, 480-485 (2007).
2. Tamura, K., Stecher, G. & Kumar, S. MEGA11: Molecular evolutionary genetics analysis version 11. *Mol Biol Evol* **38**, 3022-3027 (2021).
3. Enghiad, B. et al. Cas12a-assisted precise targeted cloning using in vivo Cre-lox recombination. *Nat Commun* **12**, 1171 (2021).
4. Kieser, T., Bibb, M.J., Buttner, M.J., Chater, K.F. & Hopwood, D.A. Practical streptomyces genetics (John Innes Foundation, 2000).
5. Fu, J. et al. Full-length RecE enhances linear-linear homologous recombination and facilitates direct cloning for bioprospecting. *Nat Biotechnol* **30**, 440-446 (2012).
6. Peter, D.M., Vogeli, B., Cortina, N.S. & Erb, T.J. A chemo-enzymatic road map to the synthesis of CoA esters. *Molecules* **21**, 517 (2016).
7. Stewart, J.J.P. MOPAC2016, Stewart Computational Chemistry, Colorado Springs, CO, USA.
8. Jumper, J. et al. Highly accurate protein structure prediction with AlphaFold. *Nature* **596**, 583-589 (2021).
9. Morris, G.M., Goodsell, D.S., Huey, R. & Olson, A.J. Distributed automated docking of flexible ligands to proteins: Parallel applications of AutoDock 2.4. *J Comput Aided Mol Des* **10**, 293-304 (1996).
10. Krieger, E. & Vriend, G. YASARA View-molecular graphics for all devices-from smartphones to workstations. *Bioinformatics* **30**, 2981-2982 (2014).
11. Trott, O. & Olson, A.J. Software news and update autodock vina: Improving the speed and accuracy of docking with a new scoring function, efficient optimization, and multithreading. *J Comput Chem* **31**, 455-461 (2010).
12. Wang, J.M., Cieplak, P. & Kollman, P.A. How well does a restrained electrostatic potential (RESP) model perform in calculating conformational energies of organic and biological molecules? *J Comput Chem* **21**, 1049-1074 (2000).
13. Duan, Y. et al. A point-charge force field for molecular mechanics simulations of proteins based on condensed-phase quantum mechanical calculations. *J Comput Chem* **24**, 1999-2012 (2003).
14. Wang, J.M., Wolf, R.M., Caldwell, J.W., Kollman, P.A. & Case, D.A. Development and testing of a general amber force field. *J Comput Chem* **25**, 1157-1174 (2004).
15. Jakalian, A., Jack, D.B. & Bayly, C.I. Fast, efficient generation of high-quality atomic charges. AM1-BCC model: II. Parameterization and validation. *J Comput Chem* **23**, 1623-1641 (2002).
16. Pettersen, E.F. et al. UCSF ChimeraX: Structure visualization for researchers, educators, and developers. *Protein Sci* **30**, 70-82 (2021).
17. Zimmermann, L. et al. A completely reimplemented MPI bioinformatics toolkit with a new hhpred server at its core. *J Mol Biol* **430**, 2237-2243 (2018).
18. van Kempen, M. et al. Foldseek: fast and accurate protein structure search. *Biorxiv*, 2022.2002. 2007.479398 (2022).
19. Kjaerulff, L. et al. Thioholgamides: Thioamide-containing cytotoxic RiPP natural products. *Acs Chem Biol* **12**, 2837-2841 (2017).
20. Frattaruolo, L., Lacret, R., Cappello, A.R. & Truman, A.W. A genomics-based approach identifies a thioviridamide-like compound with selective anticancer activity. *Acs Chem Biol* **12**, 2815-2822 (2017).
21. Jones, P. et al. InterProScan 5: Genome-scale protein function classification. *Bioinformatics* **30**, 1236-1240 (2014).
